# Human hippocampal ripples tune cortical responses in uncertain visual contexts

**DOI:** 10.1101/2023.08.30.555474

**Authors:** Darya Frank, Stephan Moratti, Johannes Sarnthein, Ningfei Li, Andreas Horn, Lukas Imbach, Lennart Stieglitz, Antonio Gil-Nagel, Rafael Toledano, Karl J. Friston, Bryan A. Strange

**Affiliations:** Laboratory for Clinical Neuroscience, Centre for Biomedical Technology, Universidad Politécnica de Madrid, Spain; Department of Experimental Psychology, Complutense University of Madrid, Spain; Department of Neurosurgery, University Hospital and University of Zurich, Switzerland; Neuroscience Center Zurich, University of Zurich and ETH Zurich, Switzerland; Movement Disorders and Neuromodulation Unit, Department of Neurology, Charité – Universitätsmedizin Berlin, corporate member of Freie Universität Berlin and Humboldt-Universität zu Berlin, Berlin, Germany; Department of Neurosurgery, Harvard Medical School, Massachusetts General Hospital, Boston, MA, United States; Brigham & Women’s Hospital, Center for Brain Circuit Therapeutics, Boston, MA, United States; Swiss Epilepsy Center, Klinik Lengg, Zurich, Switzerland; Epilepsy Unit, Department of Neurology, Hospital Ruber Internacional, Madrid, Spain; Wellcome Trust Centre for Neuroimaging, Institute of Neurology, University College London, London, UK; Reina Sofia Centre for Alzheimer’s Research, Madrid, Spain

## Abstract

To be able to encode information efficiently, our perceptual system should detect when situations are unpredictable (i.e., informative), and modulate brain dynamics to prepare for encoding. Here we show, with direct recordings from the human hippocampus and visual cortex, that after exposure to unpredictable visual stimulus streams, hippocampal ripple activity increases in frequency and duration prior to stimulus presentation, indicating context and experience-dependent prediction of predictability. Pre-stimulus hippocampal ripples suppress changes in visual (occipital) cortex gamma activity associated with uncertainty, and modulate post-stimulus prediction error gamma responses in higher-level visual (fusiform) cortex to surprising (i.e., unpredicted) stimuli. These results link hippocampal ripples with predictive coding accounts of neuronal message passing—and precision-weighted prediction errors—revealing a mechanism relevant for perceptual synthesis and subsequent memory encoding.

---

An efficient recognition system should be able to prioritise the information that will constrain or inform perceptual representations^1^ and, eventually, be accumulated or retained. One way to facilitate this is to continuously generate predictions about upcoming inputs and retain information—i.e., revise beliefs—when these predictions are violated. Prediction is considered central in this account of the brain, by building a generative model of the world to minimise prediction error when sampling the sensorium^2,3^. In predictive coding formulations, predictions are transmitted in a top-down manner, and when there is a mismatch between the predicted and the observed input, a bottom-up prediction error is returned to update or revise the source of predictions at a higher hierarchical level^4,5^. Crucially, prediction errors are weighted by the level of uncertainty (i.e., their precision) associated with the given context^6^, balancing top-down and bottom-up information streams to scale the influence of prior predictions and sensory evidence, respectively^7^. This is sometimes framed in terms of precision-weighted prediction errors that instantiate the Kalman gain in Bayesian filtering formulations of predictive coding^1,8^. The implicit encoding of uncertainty lends a dual aspect to predictive processing that encompasses both the predictions of a particular sensation and predictions of its predictability (i.e., precision) that modulate the influence of the ensuing prediction error^1,8–12^. Importantly, these generative properties of predictive processing rely on ongoing integration of sensory inputs with internally-generated, experience-dependent sequences, and are therefore thought to involve hippocampal-neocortical interactions^13–15^.

A hippocampal role in prediction is likely related to its function in extracting statistical regularities^16–19^, that can be applied to novel situations^20,21^. Therefore, the hippocampus should represent the expected information gain of an event before it occurs—as a function of predictability^15,19^—and estimate the validity of the prediction upon observation of the event^22^. Indeed, the hippocampus has long been postulated to hold a cognitive map that is used to form predictions about upcoming inputs^23–26^. Such predictive information might follow successor-like representation^27,28^, for example in place cell firing^29^. The cortex may also play its own predictive role through communication between deep layers feeding-back predictions to the superficial layers of preceding regions in the processing hierarchy^27,28^. Mechanistically, predictions and their violation have been associated with spiking activity ^30^ and oscillatory dynamics in the cortex^31,32^. Specifically, gamma-band activity has been associated with bottom-up prediction errors (reflecting surprise signals from primary sensory cortices), and alpha/beta oscillations with top-down predictions (from higher-level regions such as prefrontal cortex)^33–35^.

However, the mechanism through which predictions of predictability about upcoming sensory inputs are generated in the hippocampus and communicated to the cortex is still unclear^15^. Irrespective of these mechanisms, they should manifest in terms of a differential modulation of prediction error responses in visual cortex depending upon the predictability of the current context, which we hypothesise is itself recognised and broadcast via hippocampal processing. Specifically, when upcoming stimuli are unpredictable, they are inherently informative; in the sense they resolve uncertainty when observed (technically, they have a greater expected information gain). This leads to the hypothesis that the hippocampus plays a role in precision-weighting by modulating the electrophysiological correlates of prediction errors—i.e., event related gamma activity in the visual hierarchy—as a function of predictability (or entropy).

Hippocampal sharp-wave ripples (ripples henceforth) are found in every investigated mammalian brain and consist of sharp waves (large-amplitude, negative-polarity activity resulting from synchrony in the apical dendritic layer of CA1 pyramidal neurons) and ripples (∼140-200/s in rodents, resulting from interactions between excitatory and inhibitory neurons in CA3^36,37^). They are viewed as a pre-conscious mechanism to explore the organism’s options, searching for past experiences to extrapolate and predict future outcomes^25,38^. This is in addition to their role in replay of past experiences^39,40^. As prospection relies on past experiences, and is mediated by the hippocampus^24^, ripples may also underlie anticipation of future outcomes to guide subsequent behaviour. Indeed, ripples have been shown to portend behaviour in the immediate future, in the form of experienced trajectories^41^ and novel paths^42,43^, as well as pre-play of future events during sleep^44^. Specifically, the sequential firing pattern that occurs during rodent SWRs, in addition to ‘replay’ of spatial trajectories, has been shown to reflect all physically available trajectories within the environment not realised in prior behaviour ^45^, and trajectories taken by subjects in subsequent goal-directed navigation^46^. Furthermore, there is evidence that cortical activity is modulated in a peri-ripple manner, both through enhancement and inhibition of activity^47,48^, and as fluctuations in resting-state networks^49^. This is in line with predictive processing frameworks in which predictions and prediction errors are exchanged between levels in cortical hierarchies, with a special role for regions such as the hippocampus in contextualising this exchange^4,15^. Whilst ripples might represent possible outcomes, there is mixed evidence regarding whether they influence subsequent behaviour^50–52^. Importantly, the functional role of ripples is often examined using navigational tasks in rodents, which heavily tap episodic memory and are biased by the provision of reward at the goal location. In humans, although there is emerging evidence for ripples supporting memory recall processes^53–55^, it remains unclear to what extent they play a role in prediction, in the absence of memory demands.

We hypothesised that high-order predictions—namely, predictions of predictability or precision—would be generated by the hippocampus prior to stimulus presentation, as a function of uncertainty (i.e., predictability), and subsequently modulate cortical processing, or lower-level prediction errors. Ripples are a promising candidate to mediate the requisite precision-weighting of prediction errors. We tested this hypothesis using intracranial local field potential (LFP) recordings from human epilepsy patients, to examine how the hippocampus and ventral visual stream regions (occipital cortex and fusiform) implement predictive processing. We used a simple paradigm—void of any explicit demands on episodic memory—in which participants were presented a sequence of coloured shapes and performed a visuo-motor selection task, given a target stimulus and four options, presented on-screen simultaneously. The probability distribution of stimulus presentation varied across task blocks, allowing us to quantify stimulus-bound information-theoretic measures of entropy (i.e., uncertainty or unpredictability of an outcome before it occurs) and self-information (i.e., surprise or violation of predictions reflecting the improbability of a particular event) within each block and they can therefore be dissociated with respect to stimulus onset.

## Results

Fifteen participants successfully completed the task, with trial-by-trial measures of entropy and surprise modulating reaction time (RT) in accordance with Hick’s law^56^. Replicating results from this task in healthy adults^19^, participants’ RTs increased significantly per bit of surprise (t(14) = 10.26, p < 0.001, Cohen’s d = 2.64) and entropy (t(14) = 4.96, p < 0.001, Cohen’s d = 1.28; Figure 1b).

**Figure 1.**
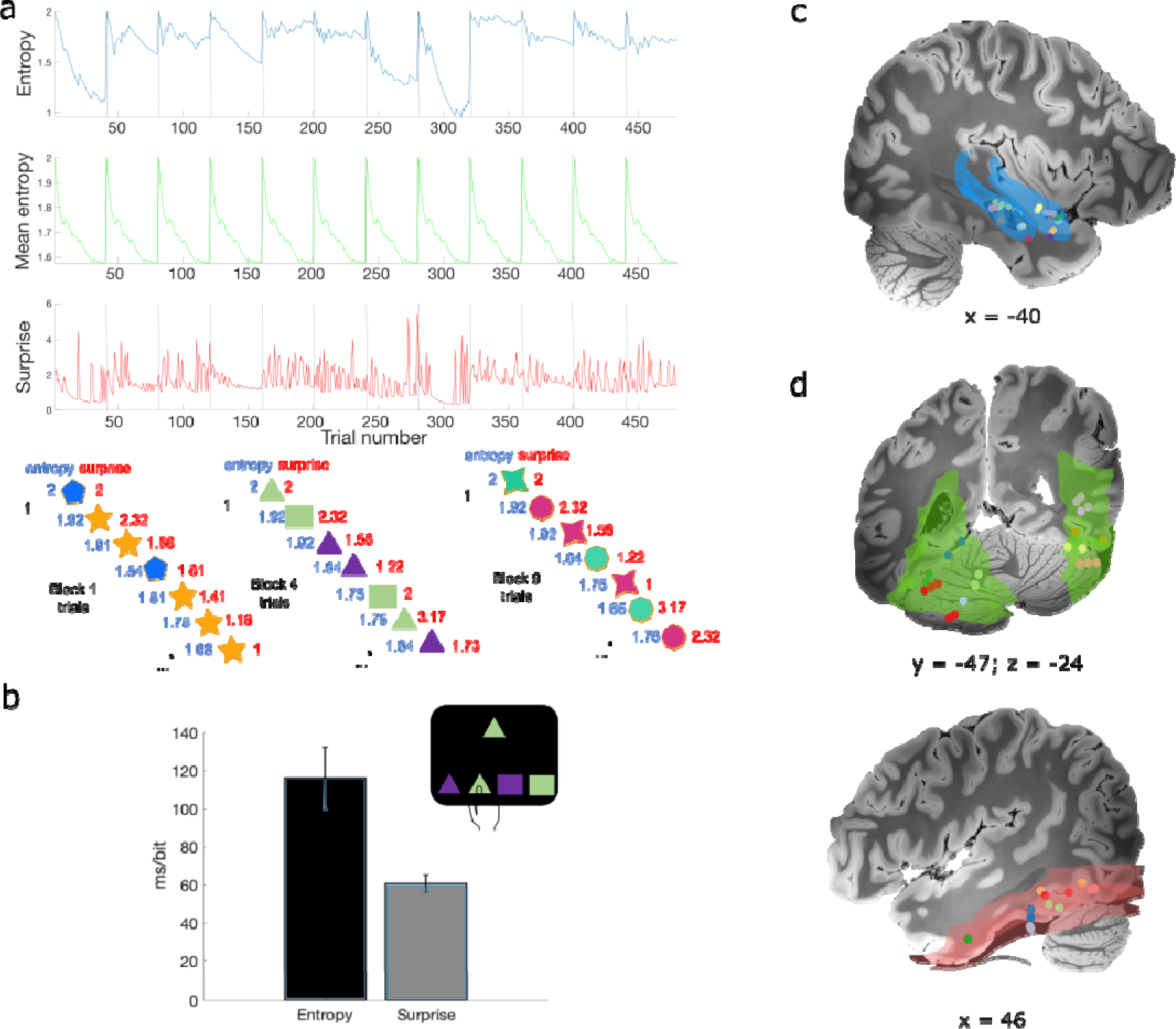
Task design and electrode contact localisation. **a)** Distribution of entropy, mean entropy and surprise values across trials for one example participant. Below, example trials from three of the 12 different blocks, and their associated entropy (blue) and surprise (red) values. Note that the 4 coloured shapes presented were unique to each block, with the probability of occurrence of these 4 stimuli varying between blocks. Participants were asked to perform a visuo-motor selection task, choosing the corresponding coloured shape from four alternatives with a button press, as depicted for block 4 in the current example. **b)** Group average reaction time as an increase in ms per bit of information-theoretic measure. **c-d)** Illustration of contacts in the hippocampus (smoothed hippocampus mask from the Automated Anatomical Labelling atlas for visualization; blue), fusiform (red), and occipital cortex (green) overlaid on a 100-μm T1 scan of an ex-vivo human brain, acquired on a 7T MRI scanner (https://openneuro.org/datasets/ds002179/versions/1.1.0); each colour represents a patient. See Figure S1 for individual patients’ contacts. Unless otherwise stated, error bars represent the standard error of the mean.

### Pre-stimulus ripple frequency and duration increase under uncertainty

In view of a possible generative role for hippocampal ripples in predicting the predictability of future events, we tested whether ripple occurrence was associated with increased uncertainty. Using a previously established ripple detection method^51^, a total of 2567 hippocampal ripples were detected in all participants while they performed the visuomotor task (Figure 2a). On examining the 2D distribution of ripple peak time as a function of peri-stimulus time and entropy (Figure 2b), we found a large proportion of ripples occurred in high entropy (1.7-1.9 bits) trials between −800 to −400ms pre-stimulus. There was also a peak in ripple probability just before the stimulus onset (−200 to 0ms) for entropy values from 1.6-1.7 and 1.8-1.9 bits. A 2-samples Kolmogorov-Smirnov, showed a significant difference between the ripple distribution and a uniform distribution with the same minimum and maximum counts (k = 0.183, p = 0.038). We also performed a Spearman’s correlation between the observed and permuted distributions to ensure ripple distribution was not correlated with random noise. The correlation between the two was computed in every permutation (1000 permutations in total) per participant, and the mean correlation across participants was compared to 0 using a one-sample t-test. The observed 2D distribution was not correlated with the random permutations (t(14) = 0.629, p = 0.53). We have also performed a mixed-effects GLM on the normalized count values to identify the bins showing the largest number of ripples. We found a significant interaction between entropy and ripple peak time (χ^2^(99) = 200.1, p < 0.001), with the five largest estimated marginal means identified around −1000 to −400ms and 1.6-1.8 entropy bins, as well as just before stimulus onset (−0.2 – 0, entropy values 1.8-1.9) and just after stimulus onset (0 – 0.2, entropy values 1.8-1.9). This is in line with the visual representation of the 2D distribution shown in Figure 2c (all estimated marginal mean values shown in Supplementary Figure S3). Given that we do not see many ripples after 600ms post-stimulus, the pre-stimulus ripples identified are unlikely to reflect post-processing of the previous stimulus (see also Figure S10 showing ripple distribution as a function of surprise from 0-2.2s).

**Figure 2.**
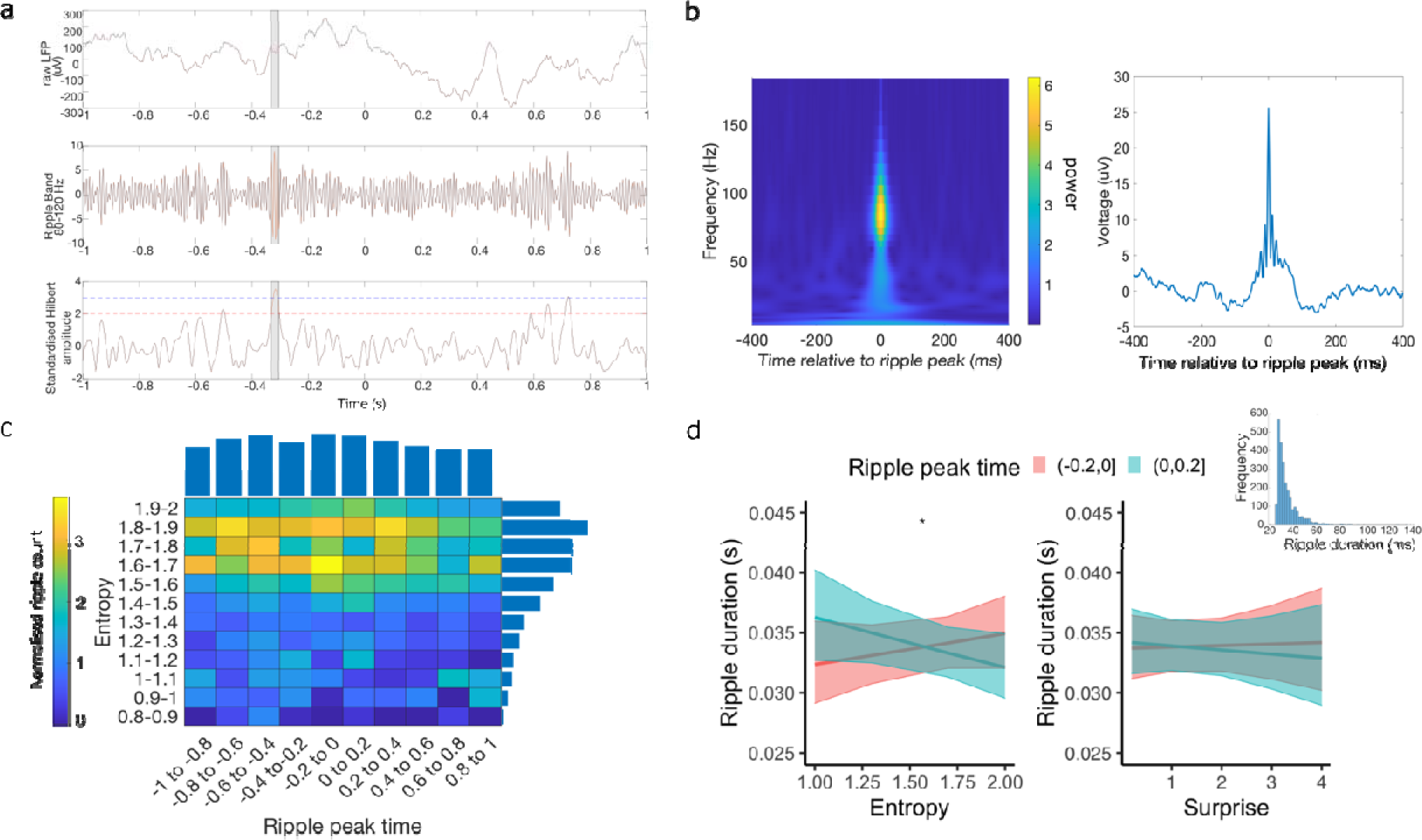
The frequency and duration of hippocampal ripples increase with uncertainty or expected information gain. **a) an example ripple detected using the Vaz et al (2019) method, showing the raw LFP trace (top), the ripple band signal (middle) and the standardised envelope (bottom) b)** Grand average peri-ripple wavelet spectrogram (left) and raw field potential centred on ripple peak (n = 2567 ripple events from 15 participants). **c)** Normalized ripple distribution across peri-stimulus time and entropy levels, accounting for the total number of trials in each entropy bin. Normalized ripple count for each time and entropy bin is colour-coded; average counts for each time and entropy bin are plotted as bars above and to the right of the 2D distribution, respectively. The majority of ripples occurred pre-stimulus and in high (but not highest) entropy levels (see Figure S2c for peri-stimulus ripple distribution). **d)** Ripple duration as a function of peri-stimulus time and information-theoretic measures. Ripple duration increased with entropy (i.e., expected information gain for pre-stimulus ripples, but decreased for post-stimulus ripples). Ripple duration histogram, over all recorded ripples, in shown above right. Shaded areas represent the 95% confidence interval. See Extended Data for replication using a different ripple detection method^53^.

The duration of ripples has been shown to be functionally relevant for memory consolidation in rodents^57^. In predictive coding formulations, this rests upon the augmentation of associative (i.e., activity or experience-dependent) plasticity by precision-weighting that increases presynaptic (prediction error) afferents^58,59^. To test for changes in ripple duration with uncertainty, we next examined the relationship between ripple duration and the information-theoretic measures. As ripple duration followed a right-skewed distribution (Figure 2d), a mixed-effects Gamma regression was employed. First, we tested whether ripple duration was modulated by peri-stimulus time, entropy, and surprise. We did not observe any significant main effects (all p’s > 0.219) or interactions (time window by entropy χ^2^(1) = 0.876, p = 0.349; time window by surprise χ^2^(1) = 0.264, p = 0.607; see Figure S2g). However, comparing ripples occurring just before (−200 to 0ms; where there was an increase in ripple frequency; Figure 2b) and the corresponding period just after the stimulus (0 to 200ms), revealed a significant main effect of peri-stimulus time window (χ^2^(1) = 5.08, p = 0.0241), as well as an interaction between time window and entropy (χ^2^(1) = 3.97, p = 0.0463, Figure 2d). This interaction suggests that as entropy increased, pre-stimulus ripple duration increased, but decreased for post-stimulus ripples. Ripple duration was again not modulated by surprise (main effect χ^2^(1) = 0.03, p = 0.862; interaction χ^2^(1) = 2.15, p = 0.643), suggesting ripple duration reports predictability (i.e., expected information gain) as opposed to violations of predictions (i.e., observed information gain).

### Reaction time as a function of pre-stimulus ripples

Given that there is a relationship between entropy and RT^19,56^, as well as between ripples and entropy, we examined whether the presence of ripples modulates the relationship between RT and entropy**—** and surprise**—** on a trial-by-trial basis. To ensure sufficient pre-stimulus ripple trials were included, this analysis was restricted to trials with 1.6 to 1.9 bits of entropy, where most pre-stimulus ripples were observed (50% of pre-stimulus ripple trials and 50% of no ripple trials are within these values). We found a significant main effects of ripple status, with faster responses in trials with a pre-stimulus ripple (χ^2^(1) = 4.33, p = 0.037). This is an important observation because the precision of prediction errors corresponds to the rate of evidence accumulation that underwrites reaction speed. In other words, prediction errors that are afforded more precision exert their effects on belief updating more rapidly. There was also a main effect of surprise, with faster RTs for lower levels of surprise (χ^2^(1) = 83.3, p < 0.001), as per Hick’s law. The interaction between ripple status and surprise, however, was not significant (χ^2^(1) = 2.24, p = 0.134) indicating that ripples did not modulate the relationship between RT and surprise. There was a trend towards an interaction between entropy and ripple status (χ^2^(1) = 3.46, p = 0.0628; Figure S2e), with simple slopes of entropy in trials without ripples (estimate = 0.11, t(14) = 1.29, p = 0.2) and in trials with pre-stimulus ripples (estimate = 0.37, t(14) = 3.19, p = 0.001) indicating a steeper positive slope in trials with pre-stimulus ripples.

### Hippocampal and occipital cortex pre-stimulus gamma activity track uncertainty in opposing ways

To further characterise hippocampal and cortical correlates of entropy and surprise, we next examined time-resolved spectral responses as a function of these measures. The focus was on two cortical regions: the fusiform gyrus, previously shown with functional MRI to respond to surprise in this task^19^, and occipital cortex, lower down in the visual cortical hierarchy than the fusiform and putatively supplying it with bottom-up information (e.g., prediction errors). Hippocampal activity in the gamma range (47.5-97.5 Hz; Figure 3a) was negatively associated with entropy in the pre-stimulus period, around −1000ms to −660ms pre stimulus (summed cluster t value = −1250.4, p = 0.0074; smallest t value in cluster = −6.23, p = 0.0058, Cohen’s d = 1.6), showing reduced gamma power preceding more uncertain outcomes. This time window partly overlaps with peaks in ripple occurrence, although, notably, the gamma power around this time-window was reduced under high entropy. Further examination of the negative association with entropy, as a function of trial in block, showed that the reduction in gamma power was mostly concentrated in the first few trials of the block (Figure S4a), prior to the emergence of pre-stimulus ripples.

**Figure 3.**
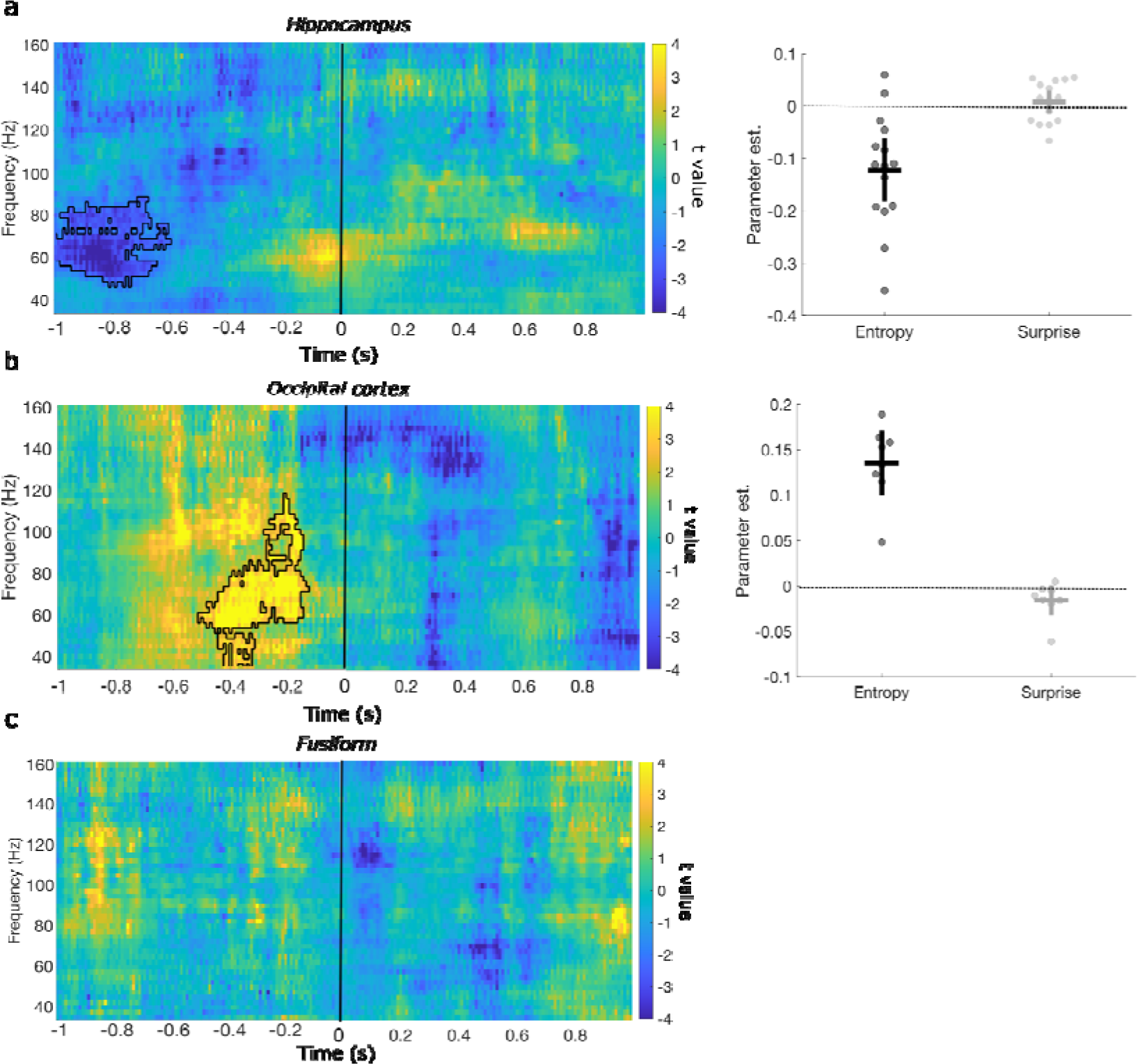
Hippocampal and occipital cortical gamma activity is modulated by entropy *before* stimuli are presented. **a)** Left. Time-resolved spectral power in the hippocampus shows a significant negative association between gamma-power and entropy around 800ms pre-stimulus (significant cluster outlined in black here and in subsequent figures). During highest entropy levels, gamma power was lowest (see Figure S3a for gamma power as a function of trial in block). No significant clusters were found in the post-stimulus time window. Right. Parameter estimates for each predictor within the significant cluster; each point pertains to one patient, horizontal bars the group mean, and vertical error bars represent 95% confidence interval. **b**) In occipital cortex there was an increase gamma power as entropy increased, around 400ms pre-stimulus. Again, entropy was not associated with changes in the post-stimulus time windows. **c)** In the fusiform cortex, no effects of entropy were observed either pre- or post-stimulus.

In occipital cortex, on the other hand, a positive association between gamma activity (35-117.5 Hz; Figure 3c) and entropy was observed around −510ms to −130ms pre-stimulus (summed t value = 1694.1, p = 0.0078, largest t value in cluster = 10.67, p = 0.0078, Cohen’s d = 3.77). No significant responses to entropy were found in the fusiform, or in the post-stimulus time period for all three areas (Figure 3). Under the simplifying assumption that gamma activity reflects the amplitude of precision-weighted prediction errors, these results are consistent with an increase in the precision of visual prediction errors, with a concomitant decrease in the precision of hippocampal prediction errors, prior to stimuli with higher expected information gain. This could be read as instantiating the right kind of attentional set, when salient information can be anticipated in advance^60,61^.

### Ripple-triggered cortical responses

Next, we investigated whether ripple occurrence instates a temporal relationship between the hippocampus and visual processing regions in which pre-stimulus modulations of gamma activity were also observed as a function of entropy. First, we compared peri-ripple cortical time-frequency responses, dichotomised by whether the hippocampal ripple occurred before or after stimulus onset (Figure 4a-b). We found an overall increase in high gamma power in the fusiform (−200 to 200ms peri-ripple, 105-160Hz, summed cluster t value = 2171.1, p = 0.0156; largest t value in cluster = 4.44, p = 0.0156, Cohen’s d = 1.67) as well as in occipital cortex (−180 to 200ms peri-ripple, 95-160Hz, summed cluster t value = 1474, p = 0.0039; largest t value in cluster = 9.15, p = 0.0039, Cohen’s d = 3.27) locked to hippocampal ripples. Furthermore, in occipital cortex, pre- and post-stimulus ripples modulated activity differently (−10 to 200ms peri-ripple, 35-160Hz, Figure 4c; summed t value = −1094.5, p = 0.0039; smallest t value in cluster = −8.99, p = 0.0039, Cohen’s d = 3.17). Specifically, occipital gamma activity was reduced following pre-stimulus ripples compared to post-stimulus ripples. This effect was partly driven by a power suppression, compared to a baseline period, of occipital gamma around pre-stimulus ripples (Figure 4d shows raw power suppression in relation to baseline). That is, when examining occipital gamma activity in the pre-stimulus period in which a positive association with entropy was found (indicating an overall power increase), there was reduced gamma power in trials with a hippocampal ripple in the period before the significant cluster compared to trials with no ripples (t(7) = 2.51, p = 0.040, Cohen’s d = 0.889; Figure S7a). Taken together, these findings are indicative of a hippocampal pre-stimulus ripple-induced suppression of pre-stimulus gamma power in the occipital cortex. No differences were observed between pre- and post-stimulus ripples modulation of fusiform activity, or in lower frequencies in either cortical region.

**Figure 4.**
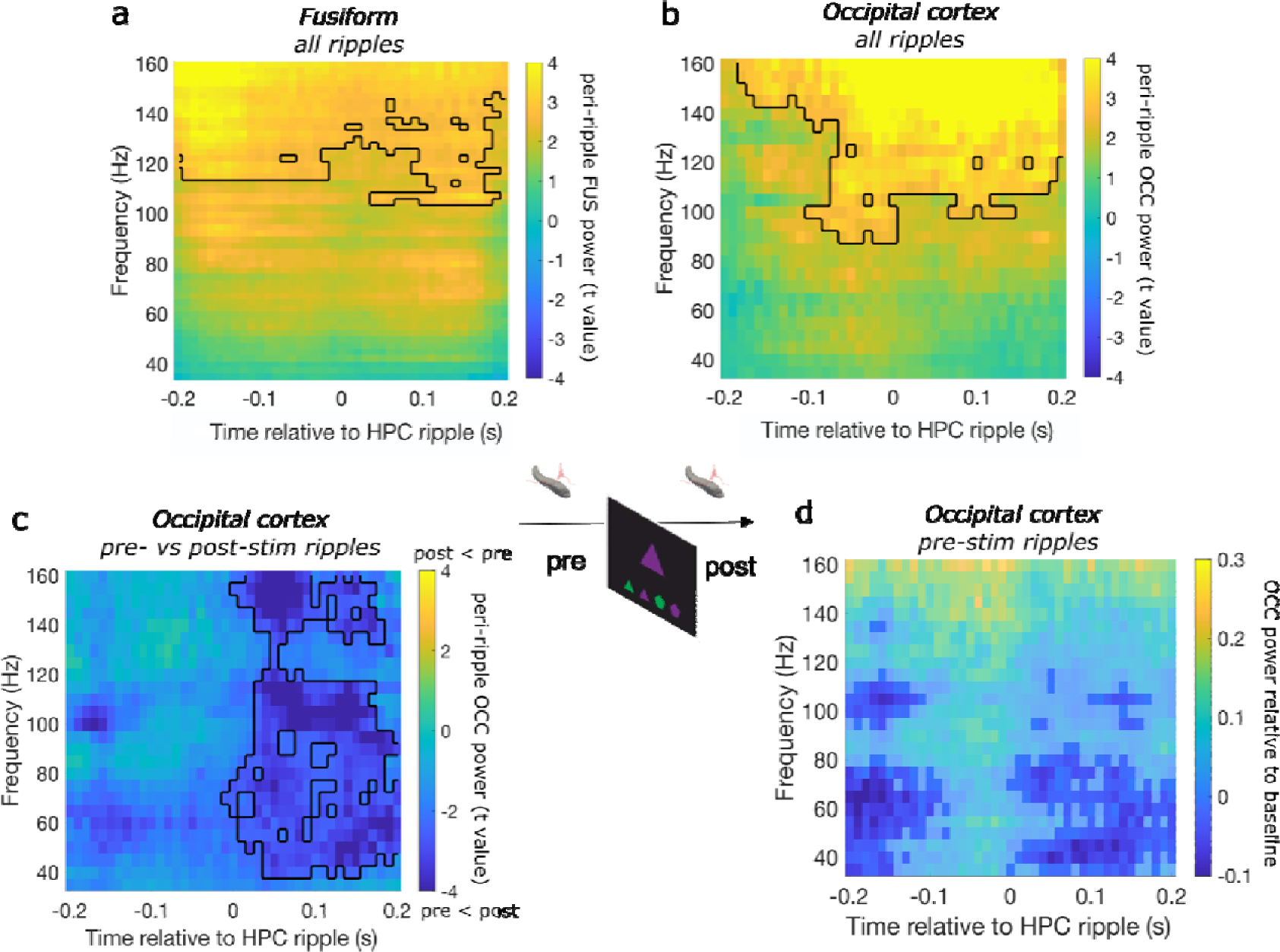
Peri-hippocampal ripple cortical activity. Cortical time-frequency analyses time-locked to the hippocampal ripple peak time. Across all ripple trials there was an overall increase in high gamma power in the fusiform (**a**) and occipital cortex (**b**) time-locked to hippocampal ripple**. c)** Comparing trials with pre-versus post-stimulus hippocampal ripples, there was a reduction in occipital gamma power in trials with pre-stimulus ripples, compared to post-stimulus ripples. Further examination of this effect revealed it was driven by a suppression of gamma power relative to baseline in trials with pre-stimulus hippocampal ripples (**d**) and compared to trials with no ripples (Supplementary Figure 4a). See Extended Data for replication using a different ripple detection method

Next, we examined whether ripple-modulated cortical activity was correlated with entropy or surprise. There were no significant associations between peri-ripple cortical activity and either information-theoretic measure, suggesting that the hippocampal ripple modulation of cortical response was not affected by uncertainty or surprise, and thus ripples may act as a general mechanism for hippocampal modulation of cortical activity, with context-dependence of this modulation effected by ripple rate or duration.

### Hippocampal and fusiform – but not occipital – cortex gamma responses to surprise

We expected that prediction error, in the form of surprise, would elicit hippocampal and cortical responses. In line with previous findings, a positive association between surprise and hippocampal activity in the gamma range (55-80 Hz; Figure 5a) was observed around 200 to 540ms post-stimulus (summed cluster t value = 638.5, p = 0.0248; largest t value in cluster = 4.89, p = 0.0248, Cohen’s d = 1.2). In the fusiform, there was a positive association between surprise and gamma activity (40-95 Hz; Figure 5b) around 200-510 post-stimulus (summed cluster t value = 741.9, p = 0.0234, largest t value in cluster = 6.79, p = 0.0234, Cohens’ d = 2.56), and a negative association between surprise and alpha/beta power (10-15 Hz; Figure 5c) around 510-840ms post-stimulus (summed cluster t value = −199.5, p < 0.001; largest negative t value in cluster = −5.41, p < 0.001, Cohen’s d = 2.05). No significant associations between surprise and occipital activity were observed.

**Figure 5.**
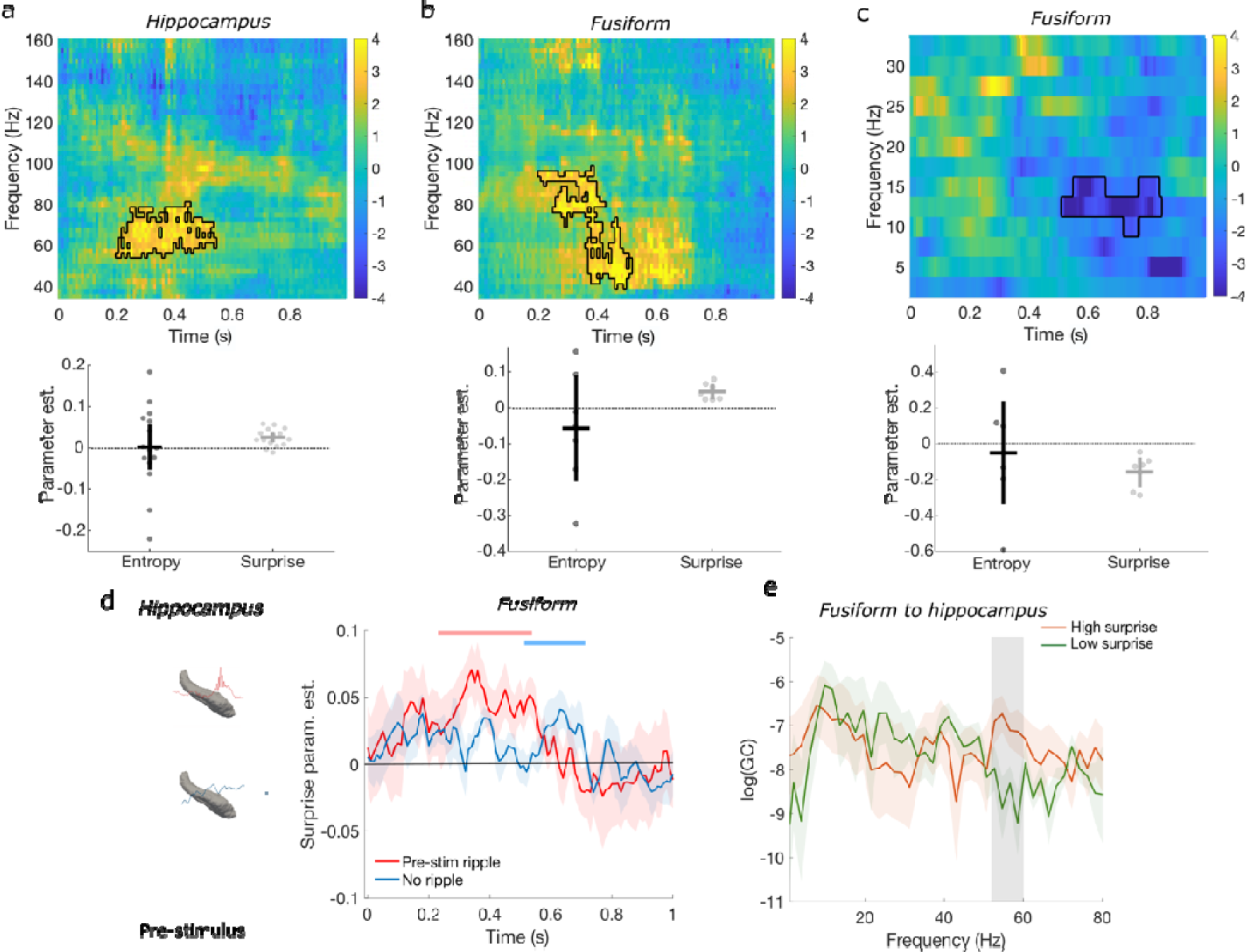
Surprise driven responses. **a)** Hippocampal increase in gamma power as surprise increases. Below: parameter estimates for each predictor within the significant cluster. **b)** Fusiform gamma power increase as surprise increases. **c)** Fusiform beta oscillations decrease as surprise increases. **d)** The increase in fusiform gamma power 40-95 Hz (extracted from significant cluster) occurs earlier in trials with pre-stimulus hippocampal ripples (red) compared to trials without ripples (blue). See Extended Data for replication using a different ripple detection method. **e)** Directed information flow from fusiform to hippocampus for high (orange) versus low (green) surprise trials was observed in the gamma band (shaded rectangle).

### Pre-stimulus hippocampal ripples modulate prediction error

Under the predictive processing framework, a cardinal function of neural activity is to minimise precision-weighted prediction error^62^. A possible role for hippocampal ripples is to broadcast the predicted precision of forthcoming prediction errors. Specifically, we hypothesised that pre-stimulus ripples would amplify precision-weighted prediction errors, evoked by stimuli in an unpredictable (high entropy) context. To test this hypothesis, we examined whether the occurrence of pre-stimulus hippocampal ripples modulated post-stimulus (0-1s) cortical responses to surprise (e.g. the increase in gamma power in the fusiform). This hypothesis was confirmed in the fusiform cortex. We found an early (250-550ms post stimulus, summed t value = 936.9, p = 0.0078) increase in fusiform gamma power (47.5-122.5Hz) in response to higher surprise in trials that were preceded by a pre-stimulus hippocampal ripple. By contrast, a later (510-710ms) increase in gamma power (50-72.5 Hz, summed t value = 473.8, p = 0.0078) was observed to higher surprise when there was no pre-stimulus ripple (Figure 5d and Figure S9a-b). The presence of a faster fusiform response to surprise in trials with pre-stimulus ripples could simply reflect a co-occurrence of the two. To rule out this possibility, we examined whether there was a double dissociation of the surprise parameter estimate using a repeated measures ANOVA with ripple status (no ripple, pre-stimulus ripple) and time window (early, later) as factors. This showed a significant interaction (F(1,6) = 19.6, p = 0.004, η²_p_ = 0.766), confirming the double dissociation (See Figure S9c) such that an early effect of surprise was observed for trials with pre-stimulus ripples, but not for trials without ripples; conversely, a later surprise effect was found in trials without ripples, but not in trials with pre-stimulus ripples. This result suggests that the presence of a pre-stimulus hippocampal ripple modulates the fusiform response to surprise, such that it is faster—and of greater amplitude—compared to trials without a pre-stimulus ripple. The presence of pre-stimulus hippocampal ripples did not modulate post-stimulus occipital responses to surprise.

### Directed information flow during prediction and prediction errors

Lastly, we examined information flow between the hippocampus and fusiform and occipital cortex by testing for Granger causality as a function of surprise and entropy. To do so, a median-split for each information-theoretic measure was applied and a non-parametric spectral Granger causality analysis performed. Guided by the time-frequency effects above, we focused on hippocampus-occipital cortex for entropy, and hippocampus-fusiform for surprise, each centred in time around the observed time-frequency effect (−1000 to −500ms pre-stimulus for entropy, and 150 to 650ms post-stimulus for surprise). For surprise, we found increased bottom-up information flow (fusiform ➔ hippocampus) in the gamma range (52-60 Hz; Figure 5e) for high versus low surprise (summed t value = 13.2, p = 0.0078). To ensure this effect was valid we ran two further analyses; a Granger causality analysis on the time-reversed data, showing an effect in the same frequency ranges in the opposite direction, as expected. And secondly, a partial directed coherence analysis which revealed a significant information flow effect from the fusiform to the hippocampus in high versus low surprise trials for the same frequency range 52-59Hz (cluster t(6) = 4, p < 0.001; Figure S9g). No significant effects were observed in the reverse direction (hippocampus ➔ fusiform) or between the hippocampus and occipital cortex for entropy (Figure S9d-f). As there were few participants with contacts in both cortical regions (N = 4), Granger causality between occipital and fusiform cortex was not examined.

## Discussion

We measured hippocampal and cortical activity using depth electrodes in human epilepsy patients as they observed sequences of simple visual stimuli drawn from a block-specific distribution, yielding different levels of stimulus-bound uncertainty and surprise. Electrophysiological intracranial activity was examined pre- and post-stimulus to assess prediction under different levels of uncertainty, and subsequent validation against observed inputs, respectively. Our results highlight an important hippocampal function in generating putative predictions and communicating them downstream to the cortex. Specifically, our findings suggest hippocampal ripples are key in this generative process; ripple probability was highest in the pre-stimulus window when uncertainty was high, but not at its highest levels, and pre-stimulus ripple duration increased with entropy. These pre-stimulus ripples were associated with subsequent faster responses to post-stimulus surprise in the fusiform cortex. This is exactly consistent with a role of hippocampal ripples reporting the predictability of an upcoming stimulus and thereby affording the ensuing prediction errors greater precision, in the context of greater expected information gain (i.e., higher entropy).

The properties and distribution of ripples as a function of peri-stimulus time and entropy strongly suggest that they play an important role in generating a certain kind of prediction; namely, a prediction of predictability. Previous work in rodents has demonstrated that ripples represent future trajectories of experienced and novel paths^41–43^, indicative of a generative process. Our results provide support for, and extend, these previous studies by showing increased ripple probability in uncertain contexts before stimulus onset, in awake human patients, as they anticipate upcoming sensory input. Specifically, predictive processing under uncertainty may represent an integral part of the proposed role for SWRs in planning future actions^63,64^. Our findings are in keeping with a recent synthesis of rodent studies suggesting that SWR trajectories comprise not only the path to be chosen or the path just completed, but many of the potential options available^65^. With an increasing number of options—and implicit unpredictability of ensuing outcomes—the ripple rate would be expected to increase, as observed here. This is also in line with recent fMRI work in humans showing successor-like representations in the hippocampus and visual cortex^28^. Importantly, our visuo-motor mapping task did not impose memory demands, as both stimulus and response mapping appeared on the screen simultaneously. The key manipulation was the underlying probability of the stimulus distribution within-block, which we argue elicited an increased need for precise predictive processing, and thus more ripple events, with increased uncertainty regarding the upcoming stimulus. This allowed us to elucidate an important functional role that ripples play in anticipation of sensory inputs and associated action planning^66^.

The increased frequency of pre-stimulus ripples occurred around the same time as the negative association between entropy and gamma power (overlapping in 80-97.5 Hz), namely, around 800ms pre-stimulus. The early hippocampal gamma effect (i.e., negative association with entropy) is—on a predictive coding narrative—consistent with an attenuation of neuronal fluctuations in populations reporting prediction errors (e.g., superficial pyramidal cells) in proportion to the predicted precision of subsequent (stimulus-bound) prediction errors^12^. Such an early effect is consistent with (Bayesian belief) updating of information from the previous trial and broadcasting the ensuing predictions down the cortical hierarchy. This finding also suggests that detected ripples were likely discrete events, dissociable from gamma activity^12,37,67^. The observed reduction in gamma activity (spanning frequencies lower than that of the ripple activity) occurred in the pre-stimulus period on trials in which pre-stimulus ripples were evident (Figure S5). This co-occurrence might be driven by the opposing effects of acetylcholine on ripple generation and gamma power (increase in ripple rate accompanied by decreased gamma-band activity, and *vice versa*^37^). This is interesting because acetylcholine has been implicated in the encoding of precision in predictive processing in several computational studies^3,15–18^.

To demonstrate that ripples carry predictive information that is subsequently propagated to the cortex, it is important to establish peri-ripple modulation of behaviour or cortical activity, as suggested by predictive processing^4,15^. There are mixed findings in the literature, with some studies showing no ripple occurrence-related modulation of subsequent behaviour in rodents^50–52^. We found limited support for ripple modulation of behaviour, with faster responses at high levels of entropy (1.6-1.8) when there were pre-stimulus ripples, but the negative correlation between pre-stimulus ripple rate and decrease in RT (as ms/bit of entropy) did not reach statistical significance (see Figure S2e). Nevertheless, hippocampal ripples did modulate cortical activity in several ways; in line with previous findings from rodents and non-human primates^47–49^, we found an overall increase in high gamma power in fusiform and occipital cortex around the ripple time, supporting the notion that occurrence of ripples facilitates enhanced interaction between the hippocampus and cortex.

In addition to the cortical modulation across all ripple trials, a unique modulation of pre-stimulus ripples on cortical processing was observed, in keeping with their predictive role. First, we found a suppression of occipital gamma activity that was time-locked to the hippocampal ripple event. Together with the overall positive association between entropy and occipital gamma power pre-stimulus, this suggests that the ripple-triggered suppression may reflect an override hippocampal signal over lower-level cortical prediction of the upcoming stimulus, or facilitate a sharper representation of the possible outcomes in occipital cortex^13,68^. Second, the presence of pre-stimulus ripples modulated post-stimulus fusiform activity; whilst there was an overall positive association between trial-wise surprise and gamma power in fusiform around 300ms post-stimulus, when splitting trials based on the presence of pre-stimulus hippocampal ripples, we found a faster fusiform gamma response to surprise, compared to trials without pre-stimulus ripples. Gamma-band activity has been shown to support bottom-up processing^34,35^, so that the faster response to surprise in the presence of pre-stimulus ripples might indicate that a prior prediction facilitates signalling of prediction errors^30^, and their propagation up the visual processing hierarchy. Further support for this claim comes from the subsequent increased fusiform to hippocampus information flow in high surprise as reflected in Granger causality for gamma activity from 150 to 650 post-stimulus, which we associate with bottom-up prediction error signalling. This is compatible with a view that pre-stimulus hippocampal ripples modulate (or prime) the fusiform to respond at earlier latencies to surprising stimuli (top-down effect), and the fusiform post-stimulus gamma response to surprise feeds back to the hippocampus (bottom-up effect). The nature of hippocampal ripple-triggered changes in cortical state remains to be determined, although the simplicity of the current behavioural task lends itself to further mechanistic interrogation using non-human animal models, where homologous predictive coding mechanisms are evident^69,70^.

Finally, we also observed post-stimulus hippocampal ripples, but did not, however, find a relationship between these and behavioural measures or cortical activity. Different cortical effects linked to pre-versus post-stimulus hippocampal ripples could imply that these ripples arise in different neuronal populations whose outputs are routed differently. Therefore, it is possible that post-stimulus ripples serve a different functional role^40,71^, or facilitate communication with other brain regions, such as the prefrontal cortex^47,48^.

In conclusion, our findings speak to an important function of hippocampal ripples in predictive processing, reporting the predictability or expected information gain of stimuli (i.e., prediction errors) before they are encountered; thereby facilitating their propagation through the visual cortex (in the absence of any memory demands). More specifically, our results reveal an increase in ripple events in uncertain trials, prior to stimulus onset, and subsequent modulation of cortical activity, pointing to enhanced hippocampal-cortical communication facilitated by ripples that serves propagation of precise prediction errors.

## Materials and Methods

### Participants

Sixteen medication-resistant presurgical epilepsy patients (mean age = 38.9, SD = 11.5; 7 males) with depth electrodes surgically implanted to aid seizure focus localization took part in the experiment. Data were acquired at two epilepsy centres. All patients signed informed consent prior to participating. Implantation sites were chosen solely on the basis of clinical criteria. Patients had normal or corrected-to-normal vision and had no history of head trauma or encephalitis. All patients had electrodes implanted in the hippocampus. For patients showing unilateral hippocampal sclerosis, only the non-pathological side was included in the analysis, otherwise all hippocampi were radiologically normal on pre-operative MRI.

Of the 16 patients who completed the task, one patient was excluded from any analysis due to poor performance on the task (less than 25% accuracy on all trials). We therefore analysed electrophysiological responses from 15 patients. In total, we analysed contacts in the right hippocampus from seven patients, and eight patients with contacts on the left side. Of these 15 patients, eight patients also had electrodes in the occipital cortex, and seven patients also had contacts in the fusiform. Our sample size was not determined prior to the experiment, but it is equal or larger than that reported in previous publications using cognitive tasks with iEEG ^53,72^. This study has been approved by the local ethics committees of Hospital Ruber Internacional, Madrid, Spain and Kantonale Ethikkommission, Zurich, Switzerland (PB-2016-02055).

### Stereotactic electrode implantation

A contrast-enhanced MRI was performed pre-operatively under stereotactic conditions to map vascular structures prior to electrode implantation, and to calculate stereotactic coordinates for trajectories using the Neuroplan system (Integra Radionics). For patients whose data was collected in Madrid (N = 12), DIXI Medical Microdeep depth electrodes (multi-contact, semi rigid, diameter of 0.8 mm, contact length of 2 mm, inter-contact isolator length of 1.5 mm) were implanted based on the stereotactic Leksell method. For patients whose data was collected in Zurich (N = 3), the depth electrodes (1.3 mm diameter, 8 contacts of 1.6 mm length, and spacing between contact centres 5 mm; Ad-Tech, Racine, WI, www.adtechmedical.com) were stereotactically implanted in the medial temporal lobes (MTL).

### Data acquisition

In Madrid, intracranial EEG (iEEG) activity was acquired using an XLTEK EMU128FS amplifier (XLTEK, Oakville, Ontario, Canada). iEEG data were recorded at each electrode contact site at a 500Hz sampling rate (online bandpass filter 0.1–150Hz) and referenced to linked mastoid electrodes. For four patients, the data were recorded with a higher sampling rate, but were later down-sampled to 500Hz. In Zurich, data were acquired using a Neuralynx ATLAS system with a sampling rate of 4000Hz (online band-pass filter of 0.5–1000Hz) against a common intracranial reference, and then down-sampled to 500Hz. Data from the two centres were comparable, and we did not observe differences between centres in any of the analysis reported in the main text.

### Electrode localization – hippocampus

To localize electrodes with contacts in the hippocampus, we used the manual procedure described previously ^72^. For each patient, the post-electrode placement CT (post-CT) was co-registered to the pre-operative T1-weighted MRI (pre-MRI). To optimize co-registration, both brain images were first skull-stripped. For CTs this was done by filtering out all voxels with signal intensities between 100 and 1300 HU. Skull stripping of the pre-MRI proceeded by first spatially normalizing the image to MNI space employing the New Segment algorithm in SPM8 (http://www.fil.ion.ucl.ac.uk/spm). The resultant inverse normalization parameters were then applied to the brain mask from SPM8, to transform the brain mask into the native space of the pre-MRI. All voxels in the pre-MRI that were outside the brain mask and with a signal value in the top 15% were filtered out. The skull-stripped pre-MRI was then co-registered and re-sliced to the skull-stripped post-CT. Next, the pre-MRI was affine registered to the post-CT, thus transforming the pre-MRI image into native post-CT space. The two images were then overlaid, with the post-CT thresholded such that only electrode contacts were visible. For all patients, only contacts in the hippocampus head and body were selected. For patients with multiple electrodes localised in the hippocampus, only the most anterior contacts were used in the analyses (see Figure 1c). Electrode contacts for each patient are shown in Figure S1.

### Electrode localization – cortical regions

To localize electrodes with contacts in the occipital cortex and fusiform, we used a semi-automatic procedure utilizing Lead-DBS^65^ (lead-dbs.org). First, the post-operative CT was registered to the pre-operative MRI using a two-stage linear registration (rigid followed by affine) as implemented in Advanced Normlization Tools^73^. The images were then normalized to the MNI template based on the pre-MRI using the SyN registration approach as implemented in ANTs. To reduce bias introduced by brain shift, the brain shift correction implemented in Lead-DBS was performed on the post-operative CT. Manual pre-reconstruction was then performed, in which the tip of each electrode and another point on the electrode trajectory were marked manually, followed by automatic reconstruction guided by the electrode specification (*i.e.*, number of contacts and spacing between them). The reconstructed electrodes were then visually inspected and refined, and the processes iterated in case of any misalignments. Once all electrodes (for any given patient) have been reconstructed, they were visualized using Lead-Group^74^. Following this process, each electrode contact was associated with MNI coordinates. To identify contacts in our cortical regions of interest, a custom MATLAB code, together with the findStructure function (https://alivelearn.net/?p=1456), was used. Each MNI coordinate was associated with a label from the Automated Anatomical Labeling (AAL) atlas. For occipital cortex, MNI coordinates labelled as ‘inferior occipital’ or ‘middle occipital’ were used; for fusiform, the ‘fusiform’ label was selected. The selected contacts were then visually inspected with the AAL overlay in MRIcron, as well as in native space (see Figure 1d and S1).

### Behavioural task

Patients performed a visuo-motor mapping task, consisting of 12 blocks with 40 trials per block, using the same design as ^19^. Each trial included a brief presentation of a coloured shape, for 500ms, with an inter-stimulus interval of 2200ms. In all trials within a block, two colours and two shapes were combined to form four possible outcomes, with different stimuli presented in the different blocks. Patients were asked to respond to the sampled item by pressing a key to identify the target’s position in the row (Figure 1a). Each trial used an independent sample from a distribution that remained constant within a block, but that varied over blocks. There was no underlying sequence governing stimulus presentation, only the relative proportions of stimuli were varied from block to block. Two information theoretic measures, entropy and surprise, were then calculated (see Figure 1b). Surprise quantifies the improbability of a given event:

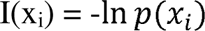

Entropy quantifies the expected (running average of) surprise over all the trials:

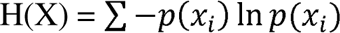

Patients were told the proportion of coloured-shapes presented within block was random, and independent of the other blocks in the task. To account for non-specific time effects within block, the mean entropy over all blocks was calculated for each patient. Trial number is therefore accounted for by the inclusion of mean entropy values which are the same across blocks. Therefore, for each trial, three values of interest were computed: entropy, surprise, and mean entropy. These values were modelled from the perspective of an ideal Bayesian observer, using the Dirichlet distribution ^19^. Only correct responses, given within 1s from stimulus onset, were used for subsequent analyses.

### Electrophysiological data analysis

*Pre-processing*. iEEG data analysis was carried out using the FieldTrip toolbox^68^ (https://www.fieldtriptoolbox.org54) running on MATLAB R2019b (the MathWorks, Natick, MA, USA). For all patient and regions of interest, recordings were transformed to a bipolar derivation by subtracting signal from adjacent electrode contacts within the region of interest (hippocampus, occipital cortex, and fusiform). Previous studies demonstrate that bipolar referencing optimizes estimates of local activity^76,77^ and connectivity patterns between brain regions^78^, as well as for analysis of sharp wave ripples^54^. Data were epoched from −1s to 1s with respect to stimulus onset (a time window selected *a priori*), demeaned and detrended. For each region of interest, every epoch was visually inspected for artefacts caused by epileptic spikes or electrical noise, first in the time domain and then in the time-frequency domain. Trials with artefacts were excluded from all subsequent analyses. Note that for analyses involving more than one region, only trials that were artefact-free in all regions were used.

### Time-frequency analyses

Time-resolved spectral decomposition was computed for each trial using 7 Slepian multitapers for high frequencies (> 35Hz) and a single Hann taper for low frequencies (< 35Hz). The selected Slepian tapers for the analysis of high frequencies were based on windows with width of 0.4s and a 10Hz frequency smoothing. The time-resolved spectral estimation was done in steps of 2.5Hz. No baseline correction was performed, given our interest in both pre- and post-stimulus activity. Trial-wise time-frequency estimates were then entered into a GLM; two predictors of interest (entropy, surprise) and a covariate (mean entropy) were used to predict power at each time-frequency point. The parameter estimates for entropy, mean entropy and surprise are proportional to the range these values can assume. This resulted in a ‘first level’ beta-map (with size time x frequency) per predictor. The spectral activity was then averaged over bipolar channels, within each region, for each patient (in some cases, there was only one bipolar channel in each brain region). These beta-maps were then used for statistical inference with a cluster-based permutation test with the maximum number of permutations allowed by our sample size, up to a maximum of 5000 permutations. In each permutation step, clusters were formed by temporal and frequency adjacency using a cluster-forming threshold of p = 0.05. In each permutation step, a one sample two-tailed t-test against a value of 0 (equivalent to H0: f3 = 0), using a threshold of p = 0.025, was calculated for each time-frequency estimate, separately for low (2.5–32.5Hz) and high (35–160Hz) frequencies, as well as pre-stimulus (−1 to 0s) and post-stimulus (0 to 1s) time windows. Gamma frequency band is used to refer to frequencies 30-100Hz, whereas high gamma refers to effects above 100Hz. The precise frequency ranges (and associated time windows) that we report for each significant effect are derived from a cluster-based permutation correction for multiple comparisons.

### Sharp wave ripple analyses

Ripples were detected using a previously established method (45; Figure S2a). Following trial-wise artefact rejection and basic pre-processing (described above), the iEEG signal was bandpass filtered between 80-120 Hz using a second-order Butterworth filter. A Hilbert transformation was then applied to the filtered signal, extracting its instantaneous amplitude. Ripples were identified as events with a maximum amplitude three standard deviations above the mean, and with a duration of at least 25 ms. Ripples that were detected within 15 ms of each other were merged. The start, peak and end time of each ripple were noted. To ensure artefacts or interictal epileptiform discharges (IEDs; Figure S2b, S27-28) were not mistakenly classified as ripples, we used an automated procedure as described in ^54^ to identify and reject IEDs. The iEEG signal was high-pass filtered at 200 Hz and a z-score was calculated based on the gradient and amplitude of the filtered signal. Any time-point exceeding a z-score of 5 was marked as an IED, together with the 100 ms before and after the event. All IEDs were excluded from the analysis, increasing the likelihood of the detected ripples being physiological ^54^. We have also applied another ripple detection method^51^ and replicated our findings (see Extended Data for Figures 2, 4 and 5). For patients who had more than one anterior hippocampal electrodes (e.g. head and body), we examined whether the same ripple was picked up by the different electrodes (within 20ms). In line with previous findings in rodents^79^, we found the vast majority of ripples (96% on average) were not detected in the adjacent electrode along the hippocampal long axis.

We then examined whether ripple occurrence was influenced by peri-stimulus time and information-theoretic measures. To do so, a 2D histogram of ripple peaks count using peri-stimulus time and entropy/surprise was derived for each patient. We normalised each histogram by the total number of trials per entropy/surprise bin, to account for potential imbalances. The normalised histograms were then averaged across participants and tested against a uniform distribution using a Kolmogorov-Smirnov test. Next, we examined ripple duration, calculated as ripple end point – ripple start point (both determined in the detection algorithm as 2 z-scores above average). To identify the bins with the highest ripple counts, we also performed a mixed-effects Gamma regression on the normalized ripple counts (as they follow a right-skewed distribution). Ripple duration was examined as a function of peri-stimulus time and entropy and surprise. This was done using generalised linear mixed-effects models implemented by the lme4 package ^80^ in R (https://www.r-project.org/). Because ripple duration follows a right-skewed distribution, we used a model from the Gamma family with a random intercept for patient using the following syntax: model <-glmer(rip_duration ∼ entropy * peri_stim_time + surprise * peri_stim_time + mean_entropy + (1|patient), family = Gamma(link=log)).

A similar approach using mixed-effects linear regression was used to examine RT, which was normalized using log-transformation. This approach accommodates any individual differences along participants (e.g. the overall ripple rate for each participant).

### Ripple-modulated cortical activity

To examine the temporal relationship between hippocampal ripples and cortical responses, we extracted time-frequency estimates in occipital cortex and fusiform with respect to peri-ripple peak time (−0.2 to 0.2s, selected *a priori*), as described above for peri-stimulus responses, for high and low frequencies. Due to the shorter time-window, the selected Slepian tapers for the analysis of high frequencies were based on windows with width of 0.2s and a 10 Hz frequency smoothing. The time-resolved spectral estimation was done in steps of 5 Hz. These time-frequency estimates were baseline corrected by calculating the relative change with respect to a baseline period of −1.4 to −1s peri-stimulus. Averaged TF estimates across patients were used for statistical inference using cluster-based permutations, as described above. In each permutation step, a one sample two-tailed paired *t*-test was performed between pre- and post-stimulus periods (threshold of p = 0.025). Next, to examine whether pre-stimulus hippocampal ripples modulated post-stimulus cortical responses to surprise we used the same GLM approach as that described above, fitting a trial-wise regression at each time-frequency point with entropy, surprise and mean entropy as predictors (see *Time-frequency analysis*).

### Granger causality analyses

Finally, to evaluate the direction of information flow between the hippocampus and our cortical regions of interest, we calculated spectral non-parametric Granger causality (GC) as a measure of directed functional connectivity using the FieldTrip toolbox, on 500ms time-windows centred on the induced responses observed to surprise in the fusiform gyrus and hippocampus, and the induced responsed to entropy in the occipital cortex and hippocampus. For patients with multiple bipolar channels in any of the regions, the most medial and anterior channel was used to compute GC. Briefly, time-resolved spectral decomposition was computed using 9 Slepian multi-tapers (frequency range 2 to 80 Hz and 10 Hz smoothing). The spectral transfer matrix was then obtained from the Fourier transformation of the data and together with the noise covariance matrix they were used to calculate the total and intrinsic power through which GC is computed. The resulting GC values were then log-transformed to normalize the data for statistical inference. We then compared information in high versus low entropy and surprise (using a median-split) in each direction (hippocampus cortex and cortex ➔ hippocampus) with non-parametric cluster-based permutations (using a dependent samples two-tailed t-test), employing the maximum number of permutations available for each pair of regions.

## Data availability

All data needed to generate the figures, and over which statistics were computed, are available in the following Github repository: https://github.com/frdarya/GenerativeRipples

## Code availability

Analyses codes are available in the following Github repository: https://github.com/frdarya/GenerativeRipples

## Acknowledgments

We thank the patients who took part in the study and the electroencephalography technicians at the Hospital Ruber Internacional and Swiss Epilepsy Center in Zurich, as well as members of the Laboratory for Clinical Neuroscience for helpful discussions.

## Author contributions

B.A.S., K.J.F., designed the experiment. B.A.S., J.S., and R.T. collected data. R.T., L.I., L.S., and A.G.-N. monitored patients and performed clinical evaluation. N.L. and A.H. developed software to perform electrode localization. D.F., S.M., and B.A.S. performed analyses. D.F., K.J.F., and B.A.S wrote the paper with input from all other authors.

## Extended Data

**Extended Data for Figure 2.**
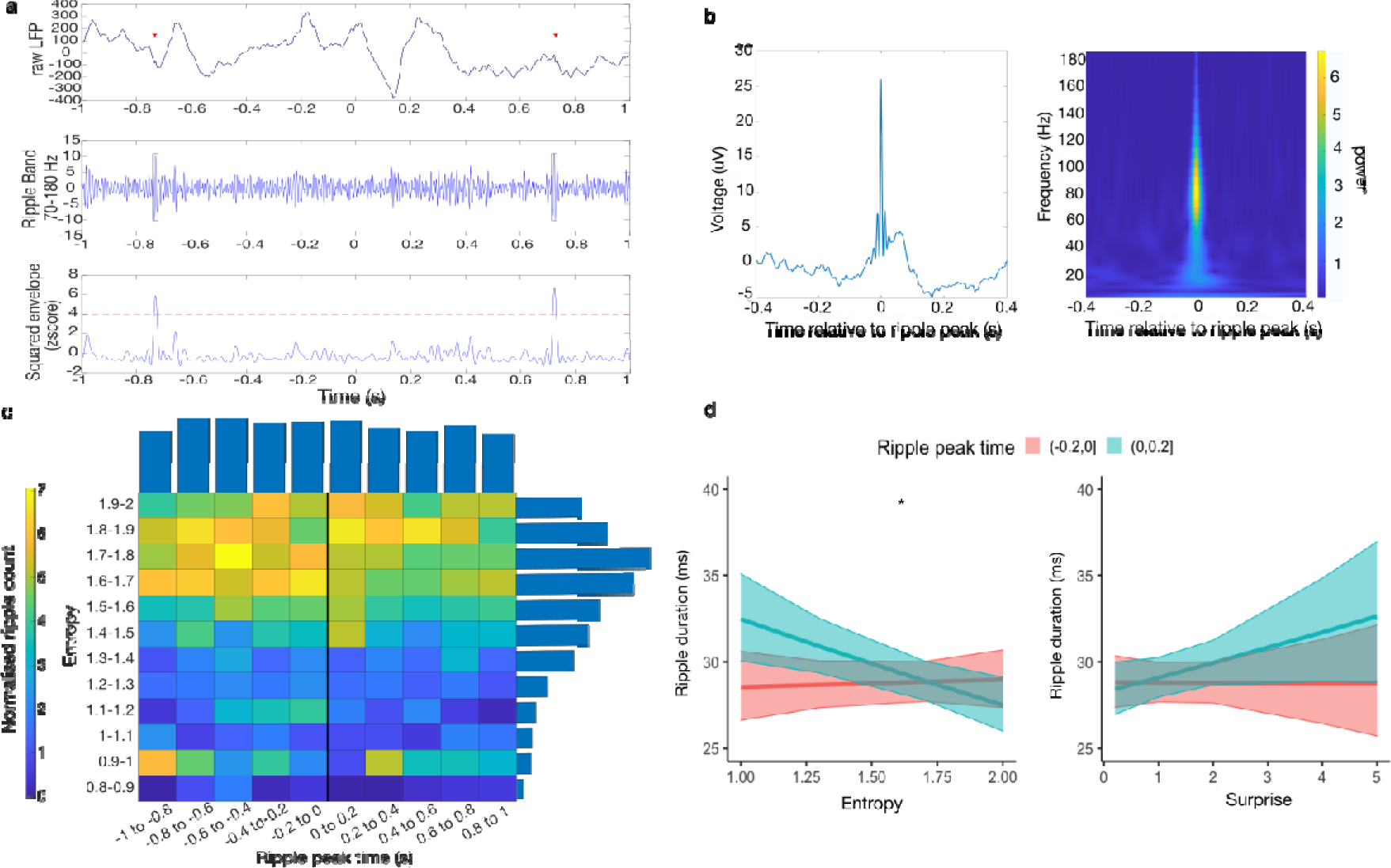
Hippocampal ripples detected using the Norman et al., 2019 algorithm. a) an example ripple detected using the Norman et al (2019) method showing the raw LFP trace (top), the ripple band signal (middle) and the standardised envelope (bottom). b) Grand average raw field potential centred on ripple peak (left) and peri-ripple wavelet spectrogram (n = 3858 ripple events from 15 participants). c) Normalized ripple distribution across peri-stimulus time and entropy levels, accounting for the total number of trials in each entropy bin. Normalized ripple count for each time and entropy bin is colour-coded; average counts for each time and entropy bin are plotted as bars above and to the right of the 2D distribution, respectively. The majority of ripples occurred pre-stimulus and in high (but not highest) entropy levels. d) Ripple duration as a function of peri-stimulus time and information-theoretic measures. Ripple duration increased with entropy (i.e., expected information gain) for pre-stimulus ripples, but decreased for post-stimulus ripples. Shaded areas represent the 95% confidence intervals.

**Extended Data for Figure 4.**
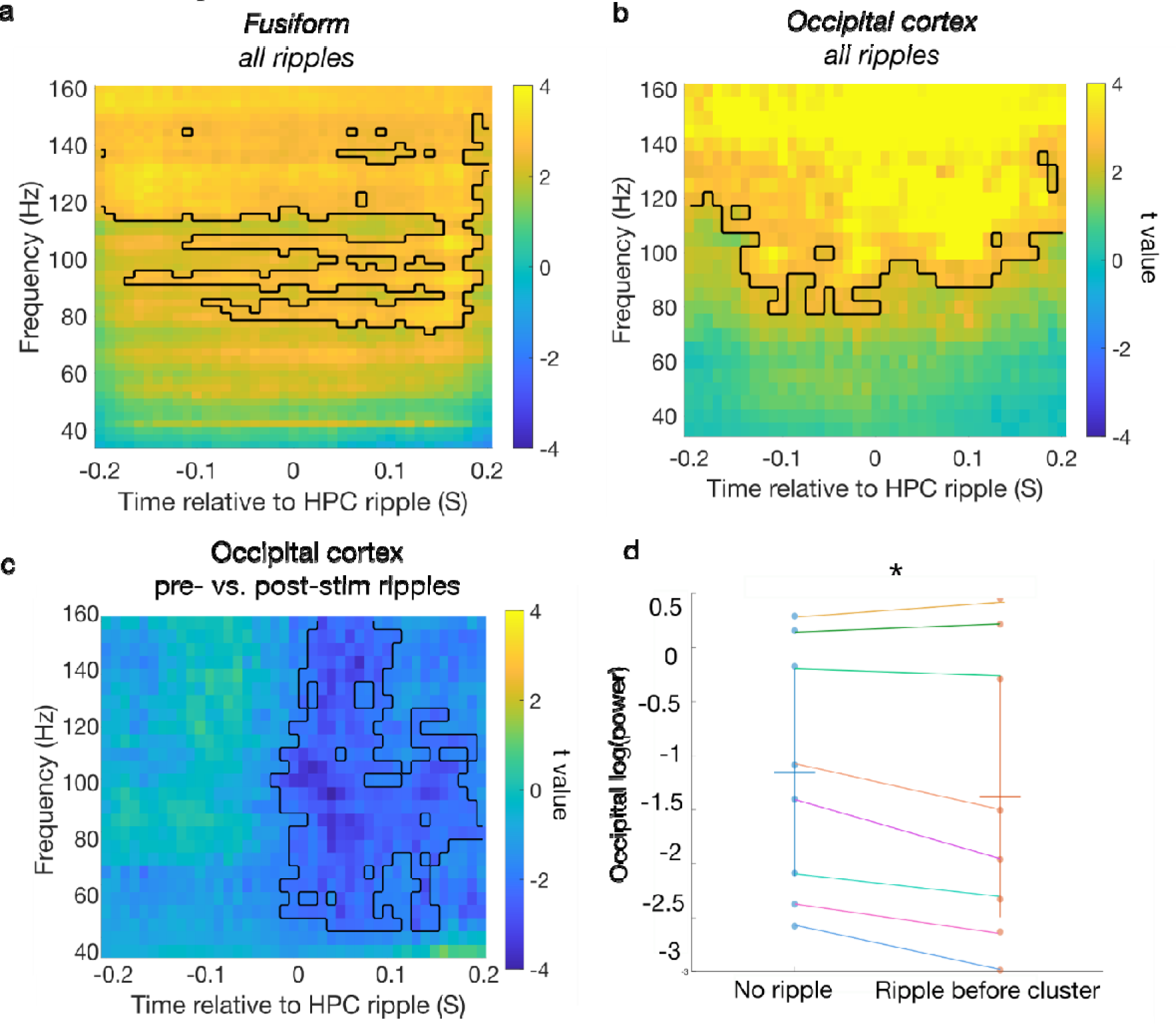
Peri-ripple cortical modulation using the Norman et al., 2019 detection algorithm. Cortical time-frequency analyses time-locked to the hippocampal ripple peak time. Across all ripple trials there was an overall increase in high gamma power in the fusiform (a) and occipital cortex (b) time-locked to hippocampal ripple. c) Comparing trials with pre-versus post-stimulus hippocampal ripples, there was a reduction in occipital gamma power in trials with pre-stimulus ripples, compared to post-stimulus ripples. d) mean of log power in occipital cortex cluster positively associated with entropy in the pre-stimulus time-window, split according to whether there was a hippocampal ripple just before the significant effect or if there was not. Occipital gamma power was suppressed when there was a hippocampal ripple compared to when there was not t(7) = 2.54, p = 0.038, Cohen’s d = 0.9.

**Extended Data for Figure 5.**
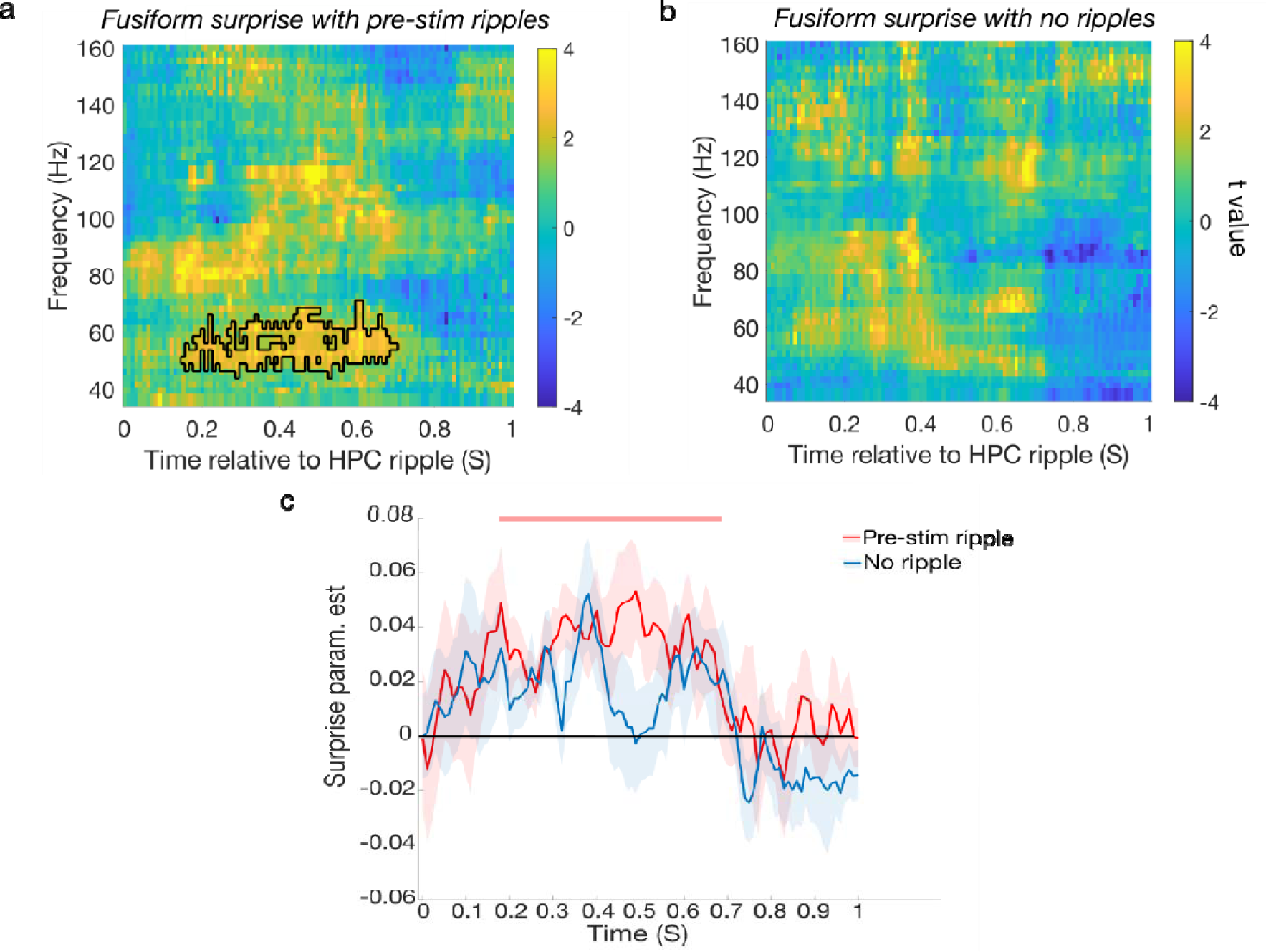
Hippocampal pre-stimulus ripples modulate fusiform response to surprise using ripples detected with the Norman et al., 2019 algorithm. a) Fusiform post-stimulus association with surprise in trials with pre-stimulus ripple compared to (b) without pre-stimulus ripples (no significant response to surprise observed). c) The increase in fusiform gamma power 40-95 Hz (extracted from significant cluster in trials with pre-stimulus hippocampal ripples (red) compared to trials without ripples (blue).

## Supplementary Materials

**Figure S1.**
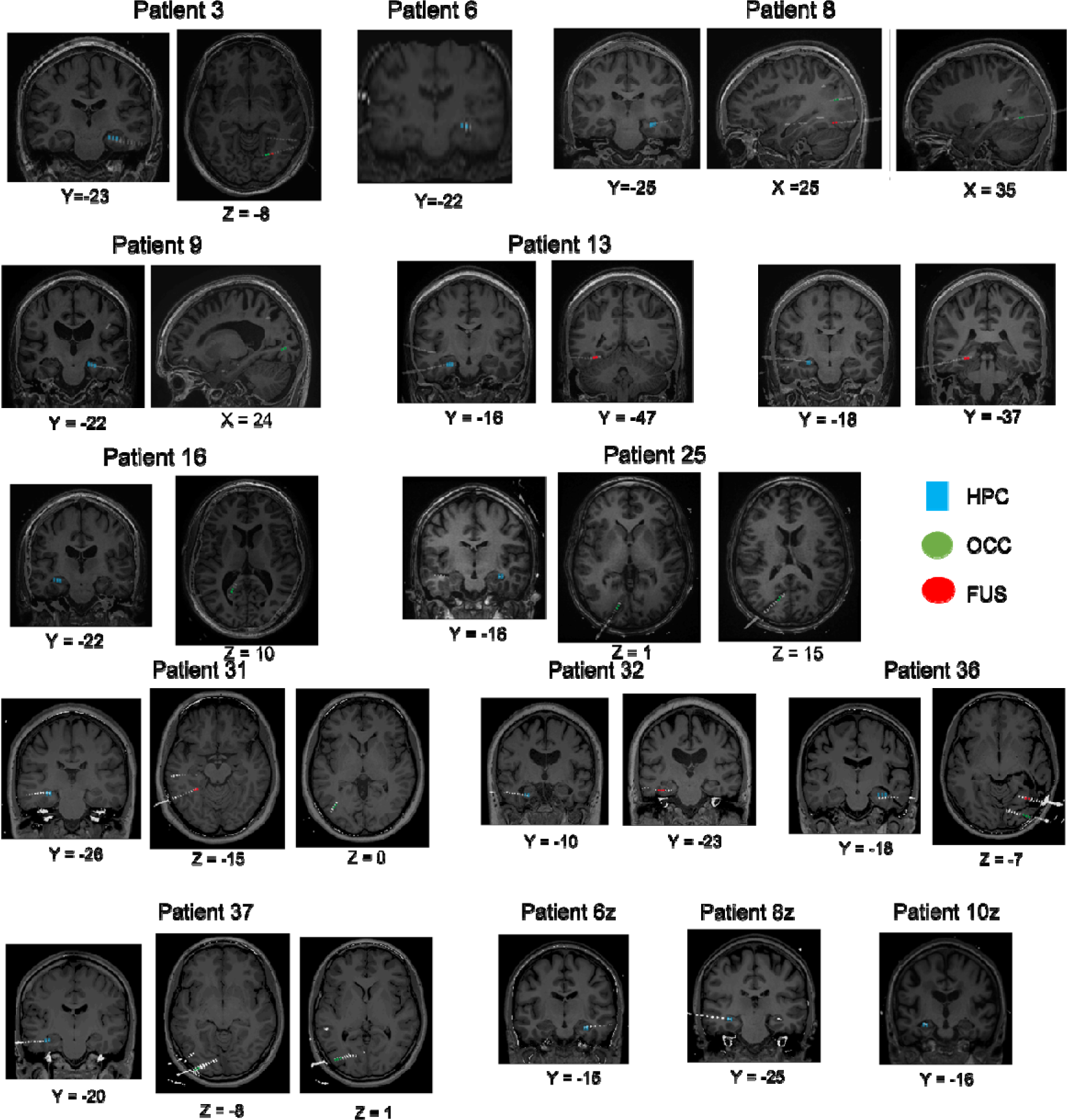
Contact localisation. Patient-specific contacts in the hippocampus (HPC; yellow), occipital cortex (green; OCC) and fusiform (red; FUS). For all patients, post-operative CT images from each patient have been normalised and co-registered with their corresponding pre-operative MRI scans in MNI space and superimposed to display hippocampal, fusiform and occipital cortex contacts (CTs have been thresholded so as to only show electrode contacts).

**Table S1.** Patient demographic and clinical information. M/F male, female: L/R left, right.

| **ID** | **age** | **gender** | **handedness** | **Aetiology** | **Lesion location** |
| --- | --- | --- | --- | --- | --- |
| **3** | 55 | M | R | Focal dysplasia | Right temporal neocortex |
| **6** | 48 | M | R | Hippocampal sclerosis | Left hippocampal sclerosis |
| **8** | 28 | F | R | Focal dysplasia | Extensive right posterior dysplasia involving the convexity and medial aspect of parietal, occipital and posterior temporal lobes |
| **9** | 38 | M | R | Focal dysplasia | Right temporal lobe |
| **13** | 42 | F | R | Focal dysplasia | Right basal temporal cortex |
| **15** | 35 | F | R | Focal dysplasia | Left temporal pole |
| **16** | 30 | M | R | Reactive gliosis, diffuse microglia activation and small vessel vasculopathy | Medial wall of the left parietal region (precuneus and posterior cingulum) |
| **25** | 29 | M | R | Focal dysplasia | Right posterior temporobasal region |
| **31** | 48 | F | R | Drug-resistant epilepsy | Left frontal or fronto-temporal |
| **32** | 59 | M | R | Encephalocele + gliosis | Bilateral temporal pole (crisis registered only from the left side) |
| **36** | 28 | F | R | Focal epilepsy secondary to refractory right temporo-parietal hematoma |  |
| **37** | 24 | F | L | Non-lesional |  |
| **6z** | 29 | F | R | Unclear |  |
| **8z** | 30 | F | R | Non-lesional |  |
| **10z** | 56 | M | R | Hippocampal sclerosis | Right hippocampus |

**Figure S2.**
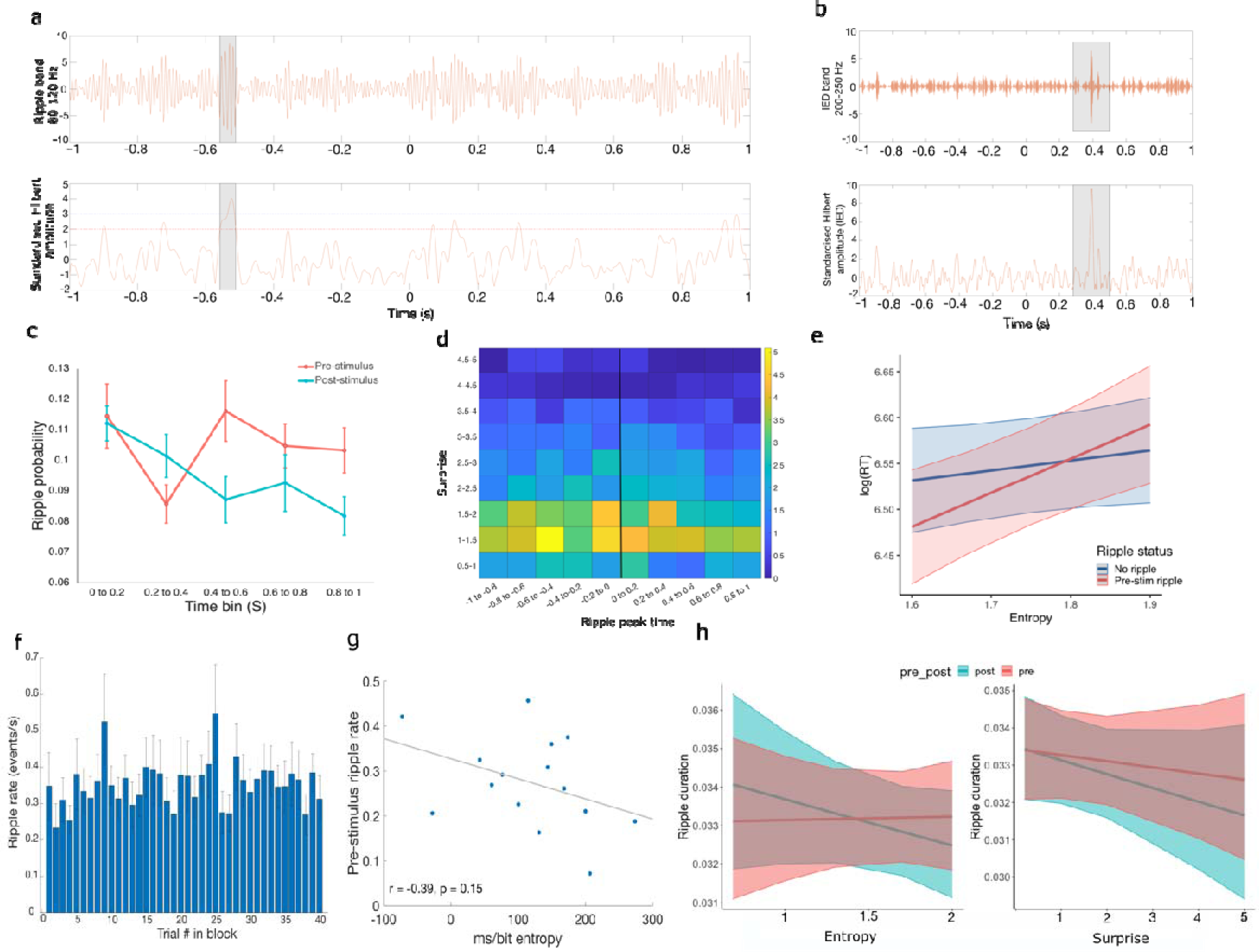
Hippocampal ripples. **a)** An example of ripple detection, showing the filtered ripple band and Hilbert amplitude thresholds. Grey shading indicates ripple duration. **b)** An example interictal epileptiform discharge (IED). **c)** Ripple frequency as a function of time-bin and peri-stimulus time; a trend towards a main effect of peri-stimulus time (p = 0.061), with more ripples observed pre-stimulus. **d)** Normalised distribution of ripples as a function of peri-stimulus time and level of stimulus-bound surprise. **e)** Reaction time (log normalized) as a function of pre-stimulus ripple status and entropy. For trials with a pre-stimulus ripple there was a faster response, more pronounced (at trend level) for entropy levels around 1.6 to 1.8 bits. **f)** Ripple rate as a function of trial in block. **g)** Correlation between pre-stimulus ripple rate and increase in reaction time as a function of ms per bit of entropy**. h)** Ripple duration as a function peri-stimulus time and information-theoretic measures.

**Figure S3.**
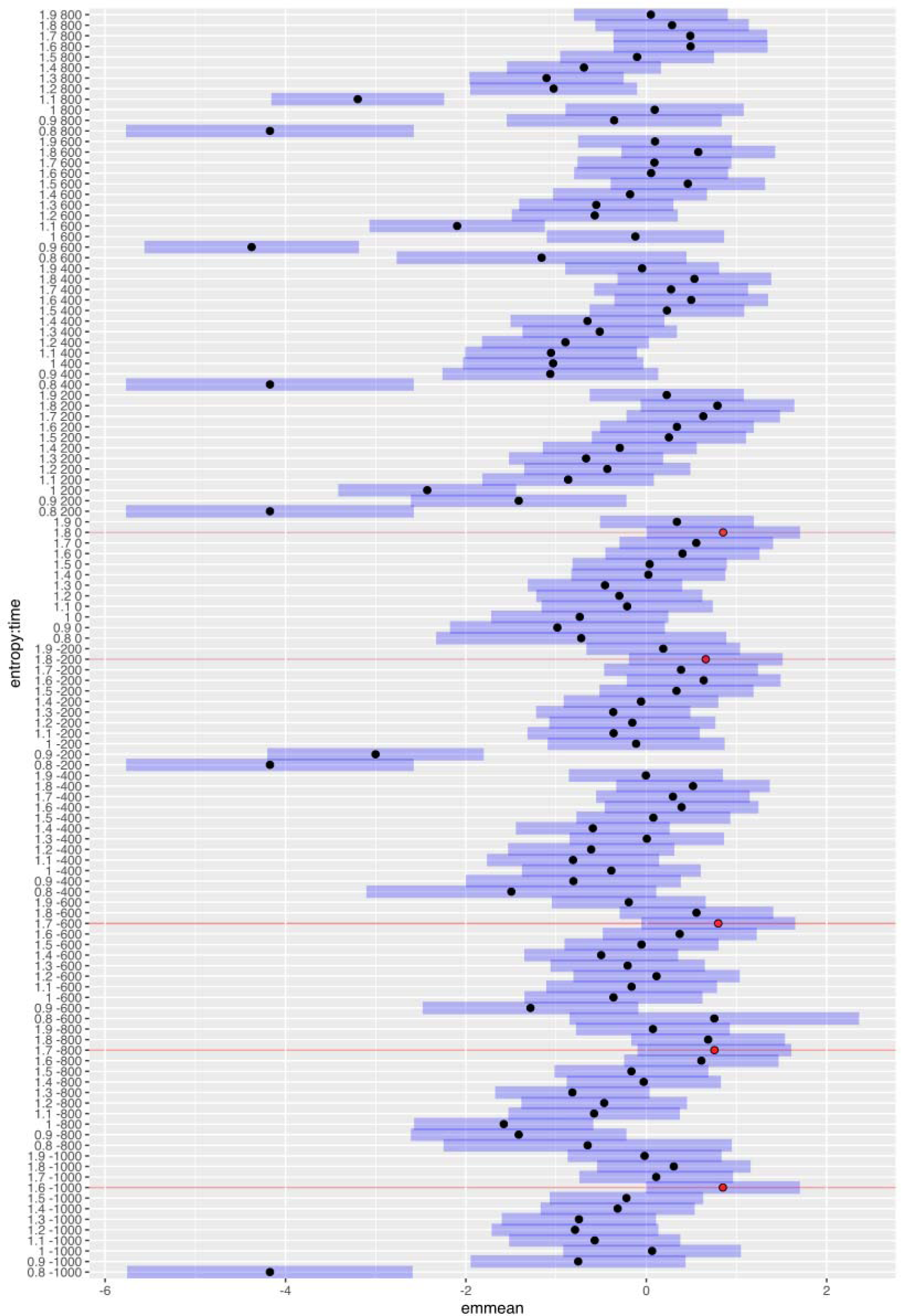
Estimated marginal means of the 2D ripple count as a function of entropy and time bins. The top 5 estimated marginal means are marked in red, blue shadings represent upper and lower 95% confidence interval.

**Figure S4.**
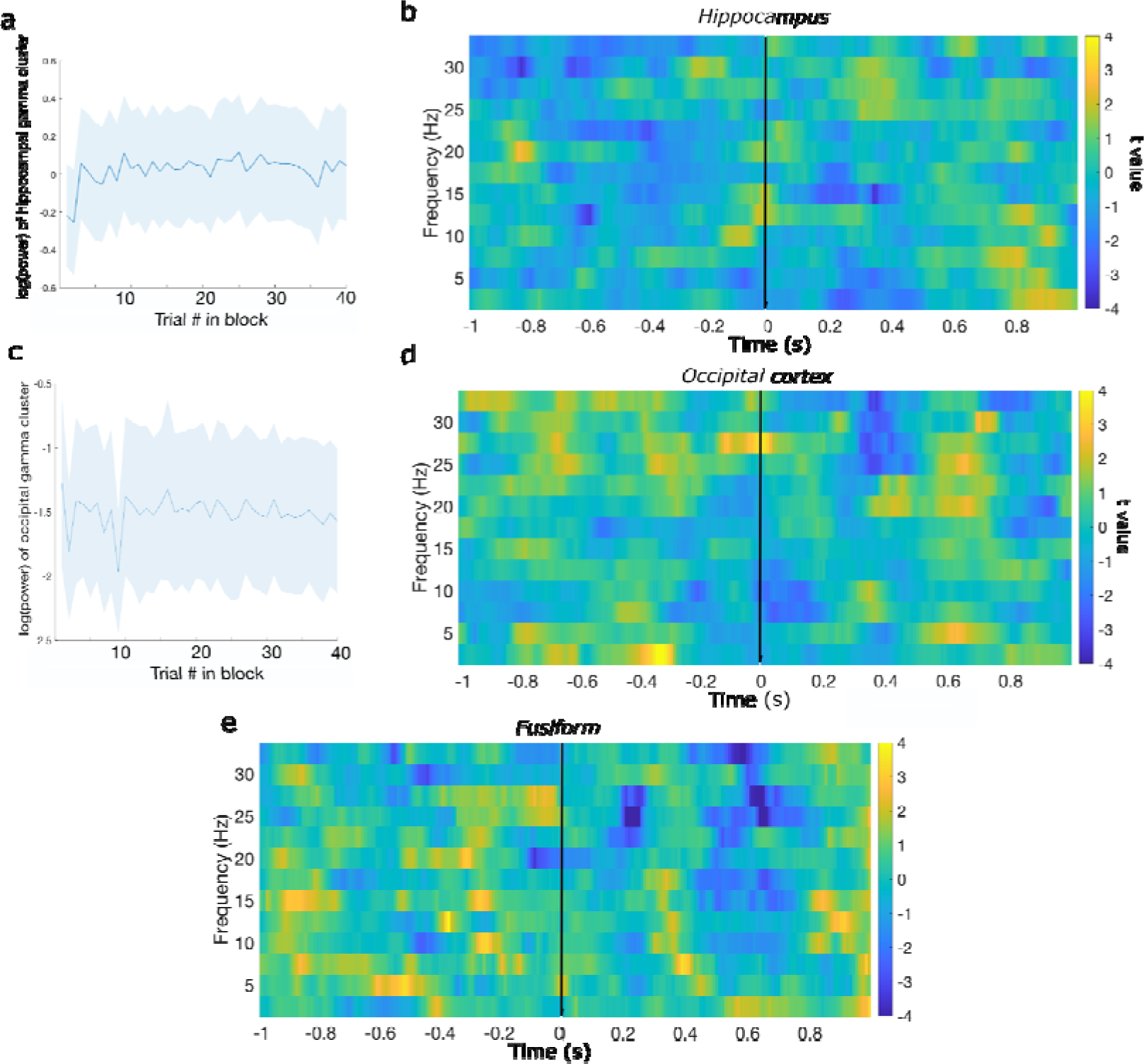
Entropy-related responses. **a**) Log-transformed gamma power in hippocampal pre-stimulus entropy cluster (shown in Figure 3a), as a function of trial in block. **b**) Hippocampal lower frequencies entropy responses, pre- and post-stimulus. **c**) Log-transformed gamma power in occipital cortex pre-stimulus entropy cluster (shown in Figure 3b), as a function of trial in block. **d**) Occipital and **e**) Fusiform cortex lower frequencies entropy responses, pre- and post-stimulus.

**Figure S5.**
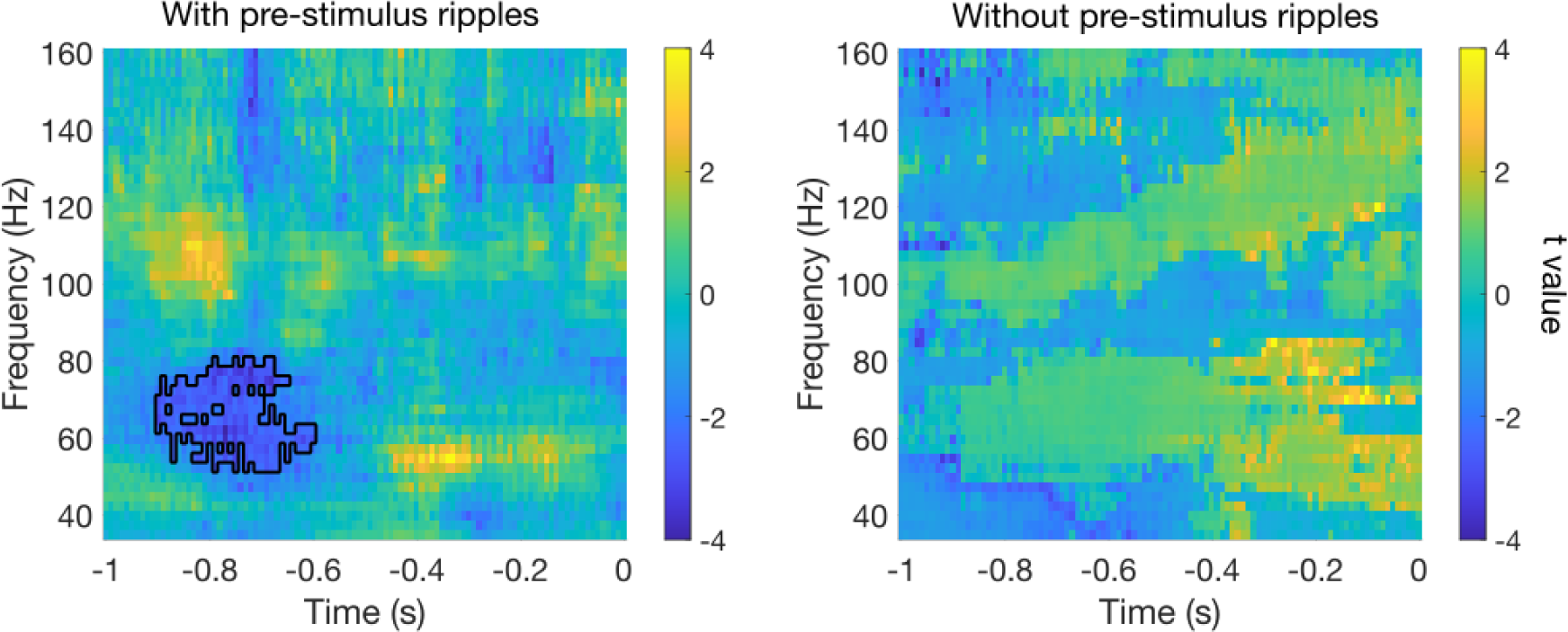
Pre-stimulus hippocampal response to entropy split by whether a pre-stimulus ripple was present.

**Figure S6.**
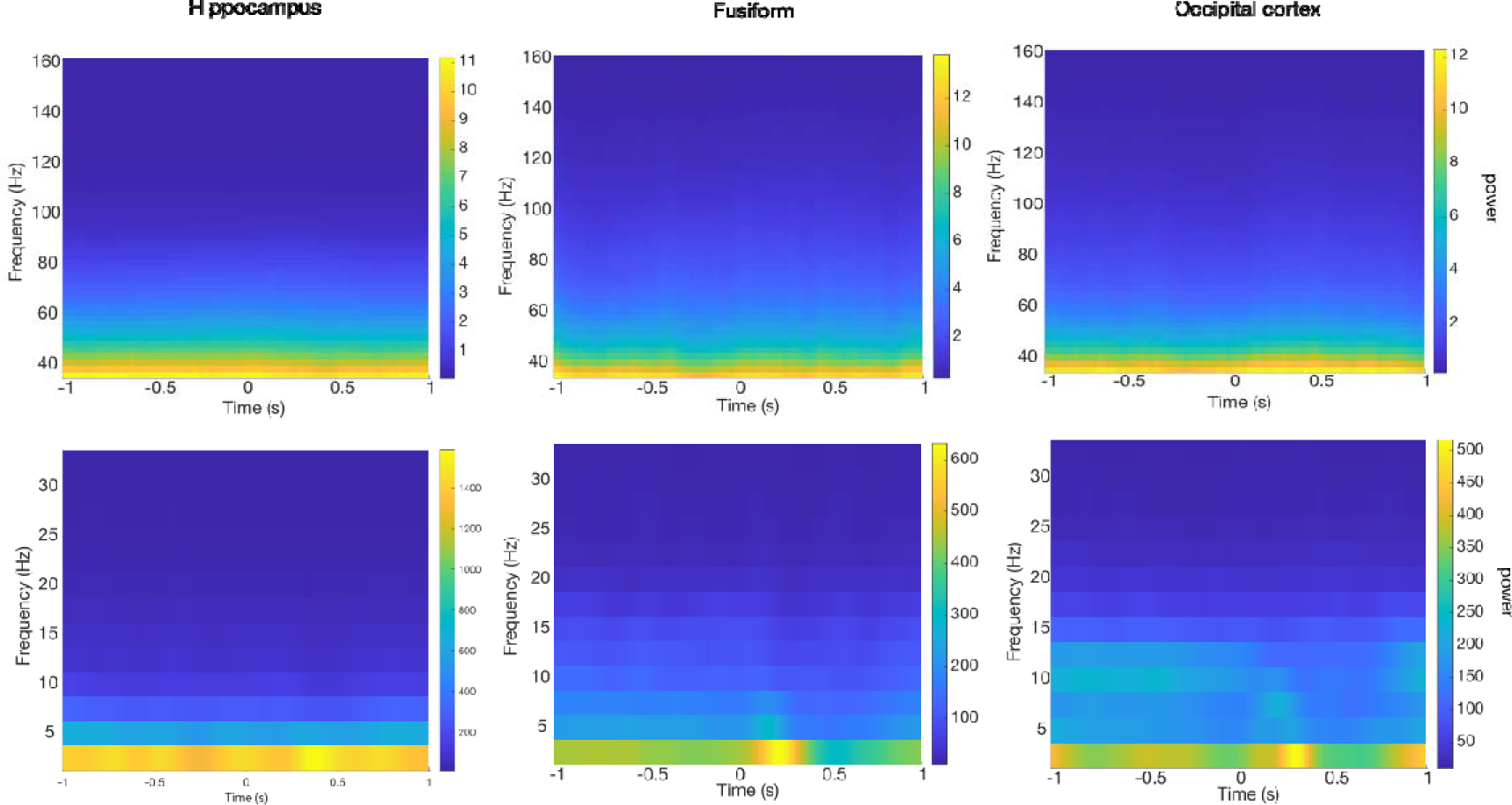
Raw spectrograms across all trials in each of the region of interest.

**Figure S7.**
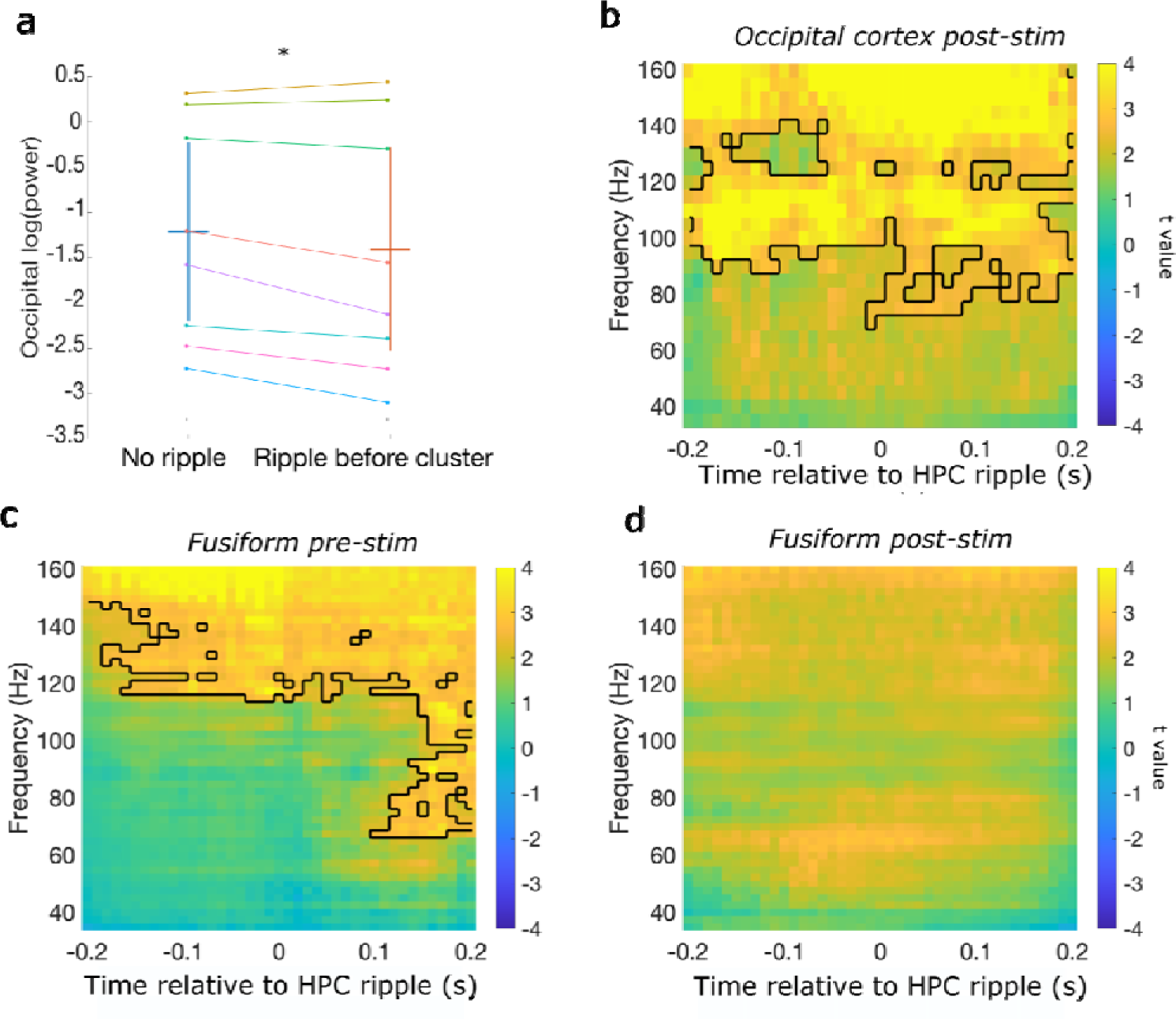
Peri-ripple cortical modulation. (**a**) mean of log power in occipital cortex cluster positively associated with entropy in the pre-stimulus time-window, split by whether there was a hippocampal ripple just before the significant effect or if there was not. Occipital gamma power was suppressed when there was a hippocampal ripple compared to when there was not. (**b**) T-statistics of peri-ripple activity in occipital cortex for post-stimulus ripples. (**c-d**) Peri-ripple activity in the fusiform for (**c**) pre- and (**d)** post-stimulus ripples. * p < 0.05

**Figure S8.**
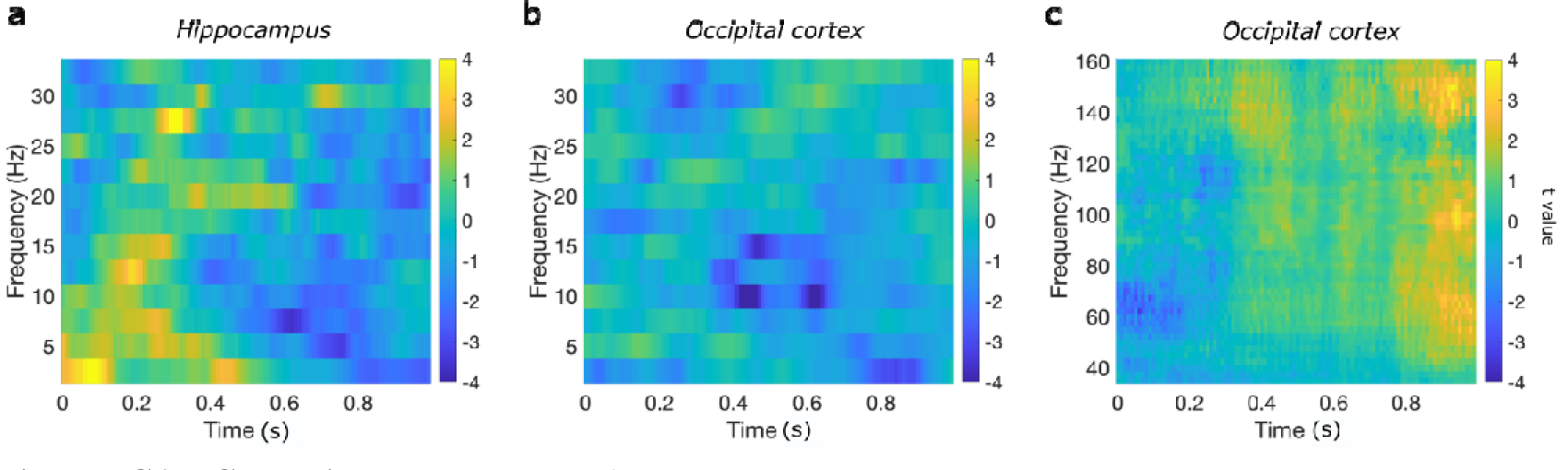
Surprise responses. **a)**Hippocampal low frequency response to surprise. (**b-c**) Occipital cortex (**b)** low and **(c)** high frequencies response to surprise.

**Figure S9.**
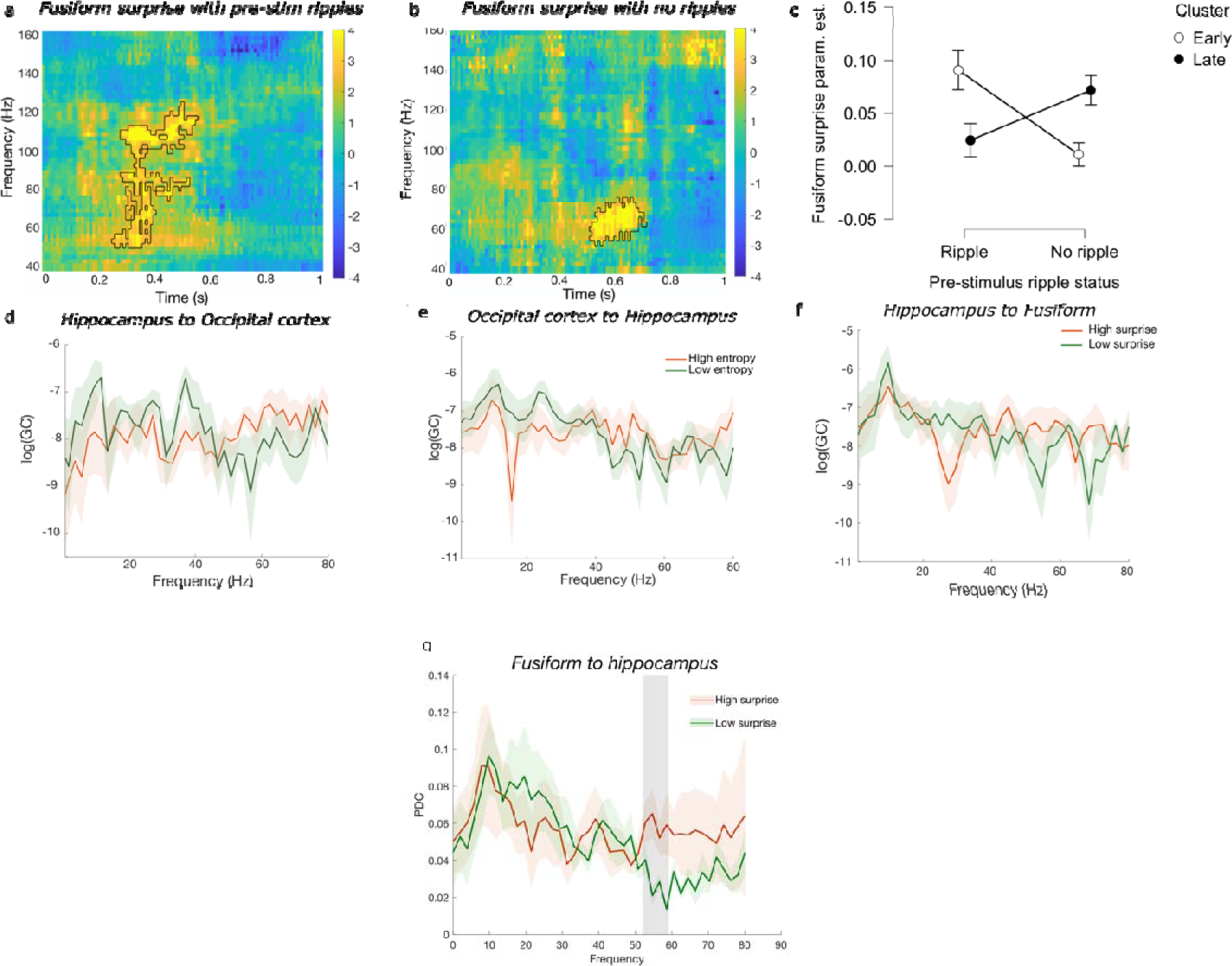
Fusiform post-stimulus response to surprise is modulated by pre-stimulus hippocampal ripples and directed cortical-hippocampal information flow. **a)** Fusiform post-stimulus association with surprise in trials with pre-stimulus ripple earlier than **b)** without pre-stimulus ripples; **d-e**) Granger causality (GC) analysis between the hippocampus and occipital cortex for entropy. **f)** Granger causality from the hippocampus to fusiform for surprise. **g)** Partial directed coherence values for high (orange) versus low (green) surprise trials were significantly higher in the gamma band (shaded rectangle). This is analogous to the results presented in Figure 5e (main text), employing a Granger casual analysis.

**Figure S10.**
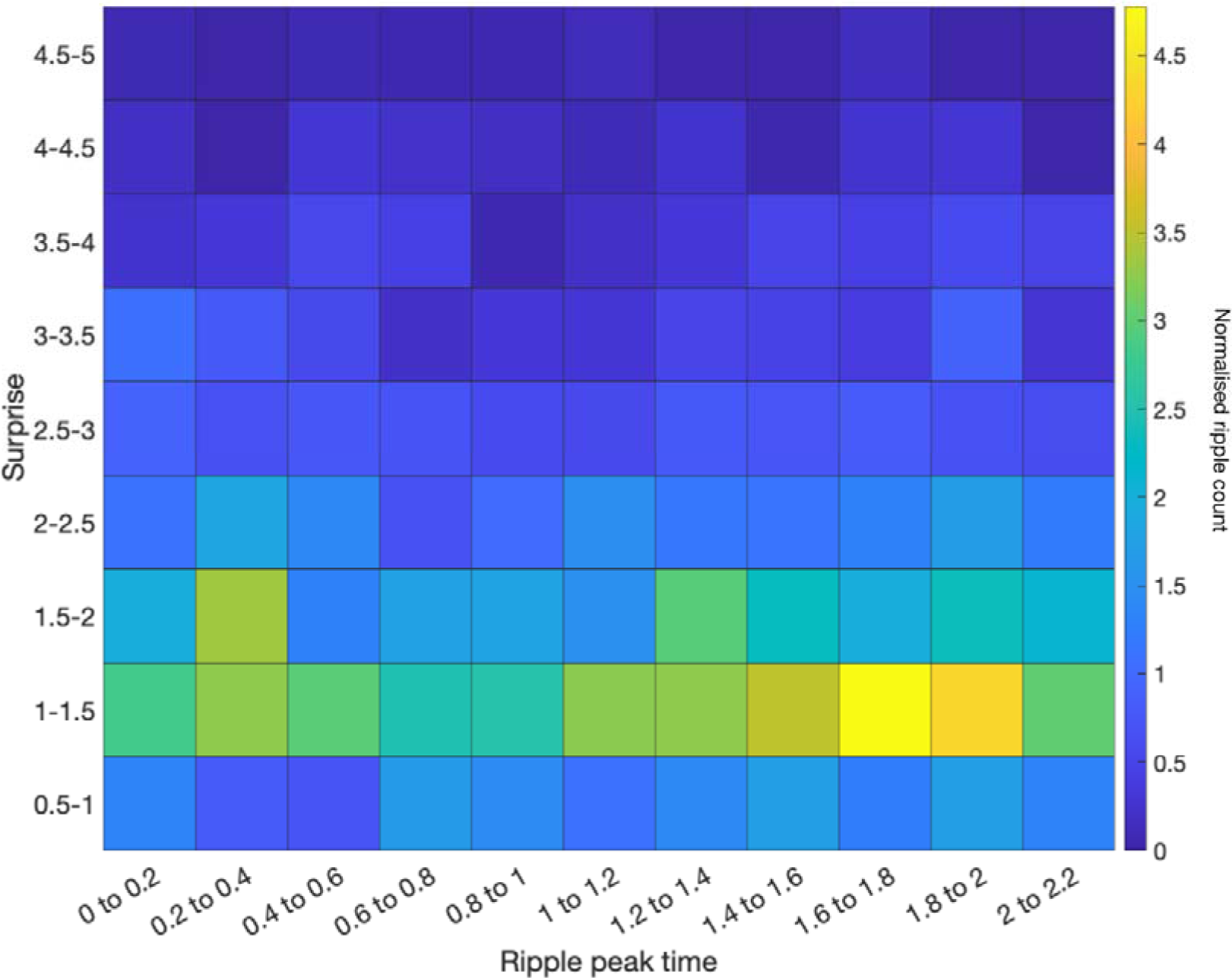
Ripple distribution as a function of surprise. If pre-stimulus ripples were reflecting post-processing of the previous stimulus, an increase in ripple count should be observed for high surprise later in the trial. When locked to stimulus onset and going forward in time (up to 2.2s post-stimulus onset, which corresponds to the time between presentation of successive stimuli), such an increase was not observed, i.e., the ripple rate in the top right quadrant of the plot shows a low ripple rate.. Instead, there is an increase in ripple count for low surprise (1-1.5 bits) at later time bins. This range of surprise values corresponds, on average, to a value of 1.67 bits of entropy. This plot is, therefore, consistent with the increase in ripple count in the pre-stimulus period for upcoming entropy values around 1.6 to 1.9 bits.

**Figure S11.**
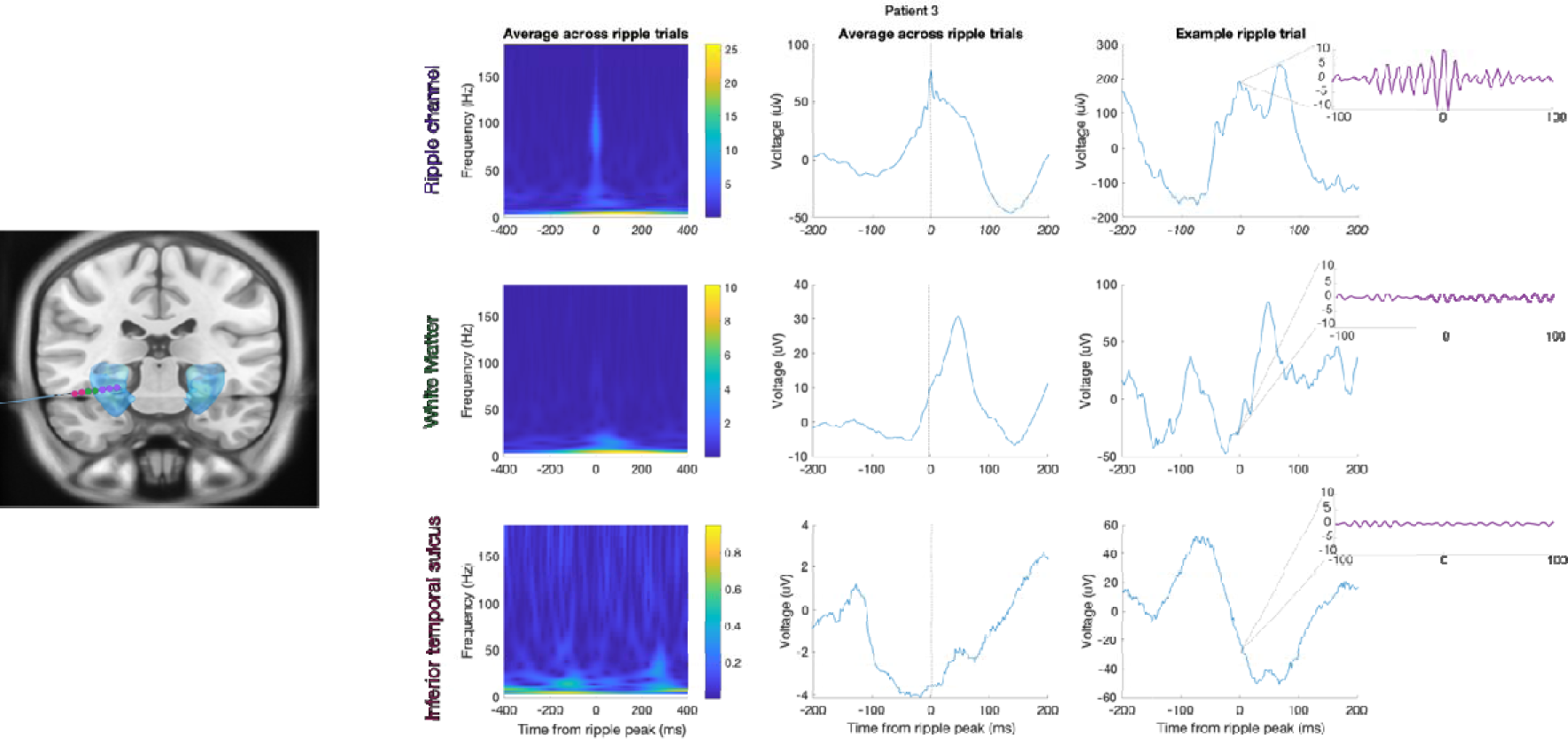
Averaged, and example single trial, ripple traces from hippocampal and adjacent contacts for Patient 3. *Left*. Electrode localisation shown on the smoothed hippocampus mask from the Automated Anatomical Labelling atlas for visualization; blue) illustrates the contacts (coloured circles) from which data are presented in the panels on the right. Purple circles: 3 hippocampal contacts i.e., two bipolar channels. Green circles: the two contacts adjacent to the hippocampus are in white matter (medial contact)/inferior temporal sulcus (lateral contact). Red circles: the two subsequent contacts are in inferior temporal sulcus (medial contact)/white matter (lateral contact). *Right*. Top, bottom and middle rows: ripple-locked activity from the hippocampal (Ripple) channels (purple circles), and adjacent bipolar channels “White matter” (corresponding to green circles) and “inferior temporal sulcus” (corresponding to red circles). The left and centre columns show the average peri-ripple wavelet spectrogram across all ripple trials (n = 320), and average raw field potential centred on ripple peak, respectively. The right column shows an example ripple trial (raw field potential centred on ripple peak) with an inset showing the ripple band activity (band pass filtered at 80-120Hz).

**Figure S12.**
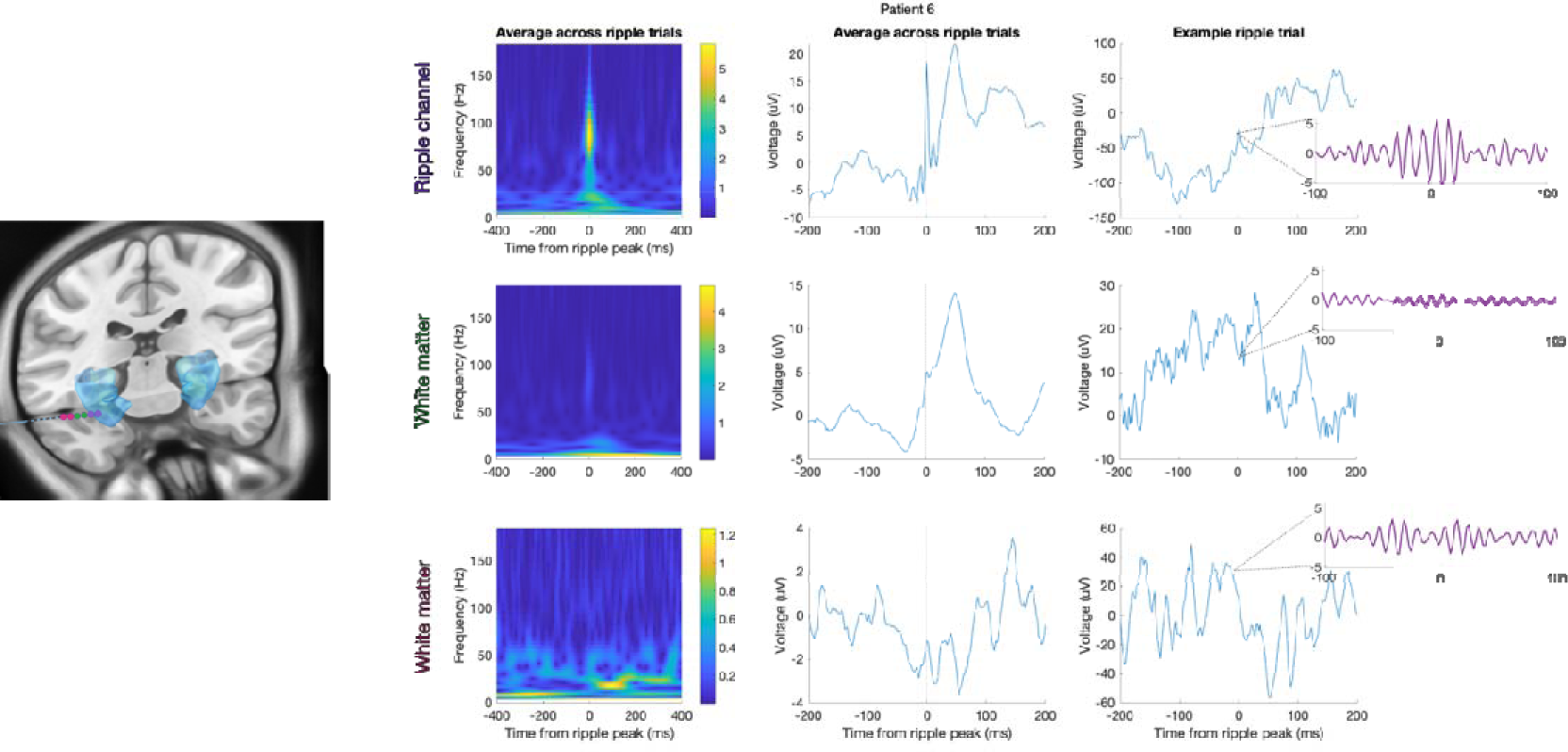
Averaged, and example single trial, ripple traces from hippocampal and adjacent contacts for Patient 6. *Left*. Electrode localisation shown on the smoothed hippocampus mask from the Automated Anatomical Labelling atlas for visualization; blue) illustrates the contacts (coloured circles) from which data are presented in the panels on the right. Purple circles: 2 hippocampal contacts i.e., one bipolar channel. Green circles: the two contacts adjacent to the hippocampus are in white matter. Red circles: the two subsequent contacts are in white matter. *Right*. Top, bottom and middle rows: ripple-locked activity from the hippocampal (Ripple) channel (purple circles), and adjacent bipolar channels “White matter” (corresponding to green circles) and “White matter” (corresponding to red circles). The left and centre columns show the average peri-ripple wavelet spectrogram across all ripple trials (n = 160), and average raw field potential centred on ripple peak, respectively. The right column shows an example ripple trial (raw field potential centred on ripple peak) with an inset showing the ripple band activity (band pass filtered at 80-120Hz).

**Figure S13.**
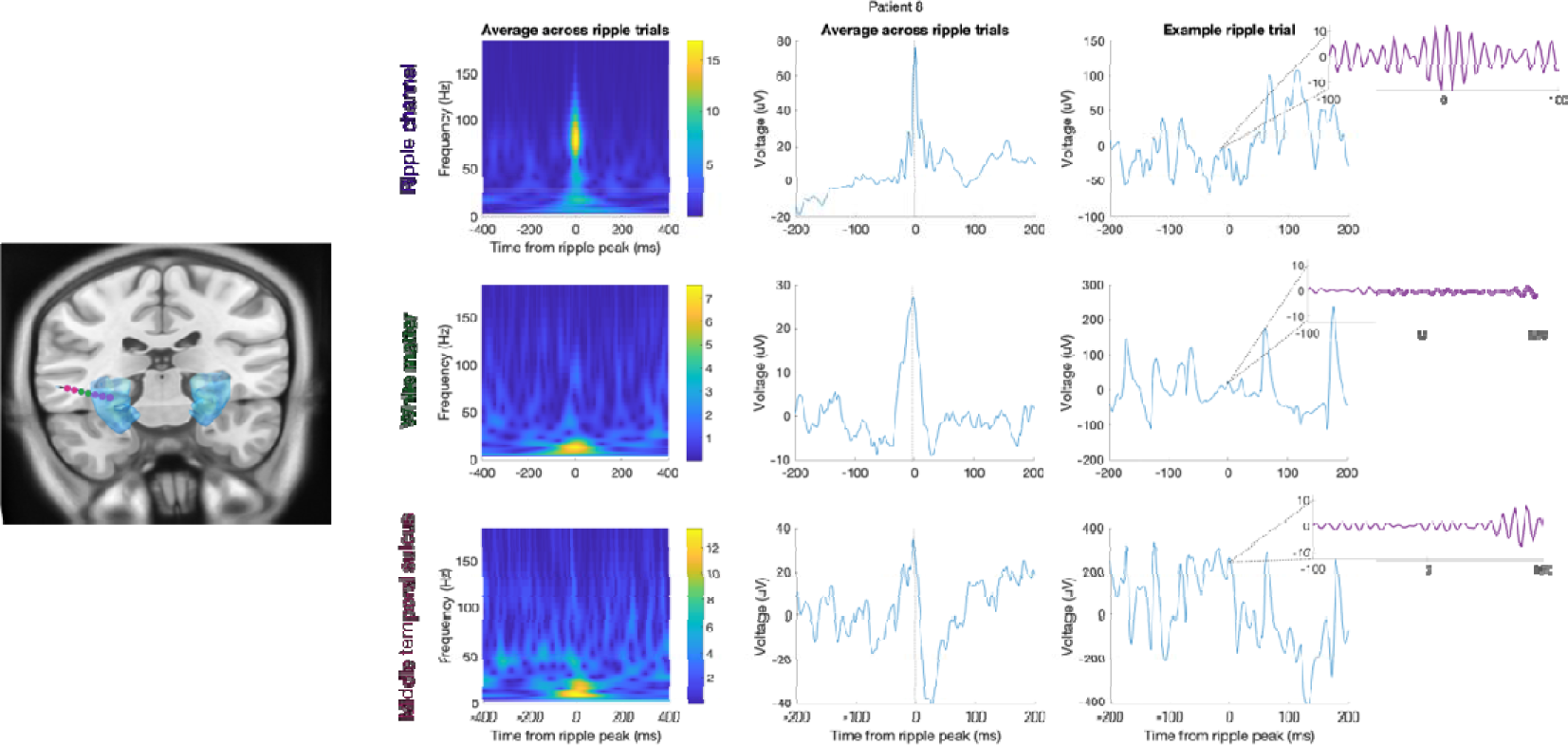
Averaged, and example single trial, ripple traces from hippocampal and adjacent contacts for Patient 8. *Left*. Electrode localisation shown on the smoothed hippocampus mask from the Automated Anatomical Labelling atlas for visualization; blue) illustrates the contacts (coloured circles) from which data are presented in the panels on the right. Purple circles: 3 hippocampal contacts i.e., two bipolar channels. Green circles: the two contacts adjacent to the hippocampus are in white matter. Red circles: the two subsequent contacts are in middle temporal sulcus. *Right*. Top, bottom and middle rows: ripple-locked activity from the hippocampal (Ripple) channels (purple circles), and adjacent bipolar channels “White matter” (corresponding to green circles) and “inferior temporal sulcus” (corresponding to red circles). The left and centre columns show the average peri-ripple wavelet spectrogram across all ripple trials (n = 210), and average raw field potential centred on ripple peak, respectively. The right column shows an example ripple trial (raw field potential centred on ripple peak) with an inset showing the ripple band activity (band pass filtered at 80-120Hz).

**Figure S14.**
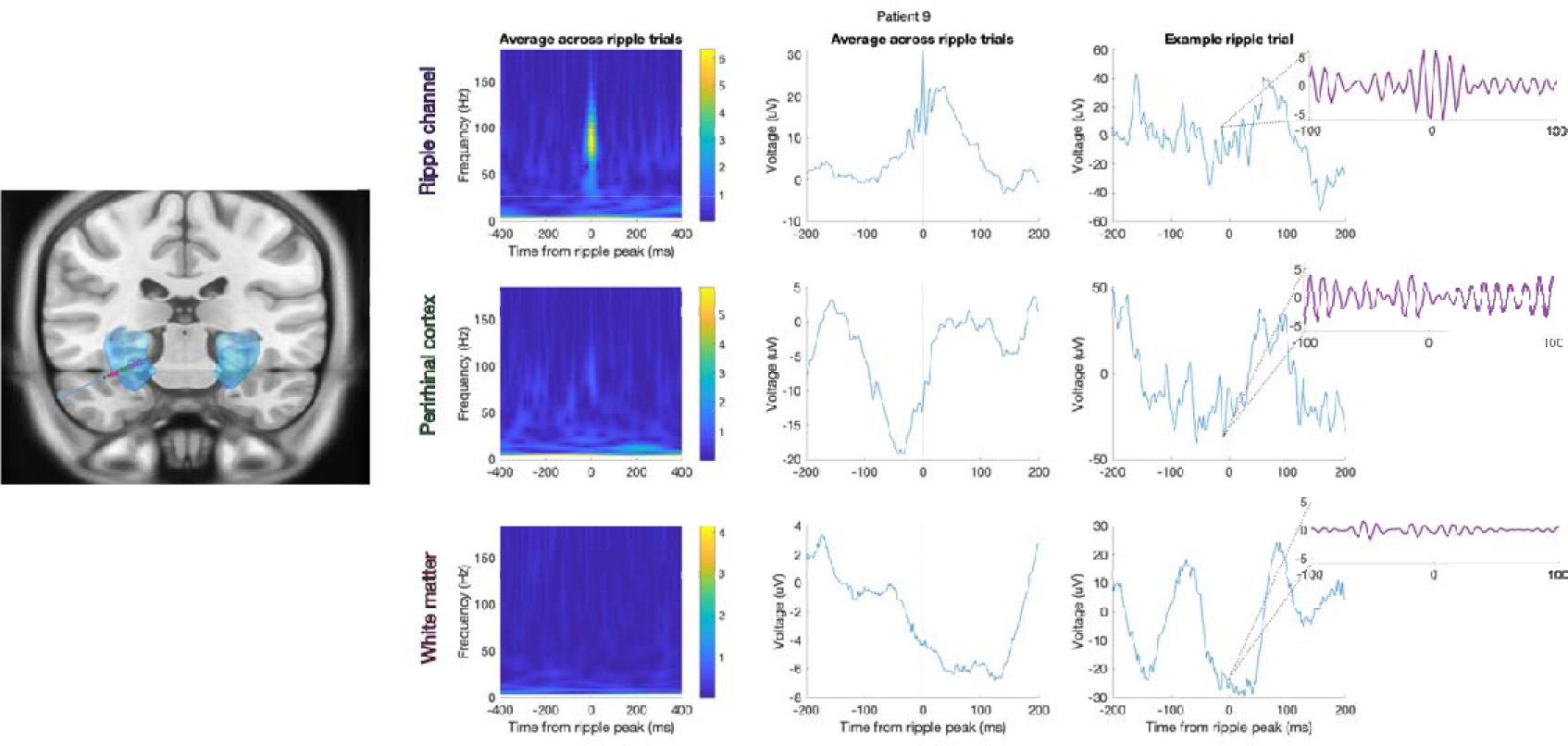
Averaged, and example single trial, ripple traces from hippocampal and adjacent contacts for Patient 9. *Left*. Electrode localisation shown on the smoothed hippocampus mask from the Automated Anatomical Labelling atlas for visualization; blue) illustrates the contacts (coloured circles) from which data are presented in the panels on the right. Purple circles: 3 hippocampal contacts i.e., two bipolar channels. Green circles: the two contacts adjacent to the hippocampus are in perirhinal cortex. Red circles: the two subsequent contacts are in white matter. *Right*. Top, bottom and middle rows: ripple-locked activity from the hippocampal (Ripple) channels (purple circles), and adjacent bipolar channels “Perirhinal cortex” (corresponding to green circles) and “White matter” (corresponding to red circles). The left and centre columns show the average peri-ripple wavelet spectrogram across all ripple trials (n = 160), and average raw field potential centred on ripple peak, respectively. The right column shows an example ripple trial (raw field potential centred on ripple peak) with an inset showing the ripple band activity (band pass filtered at 80-120Hz).

**Figure S15.**
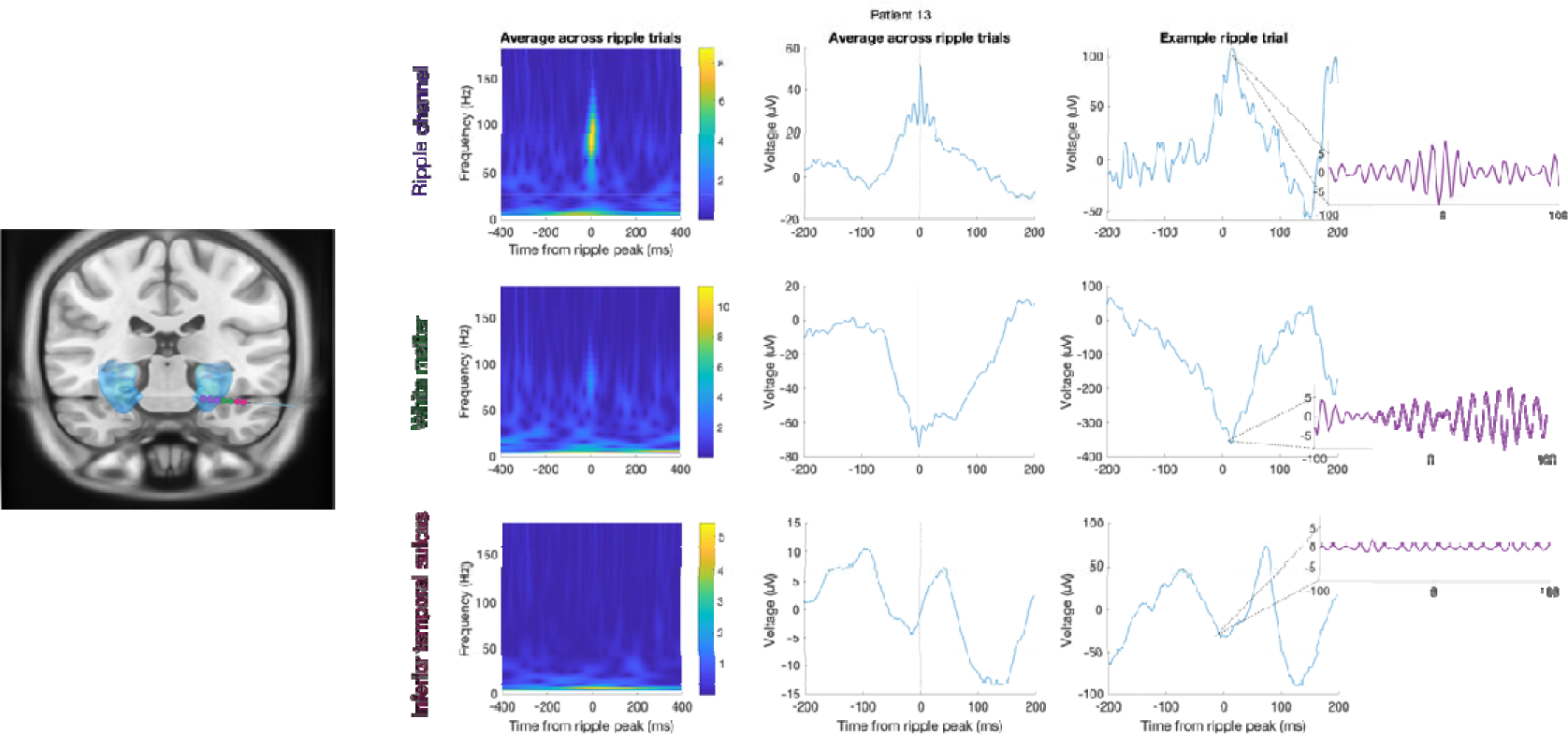
Averaged, and example single trial, ripple traces from hippocampal and adjacent contacts for Patient 13. *Left*. Electrode localisation shown on the smoothed hippocampus mask from the Automated Anatomical Labelling atlas for visualization; blue) illustrates the contacts (coloured circles) from which data are presented in the panels on the right. Purple circles: 3 hippocampal contacts i.e., two bipolar channels. Green circles: the two contacts adjacent to the hippocampus are in white matter. Red circles: the two subsequent contacts are in inferior temporal sulcus. *Right*. Top, bottom and middle rows: ripple-locked activity from the hippocampal (Ripple) channels (purple circles), and adjacent bipolar channels “White matter” (corresponding to green circles) and “inferior temporal sulcus” (corresponding to red circles). The left and centre columns show the average peri-ripple wavelet spectrogram across all ripple trials (n = 122), and average raw field potential centred on ripple peak, respectively. The right column shows an example ripple trial (raw field potential centred on ripple peak) with an inset showing the ripple band activity (band pass filtered at 80-120Hz).

**Figure S16.**
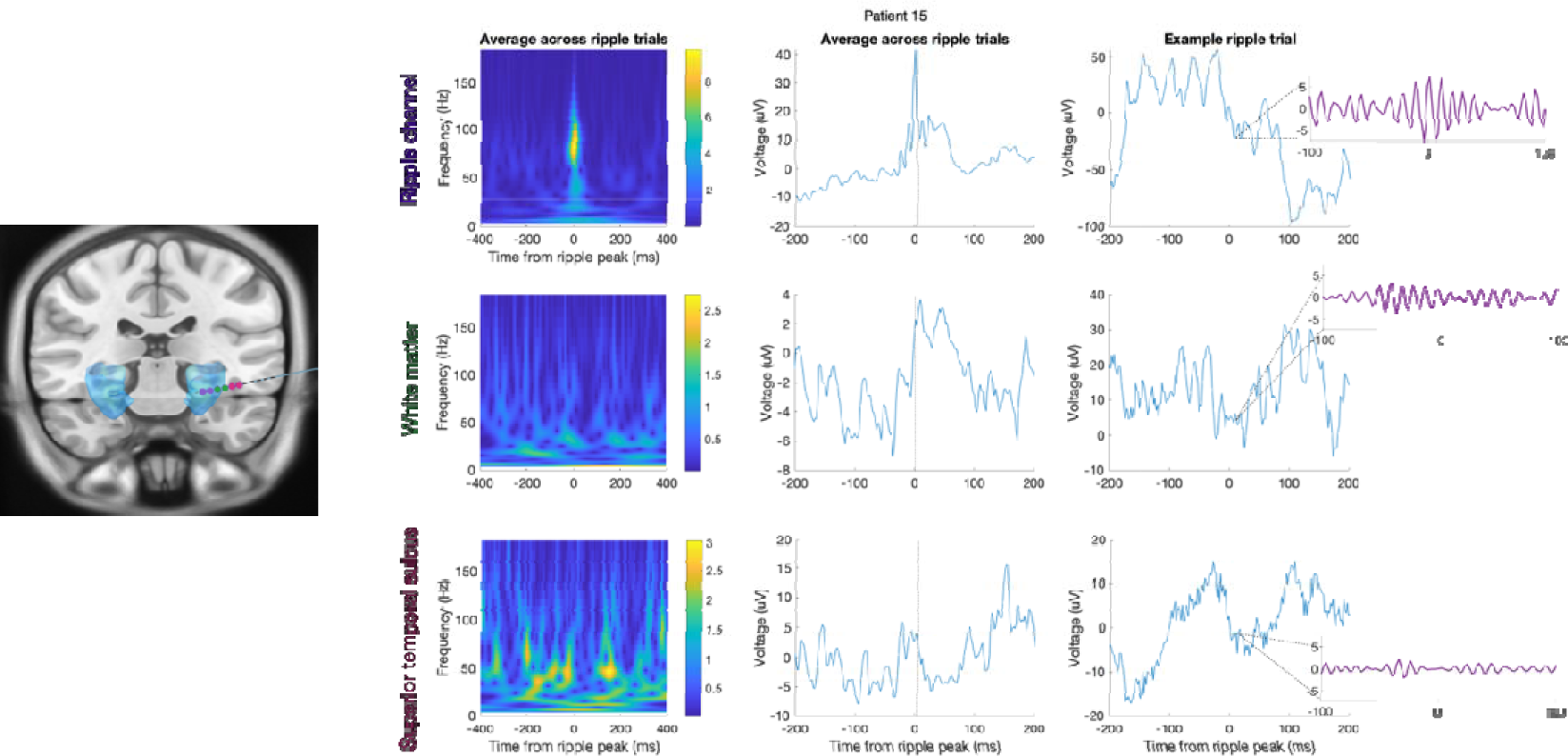
Averaged, and example single trial, ripple traces from hippocampal and adjacent contacts for Patient 15. *Left*. Electrode localisation shown on the smoothed hippocampus mask from the Automated Anatomical Labelling atlas for visualization; blue) illustrates the contacts (coloured circles) from which data are presented in the panels on the right. Purple circles: 2 hippocampal contacts i.e., one bipolar channel. Green circles: the two contacts adjacent to the hippocampus are in white matter. Red circles: the two subsequent contacts are in superior temporal sulcus (medial contact)/white matter (lateral contact). *Right*. Top, bottom and middle rows: ripple-locked activity from the hippocampal (Ripple) channel (purple circles), and adjacent bipolar channels “White matter” (corresponding to green circles) and “Superior temporal sulcus” (corresponding to red circles). The left and centre columns show the average peri-ripple wavelet spectrogram across all ripple trials (n = 169), and average raw field potential centred on ripple peak, respectively. The right column shows an example ripple trial (raw field potential centred on ripple peak) with an inset showing the ripple band activity (band pass filtered at 80-120Hz).

**Figure S17.**
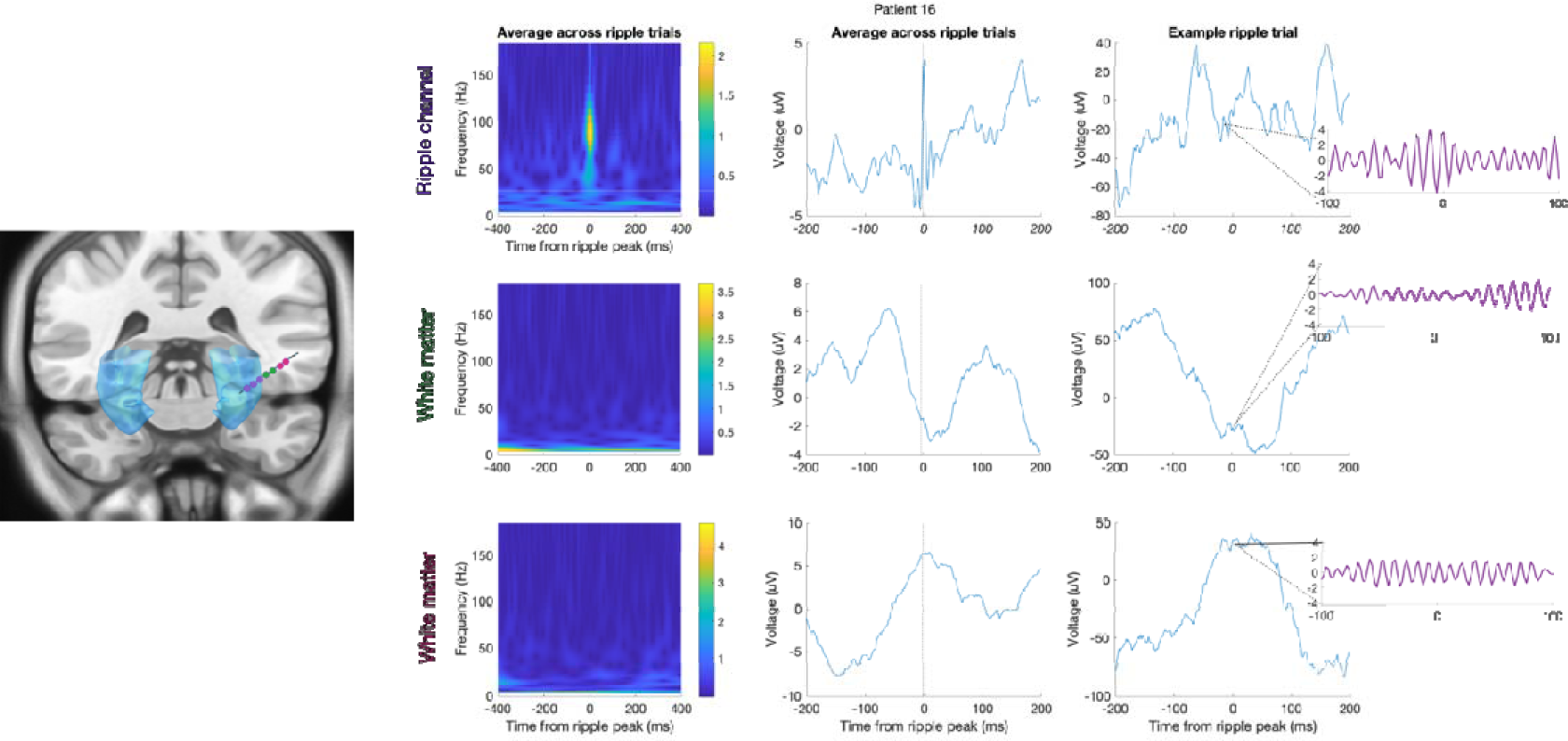
Averaged, and example single trial, ripple traces from hippocampal and adjacent contacts for Patient 16. *Left*. Electrode localisation shown on the smoothed hippocampus mask from the Automated Anatomical Labelling atlas for visualization; blue) illustrates the contacts (coloured circles) from which data are presented in the panels on the right. Purple circles: 2 hippocampal contacts i.e., one bipolar channel. Green circles: the two contacts adjacent to the hippocampus are in white matter. Red circles: the two subsequent contacts are in white matter. *Right*. Top, bottom and middle rows: ripple-locked activity from the hippocampal (Ripple) channel (purple circles), and adjacent bipolar channels “White matter” (corresponding to green circles) and “White matter” (corresponding to red circles). The left and centre columns show the average peri-ripple wavelet spectrogram across all ripple trials (n = 266), and average raw field potential centred on ripple peak, respectively. The right column shows an example ripple trial (raw field potential centred on ripple peak) with an inset showing the ripple band activity (band pass filtered at 80-120Hz).

**Figure S18.**
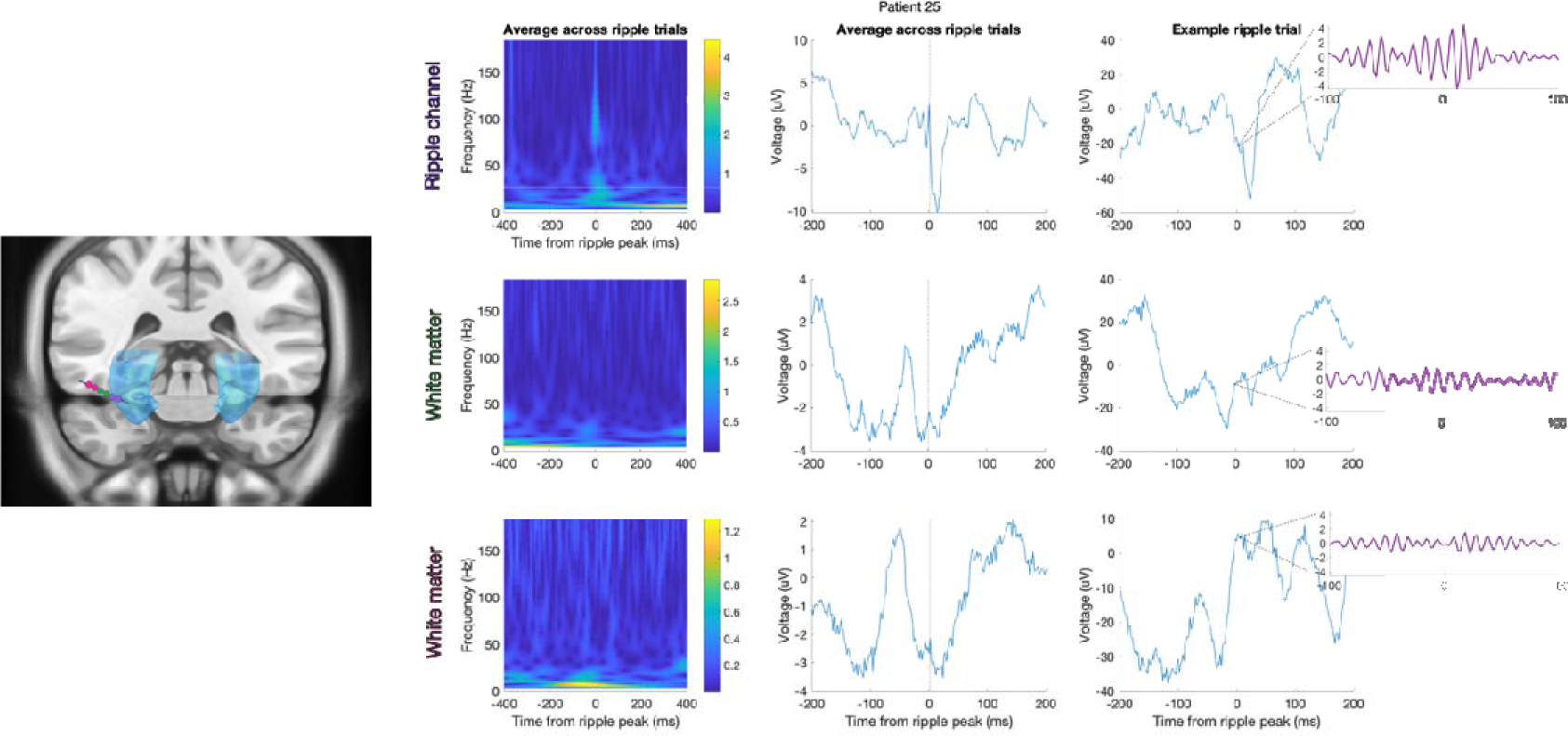
Averaged, and example single trial, ripple traces from hippocampal and adjacent contacts for Patient 25. *Left*. Electrode localisation shown on the smoothed hippocampus mask from the Automated Anatomical Labelling atlas for visualization; blue) illustrates the contacts (coloured circles) from which data are presented in the panels on the right. Purple circles: 2 hippocampal contacts i.e., one bipolar channel. Green circles: the two contacts adjacent to the hippocampus are in white matter. Red circles: the two subsequent contacts are in white matter. *Right*. Top, bottom and middle rows: ripple-locked activity from the hippocampal (Ripple) channels (purple circles), and adjacent bipolar channels “White matter” (corresponding to green circles) and “White matter” (corresponding to red circles). The left and centre columns show the average peri-ripple wavelet spectrogram across all ripple trials (n = 49), and average raw field potential centred on ripple peak, respectively. The right column shows an example ripple trial (raw field potential centred on ripple peak) with an inset showing the ripple band activity (band pass filtered at 80-120Hz).

**Figure S19.**
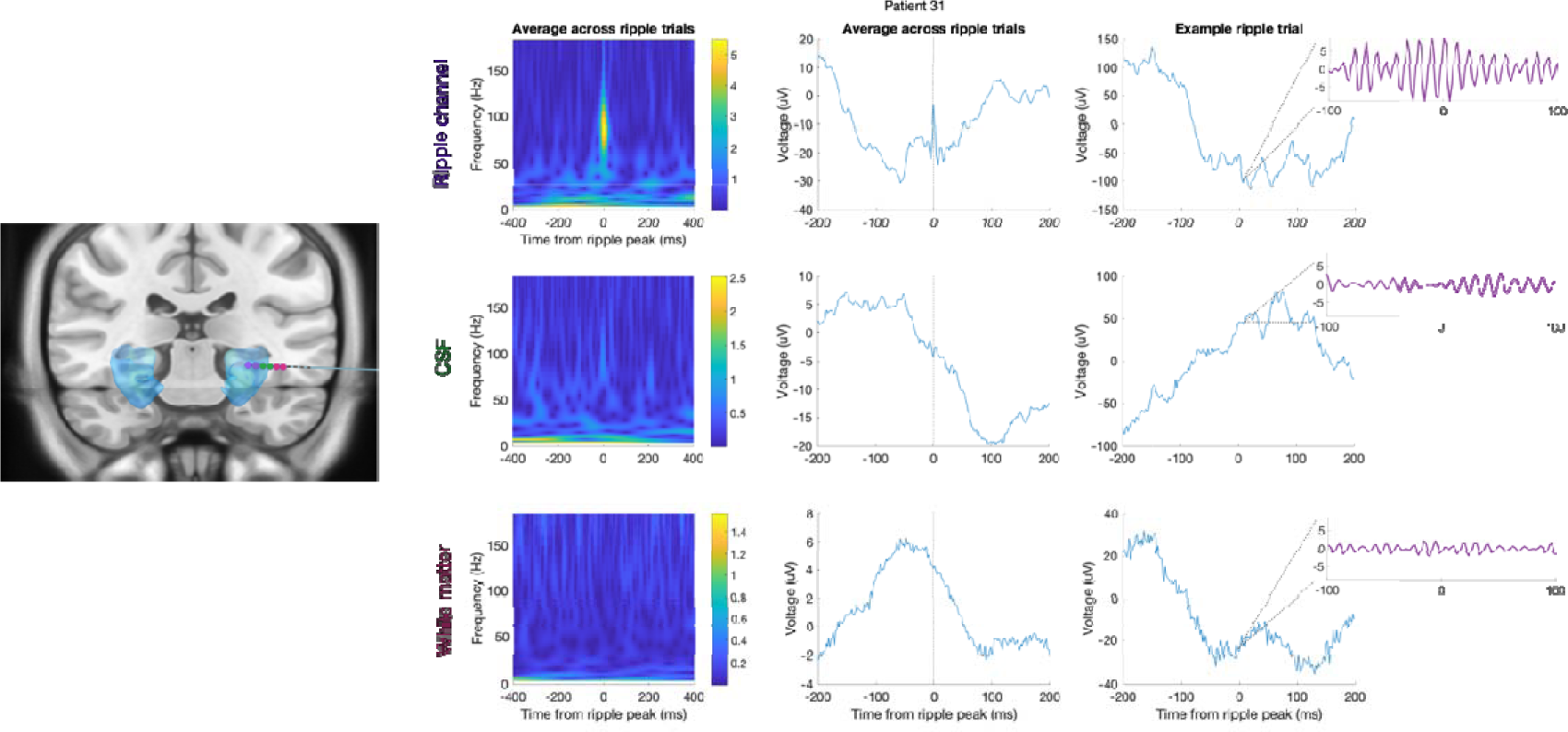
Averaged, and example single trial, ripple traces from hippocampal and adjacent contacts for Patient 31. *Left*. Electrode localisation shown on the smoothed hippocampus mask from the Automated Anatomical Labelling atlas for visualization; blue) illustrates the contacts (coloured circles) from which data are presented in the panels on the right. Purple circles: 2 hippocampal contacts i.e., one bipolar channel. Green circles: the two contacts adjacent to the hippocampus are in CSF (medial contact)/white matter (lateral contact). Red circles: the two subsequent contacts are in white matter. *Right*. Top, bottom and middle rows: ripple-locked activity from the hippocampal (Ripple) channel (purple circles), and adjacent bipolar channels “CSF” (corresponding to green circles) and “White matter” (corresponding to red circles). The left and centre columns show the average peri-ripple wavelet spectrogram across all ripple trials (n = 100), and average raw field potential centred on ripple peak, respectively. The right column shows an example ripple trial (raw field potential centred on ripple peak) with an inset showing the ripple band activity (band pass filtered at 80-120Hz).

**Figure S20.**
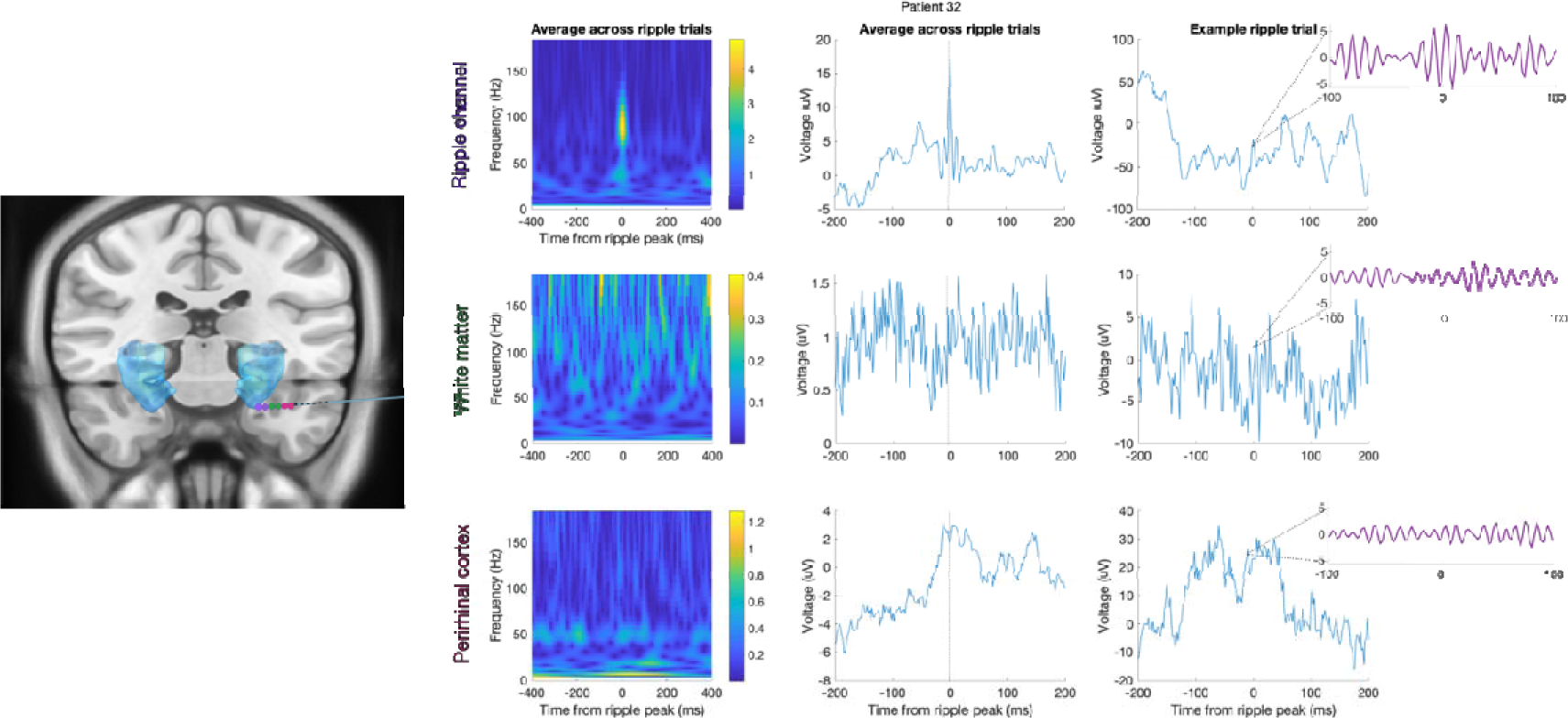
Averaged, and example single trial, ripple traces from hippocampal and adjacent contacts for Patient 32. *Left*. Electrode localisation shown on the smoothed hippocampus mask from the Automated Anatomical Labelling atlas for visualization; blue) illustrates the contacts (coloured circles) from which data are presented in the panels on the right. Purple circles: 2 hippocampal contacts i.e., one bipolar channel. Green circles: the two contacts adjacent to the hippocampus are in white matter (medial contact)/CSF (lateral contact). Red circles: the two subsequent contacts are in perirhinal cortex. *Right*. Top, bottom and middle rows: ripple-locked activity from the hippocampal (Ripple) channel (purple circles), and adjacent bipolar channels “White matter” (corresponding to green circles) and “Perirhinal cortex” (corresponding to red circles). The left and centre columns show the average peri-ripple wavelet spectrogram across all ripple trials (n = 120), and average raw field potential centred on ripple peak, respectively. The right column shows an example ripple trial (raw field potential centred on ripple peak) with an inset showing the ripple band activity (band pass filtered at 80-120Hz).

**Figure S21.**
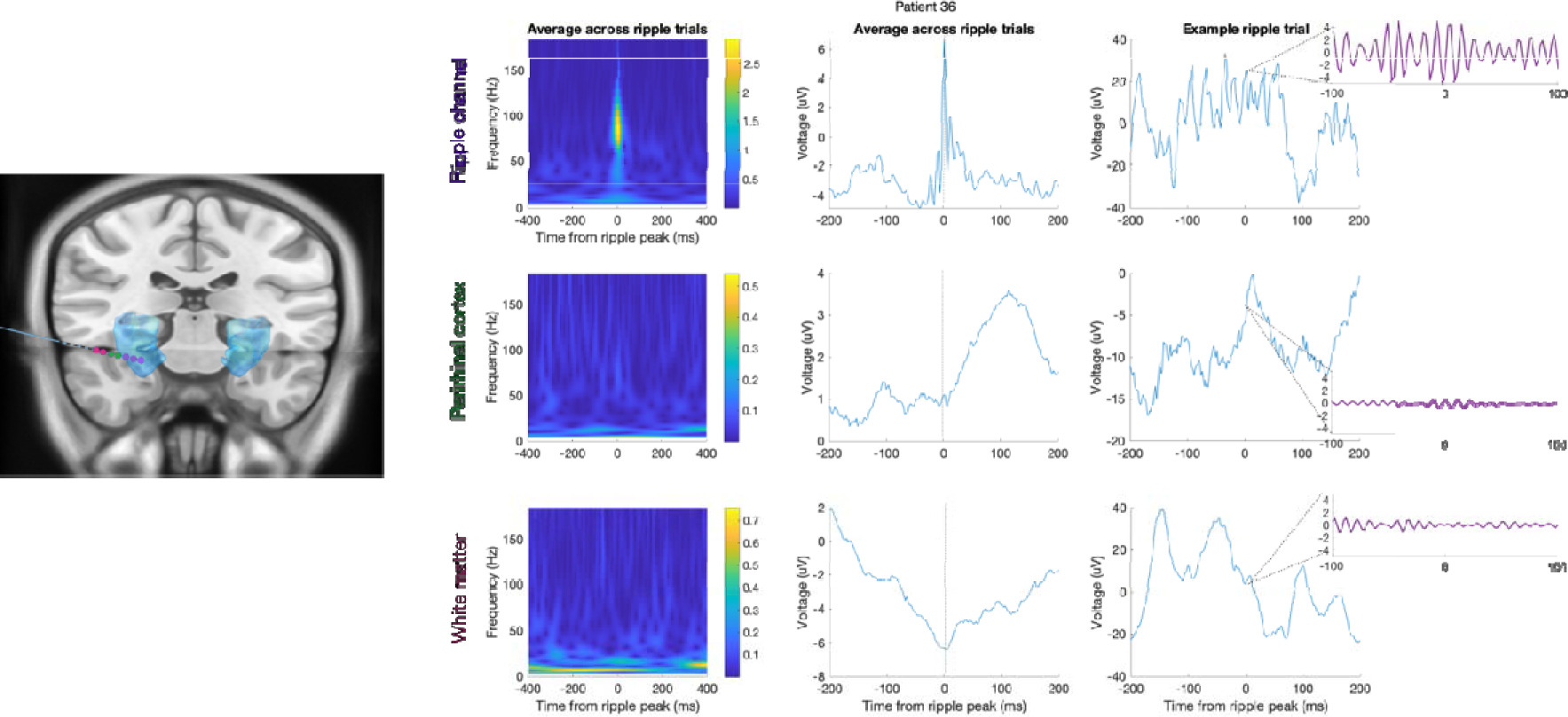
Averaged, and example single trial, ripple traces from hippocampal and adjacent contacts for Patient 36. *Left*. Electrode localisation shown on the smoothed hippocampus mask from the Automated Anatomical Labelling atlas for visualization; blue) illustrates the contacts (coloured circles) from which data are presented in the panels on the right. Purple circles: 3 hippocampal contacts i.e., two bipolar channels. Green circles: the two contacts adjacent to the hippocampus are in perirhinal cortex. Red circles: the two subsequent contacts are in white matter (medial contact)/CSF (lateral contact). *Right*. Top, bottom and middle rows: ripple-locked activity from the hippocampal (Ripple) channels (purple circles), and adjacent bipolar channels “Perirhinal cortex” (corresponding to green circles) and “White matter” (corresponding to red circles). The left and centre columns show the average peri-ripple wavelet spectrogram across all ripple trials (n = 381), and average raw field potential centred on ripple peak, respectively. The right column shows an example ripple trial (raw field potential centred on ripple peak) with an inset showing the ripple band activity (band pass filtered at 80-120Hz).

**Figure S22.**
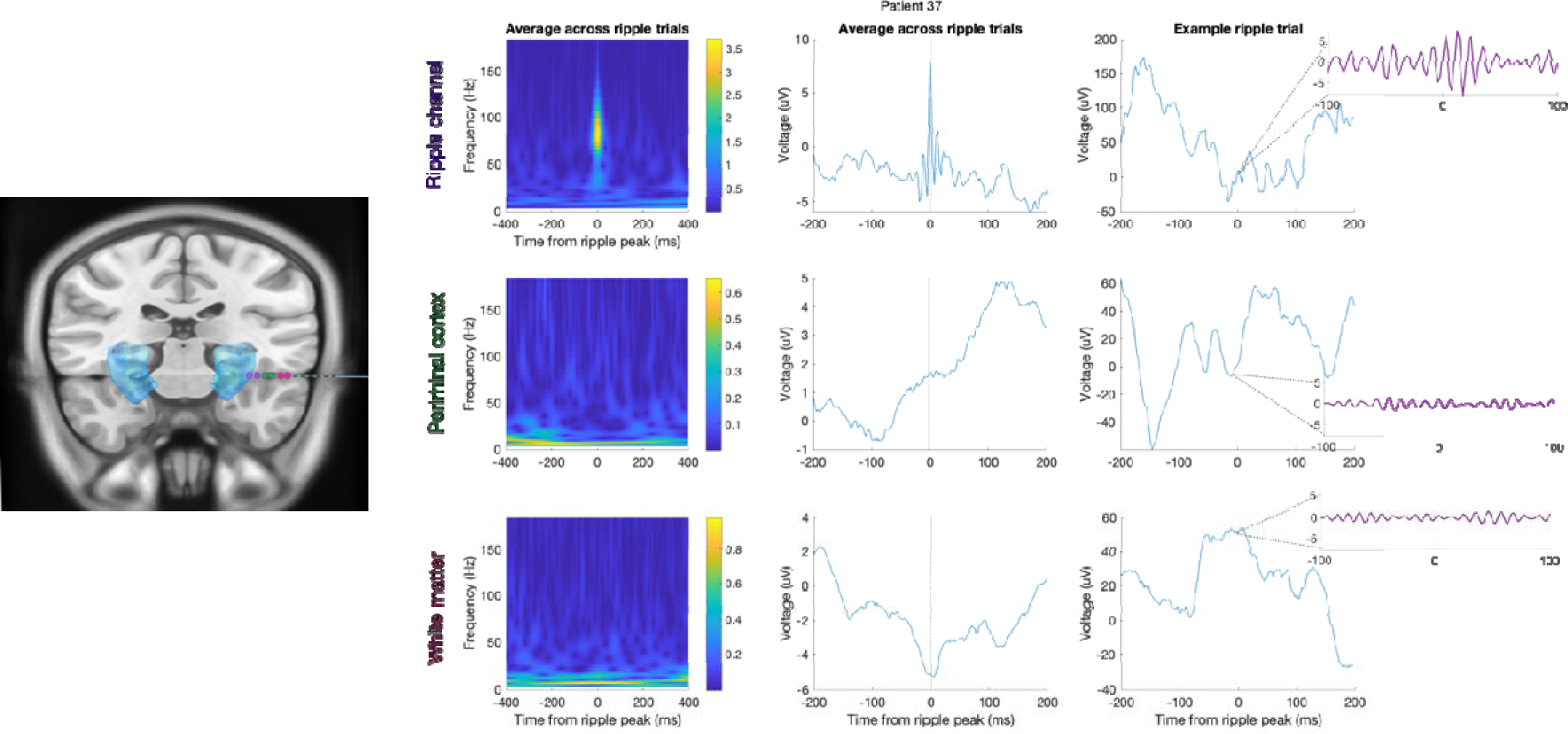
Averaged, and example single trial, ripple traces from hippocampal and adjacent contacts for Patient 37. *Left*. Electrode localisation shown on the smoothed hippocampus mask from the Automated Anatomical Labelling atlas for visualization; blue) illustrates the contacts (coloured circles) from which data are presented in the panels on the right. Purple circles: 2 hippocampal contacts i.e., one bipolar channel. Green circles: the two contacts adjacent to the hippocampus are in perirhinal cortex (medial contact)/white matter (lateral contact). Red circles: the two subsequent contacts are in white matter (medial contact)/inferior temporal sulcus (lateral contact). *Right*. Top, bottom and middle rows: ripple-locked activity from the hippocampal (Ripple) channel (purple circles), and adjacent bipolar channels “Perirhinal cortex” (corresponding to green circles) and “White matter” (corresponding to red circles). The left and centre columns show the average peri-ripple wavelet spectrogram across all ripple trials (n = 80), and average raw field potential centred on ripple peak, respectively. The right column shows an example ripple trial (raw field potential centred on ripple peak) with an inset showing the ripple band activity (band pass filtered at 80-120Hz).

**Figure S23.**
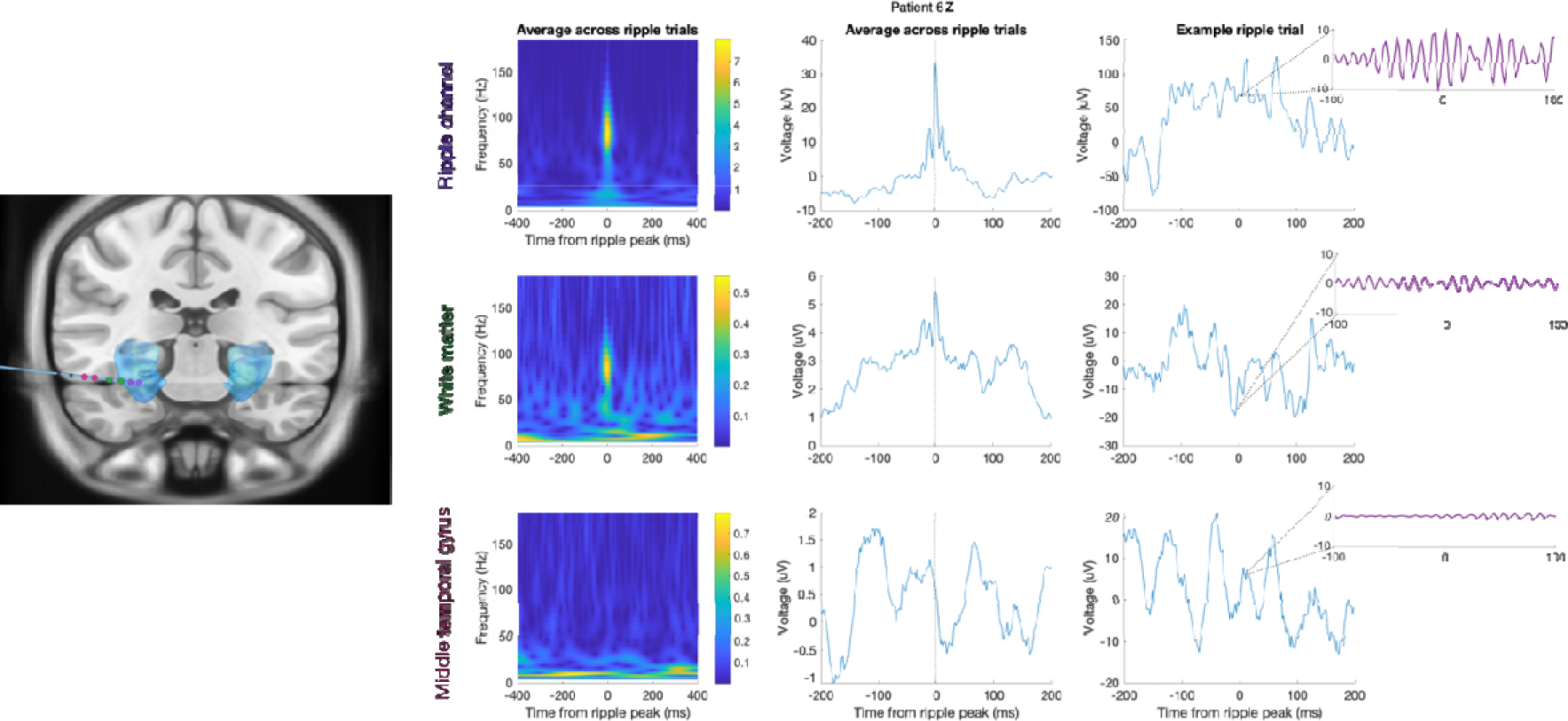
Averaged, and example single trial, ripple traces from hippocampal and adjacent contacts for Patient 6z. *Left*. Electrode localisation shown on the smoothed hippocampus mask from the Automated Anatomical Labelling atlas for visualization; blue) illustrates the contacts (coloured circles) from which data are presented in the panels on the right. Purple circles: 2 hippocampal contacts i.e., one bipolar channel. Green circles: the two contacts adjacent to the hippocampus are in white matter. Red circles: the two subsequent contacts are in middle temporal gyrus. *Right*. Top, bottom and middle rows: ripple-locked activity from the hippocampal (Ripple) channel (purple circles), and adjacent bipolar channels “White matter” (corresponding to green circles) and “Middle temporal gyrus” (corresponding to red circles). The left and centre columns show the average peri-ripple wavelet spectrogram across all ripple trials (n = 253), and average raw field potential centred on ripple peak, respectively. The right column shows an example ripple trial (raw field potential centred on ripple peak) with an inset showing the ripple band activity (band pass filtered at 80-120Hz).

**Figure S24.**
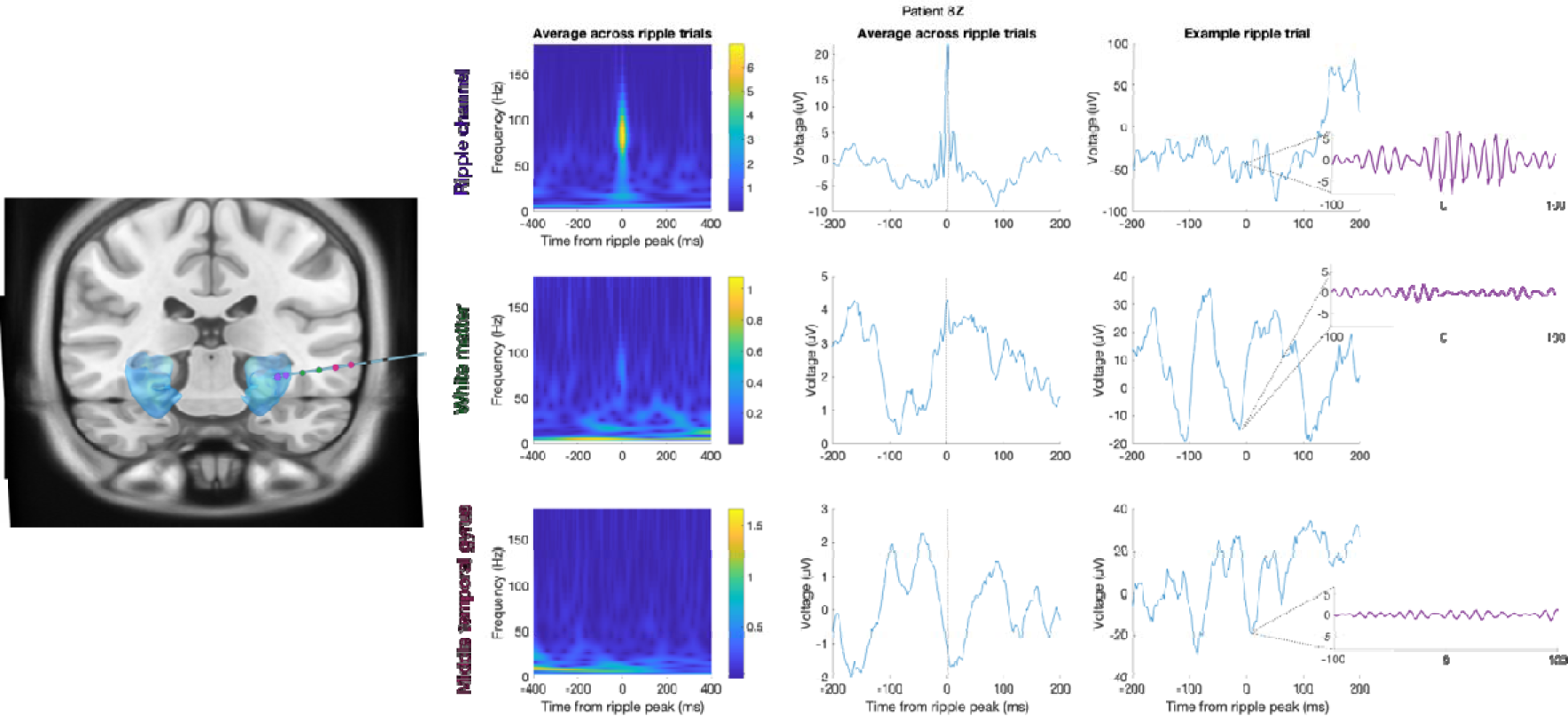
Averaged, and example single trial, ripple traces from hippocampal and adjacent contacts for Patient 8z. *Left*. Electrode localisation shown on the smoothed hippocampus mask from the Automated Anatomical Labelling atlas for visualization; blue) illustrates the contacts (coloured circles) from which data are presented in the panels on the right. Purple circles: 2 hippocampal contacts i.e., one bipolar channel. Green circles: the two contacts adjacent to the hippocampus are in white matter. Red circles: the two subsequent contacts are in middle temporal gyrus (medial contact)/white matter (lateral contact). *Right*. Top, bottom and middle rows: ripple-locked activity from the hippocampal (Ripple) channel (purple circles), and adjacent bipolar channels “White matter” (corresponding to green circles) and “Middle temporal gyrus” (corresponding to red circles). The left and centre columns show the average peri-ripple wavelet spectrogram across all ripple trials (n = 92), and average raw field potential centred on ripple peak, respectively. The right column shows an example ripple trial (raw field potential centred on ripple peak) with an inset showing the ripple band activity (band pass filtered at 80-120Hz).

**Figure S25.**
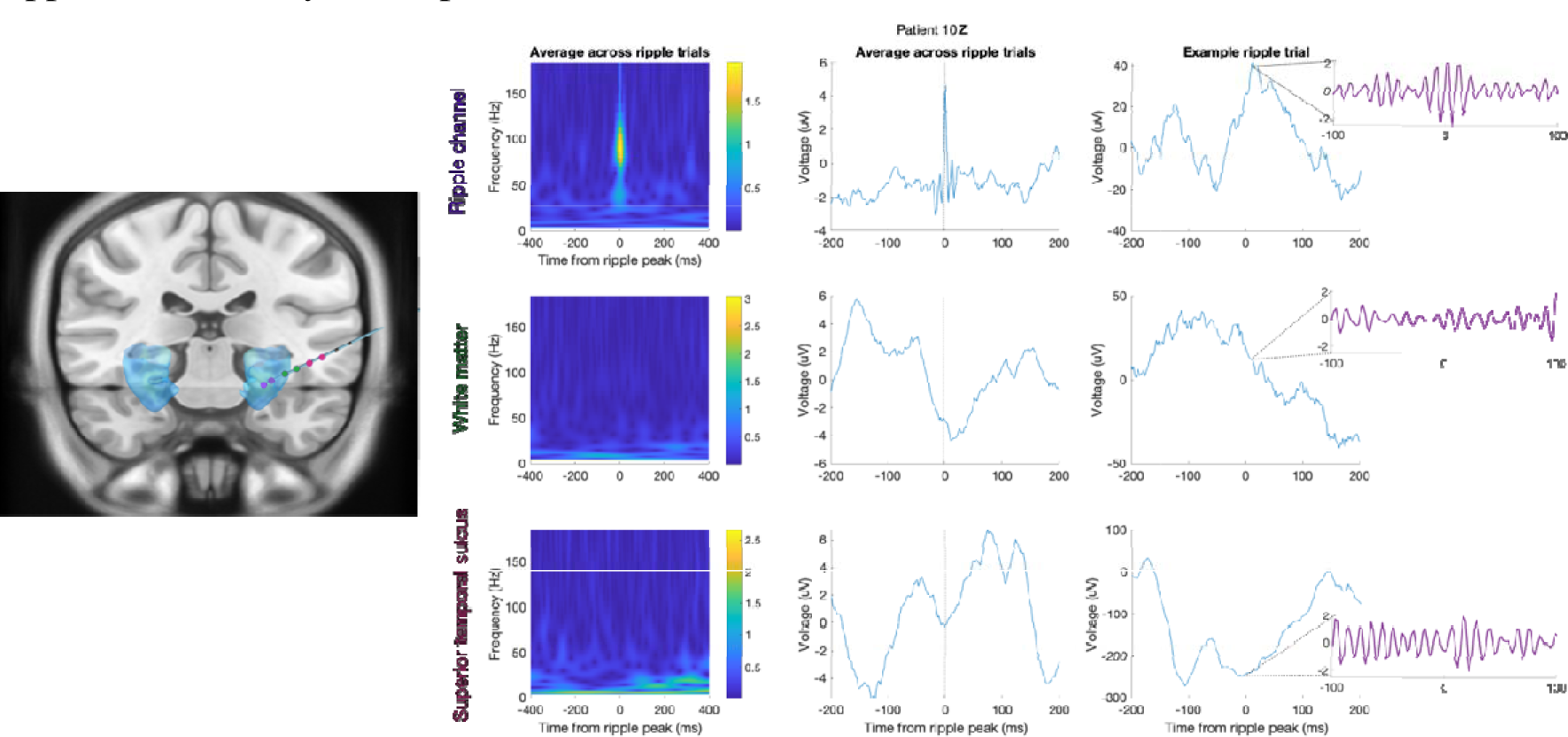
Averaged, and example single trial, ripple traces from hippocampal and adjacent contacts for Patient 10z. *Left*. Electrode localisation shown on the smoothed hippocampus mask from the Automated Anatomical Labelling atlas for visualization; blue) illustrates the contacts (coloured circles) from which data are presented in the panels on the right. Purple circles: 2 hippocampal contacts i.e., one bipolar channel. Green circles: the two contacts adjacent to the hippocampus are in white matter. Red circles: the two subsequent contacts are in superior temporal sulcus. *Right*. Top, bottom and middle rows: ripple-locked activity from the hippocampal (Ripple) channel (purple circles), and adjacent bipolar channels “White matter” (corresponding to green circles) and “Superior temporal sulcus” (corresponding to red circles). The left and centre columns show the average peri-ripple wavelet spectrogram across all ripple trials (n = 120), and average raw field potential centred on ripple peak, respectively. The right column shows an example ripple trial (raw field potential centred on ripple peak) with an inset showing the ripple band activity (band pass filtered at 80-120Hz).

**Figure S26.**
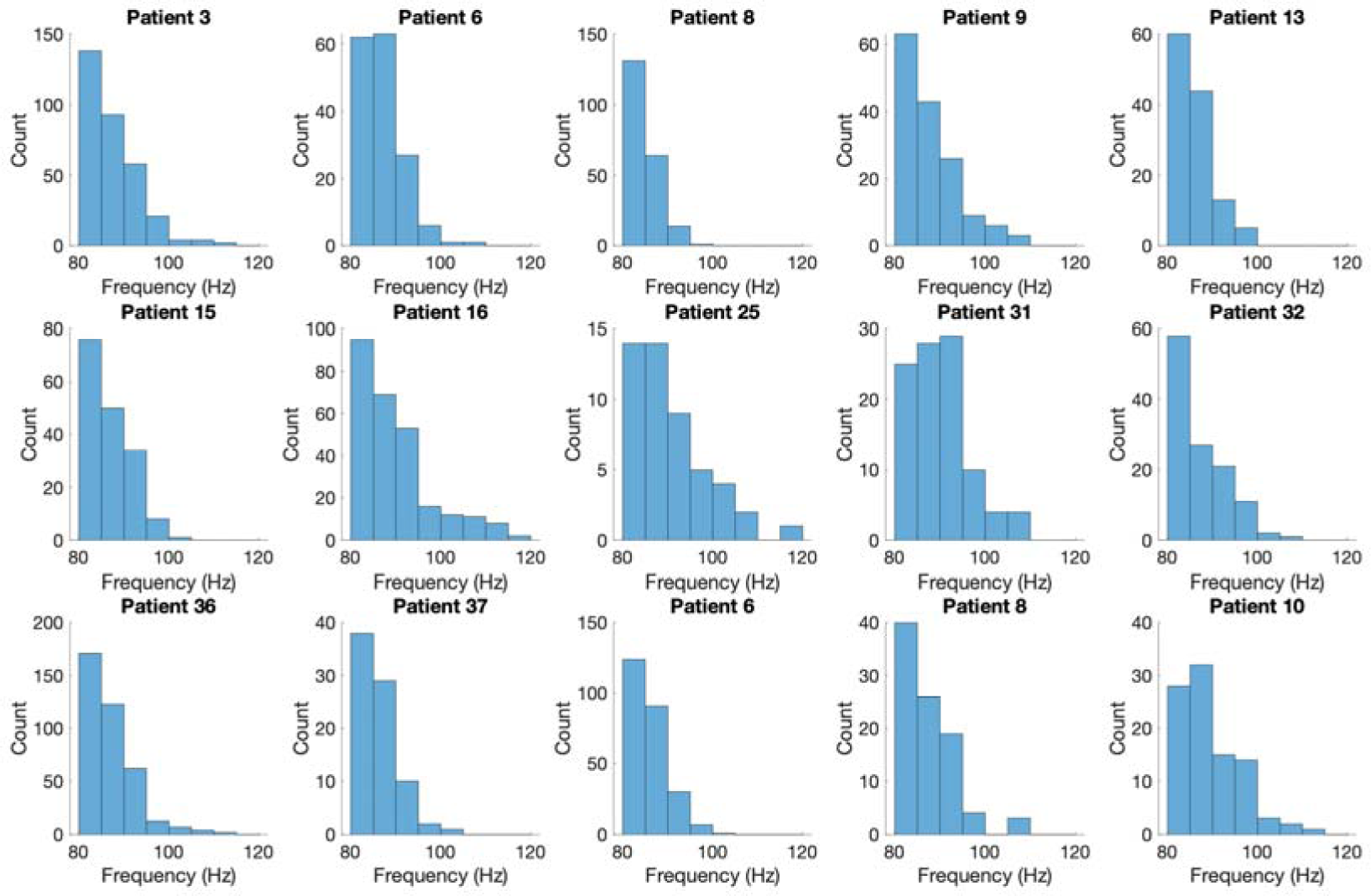
Ripple peak frequency per patient. Histograms showing the distribution of each patient’s ripples peak frequency. The majority of ripples occur at a frequency between 80 and 90 Hz (average ripple frequency over patients 87.5 Hz standard deviation 5.9).

**Figure S27.**
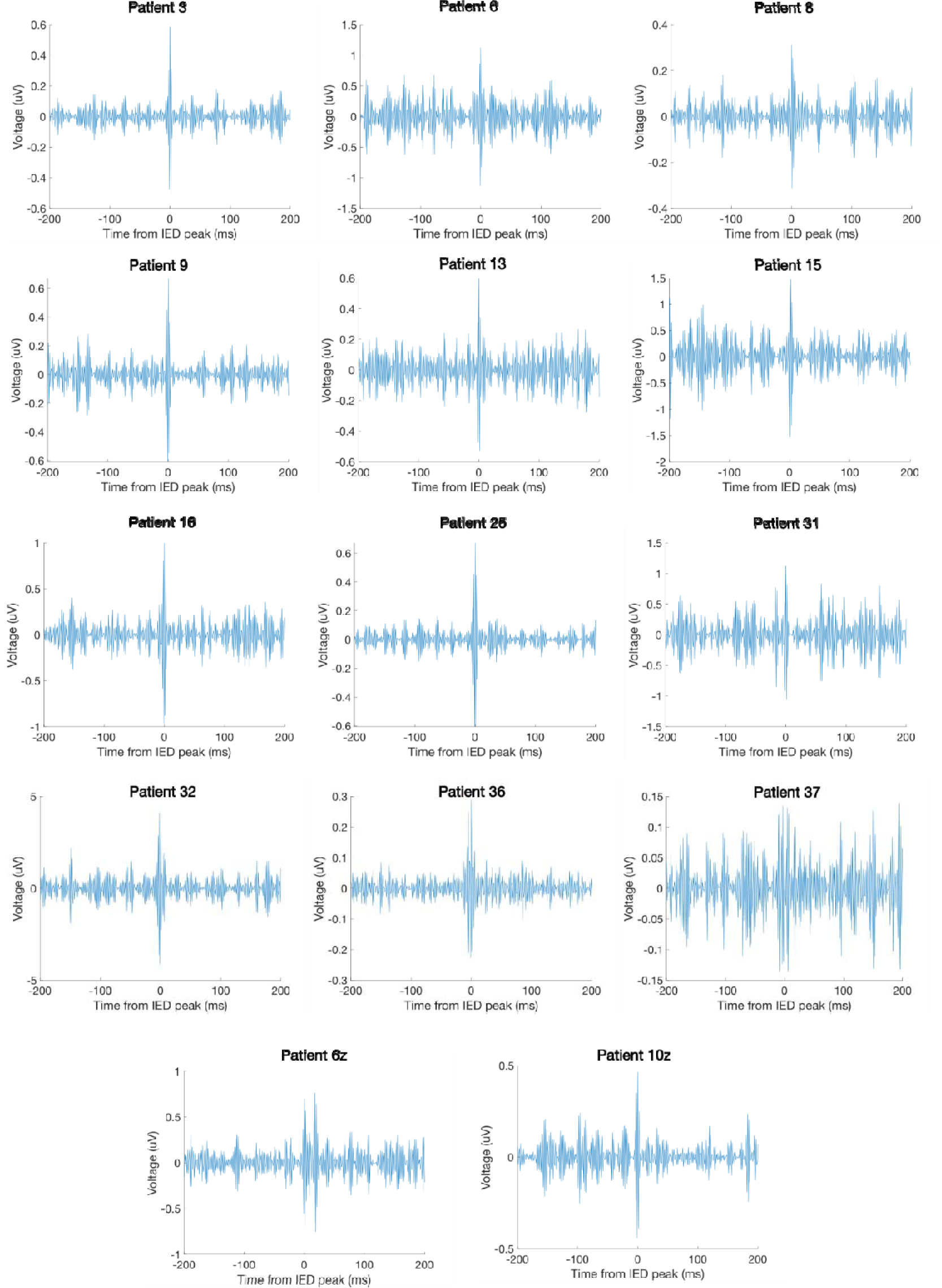
Mean IEDs per patient. High-pass filtered LFP (200Hz) across all detected IEDs, per patient.

**Figure S28.**
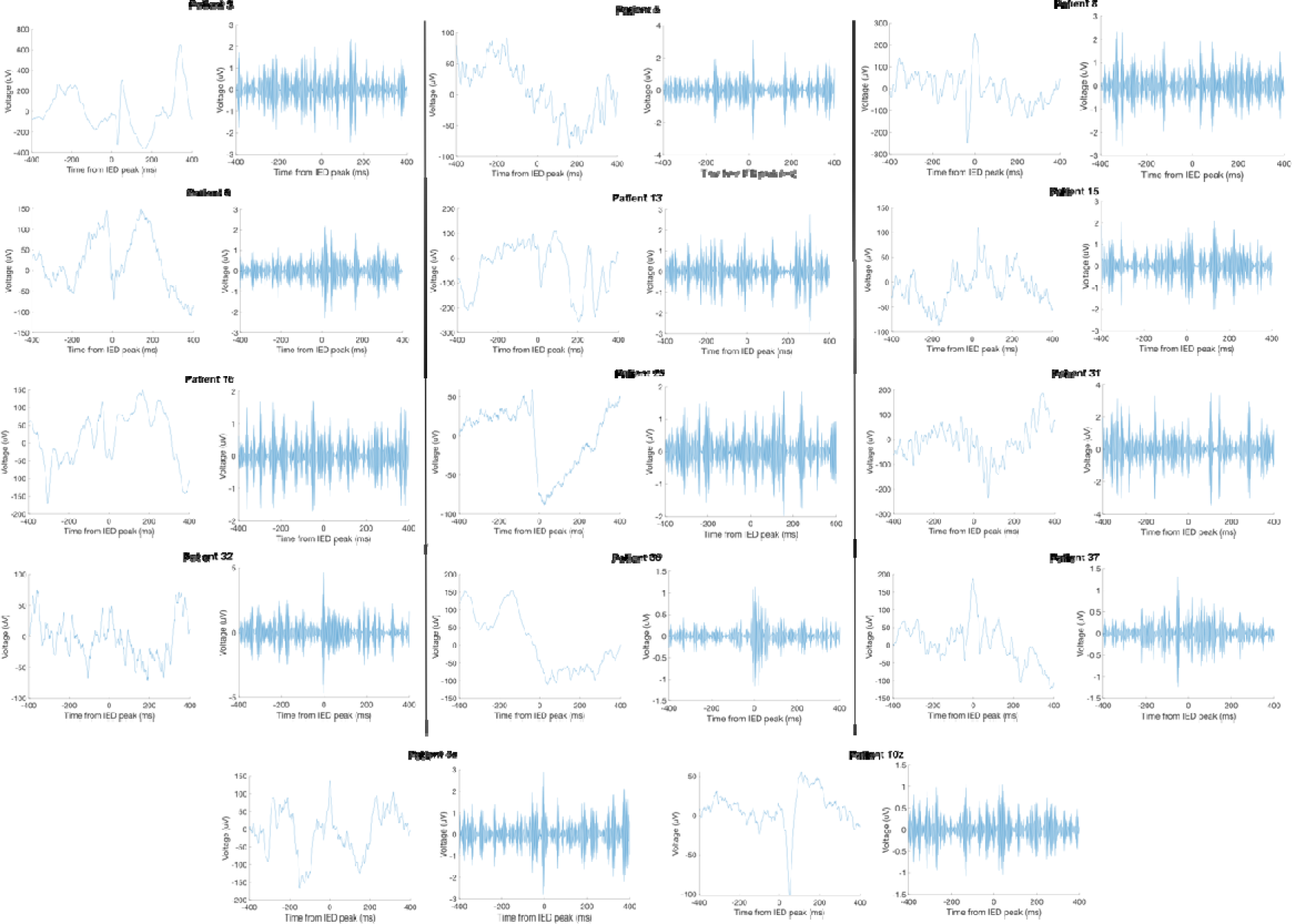
Example IEDs per patient. Raw and high-pass filtered LFP (200Hz) for an example IED, per patient.

## References

1. Gregory, R. L. Perceptions as hypotheses. Philosophical Transactions of the Royal Society of London. B, Biological Sciences 290, 181–197 (1980).

2. Friston, K. J. & Kiebel, S. J. Predictive coding under the free-energy principle. Philosophical Transactions of the Royal Society B: Biological Sciences 364, 1211– 1221 (2009).

3. Clark, A. Whatever next? Predictive brains, situated agents, and the future of cognitive science. Behavioral and Brain Sciences 36, 181–204 (2013).

4. Friston, K. J. A theory of cortical responses. Philosophical Transactions of the Royal Society B: Biological Sciences 360, 815–836 (2005).

5. Rao, R. P. N. & Ballard, D. H. Predictive coding in the visual cortex: a functional interpretation of some extra-classical receptive-field effects. Nat Neurosci 2, 79–87 (1999).

6. Mumford, D. On the computational architecture of the neocortex - I. The role of the thalamo-cortical loop. Biol Cybern 65, 135–145 (1991).

7. Yon, D. & Frith, C. D. Precision and the Bayesian brain. Current Biology 31, R1026– R1032 (2021).

8. Rao, R. P. N. An optimal estimation approach to visual perception and learning. Vision Res 39, 1963–1989 (1999).

9. Clark, A. The many faces of precision (Replies to commentaries on “Whatever next? Neural prediction, situated agents, and the future of cognitive science”). Front Psychol 4, (2013).

10. Moran, R. J. et al. Free Energy, Precision and Learning: The Role of Cholinergic Neuromodulation. Journal of Neuroscience 33, 8227–8236 (2013).

11. FitzGerald, T. H. B., Moran, R. J., Friston, K. J. & Dolan, R. J. Precision and neuronal dynamics in the human posterior parietal cortex during evidence accumulation. Neuroimage 107, 219–228 (2015).

12. Hesselmann, G., Sadaghiani, S., Friston, K. J. & Kleinschmidt, A. Predictive Coding or Evidence Accumulation? False Inference and Neuronal Fluctuations. PLoS One 5, e9926 (2010).

13. Hindy, N. C., Ng, F. Y. & Turk-Browne, N. B. Linking pattern completion in the hippocampus to predictive coding in visual cortex. Nat Neurosci 1–7 (2016) doi:10.1038/nn.4284.

14. Dimakopoulos, V., Mégevand, P., Stieglitz, L. H., Imbach, L. & Sarnthein, J. Information flows from hippocampus to auditory cortex during replay of verbal working memory items. Elife 11, (2022).

15. Barron, H. C., Auksztulewicz, R. & Friston, K. J. Prediction and memory: A predictive coding account. Prog Neurobiol 192, (2020).

16. Bornstein, A. M. & Daw, N. D. Cortical and Hippocampal Correlates of Deliberation during Model-Based Decisions for Rewards in Humans. PLoS Comput Biol 9, (2013).

17. Fuhrer, J. et al. Direct brain recordings reveal continuous. 1–14 (2021).

18. Schapiro, A. C., Turk-Browne, N. B., Norman, K. A. & Botvinick, M. M. Statistical learning of temporal community structure in the hippocampus. Hippocampus 26, 3–8 (2016).

19. Strange, B. A., Duggins, A., Penny, W., Dolan, R. J. & Friston, K. J. Information theory, novelty and hippocampal responses: Unpredicted or unpredictable? Neural Networks 18, 225–230 (2005).

20. Barron, H. C., Dolan, R. J. & Behrens, T. E. J. Online evaluation of novel choices by simultaneous representation of multiple memories. Nat Neurosci 16, 1492–1498 (2013).

21. Kaplan, R. et al. The Neural Representation of Prospective Choice during Spatial Planning and Decisions. PLoS Biol 15, 1–26 (2017).

22. Axmacher, N. et al. Intracranial EEG Correlates of Expectancy and Memory Formation in the Human Hippocampus and Nucleus Accumbens. Neuron 65, 541–549 (2010).

23. Lisman, J. E. & Redish, A. D. Prediction, sequences and the hippocampus. Philosophical Transactions of the Royal Society B 364, 1193–1201 (2009).

24. Hassabis, D. & Maguire, E. A. Deconstructing episodic memory with construction. Trends Cogn Sci 11, 299–306 (2007).

25. Kay, K. & Frank, L. M. Three brain states in the hippocampus and cortex. Hippocampus 29, 184–238 (2019).

26. Frank, D. & Kafkas, A. Expectation-driven novelty effects in episodic memory. Neurobiol Learn Mem 183, 107466 (2021).

27. Dayan, P. Improving Generalization for Temporal Difference Learning: The Successor Representation. Neural Comput 5, 613–624 (1993).

28. Ekman, M., Kusch, S. & de Lange, F. P. Successor-like representation guides the prediction of future events in human visual cortex and hippocampus. bioRxiv 124-(2022) 10.1101/2022.03.23.485480.

29. Moser, E. I., Kropff, E. & Moser, M. B. Place cells, grid cells, and the brain’s spatial representation system. Annu Rev Neurosci 31, 69–89 (2008).

30. Bell, A. H., Summerfield, C., Morin, E. L., Malecek, N. J. & Ungerleider, L. G. Encoding of Stimulus Probability in Macaque Inferior Temporal Cortex. Current Biology 26, 2280–2290 (2016).

31. Arnal, L. H. & Giraud, A. L. Cortical oscillations and sensory predictions. Trends Cogn Sci 16, 390–398 (2012).

32. Bastos, A. M. et al. Canonical Microcircuits for Predictive Coding. Neuron 76, 695– 711 (2012).

33. Chao, Z. C., Takaura, K., Wang, L., Fujii, N. & Dehaene, S. Large-Scale Cortical Networks for Hierarchical Prediction and Prediction Error in the Primate Brain. Neuron 100, 1252–1266.e3 (2018).

34. Bastos, A. M. et al. Visual areas exert feedforward and feedback influences through distinct frequency channels. Neuron 85, 390–401 (2015).

35. Bastos, A. M., Lundqvist, M., Waite, A. S., Kopell, N. & Miller, E. K. Layer and rhythm specificity for predictive routing. Proc Natl Acad Sci U S A 117, 31459–31469 (2020).

36. Joo, H. R. & Frank, L. M. The hippocampal sharp wave–ripple in memory retrieval for immediate use and consolidation. Nat Rev Neurosci 19, 744–757 (2018).

37. Liu, A. A. et al. A consensus statement on detection of hip-pocampal sharp wave ripples and differ-entiation from other fast oscillations. (2022) doi:10.1038/s41467-022-33536-x.

38. Swanson, R. A., Levenstein, D., McClain, K., Tingley, D. & Buzsáki, G. Variable specificity of memory trace reactivation during hippocampal sharp wave ripples. Curr Opin Behav Sci 32, 126–135 (2020).

39. Buzsáki, G. Hippocampal sharp wave-ripple: A cognitive biomarker for episodic memory and planning. Hippocampus 25, 1073–1188 (2015).

40. Joo, H. R. & Frank, L. M. The hippocampal sharp wave–ripple in memory retrieval for immediate use and consolidation. Nat Rev Neurosci 19, 744–757 (2018).

41. Diba, K. & Buzsáki, G. Forward and reverse hippocampal place-cell sequences during ripples. Nat Neurosci 10, 1241–1242 (2007).

42. Pfeiffer, B. E. & Foster, D. J. Hippocampal place-cell sequences depict future paths to remembered goals. Nature 497, 74–79 (2013).

43. Gupta, A. S., van der Meer, M. A. A., Touretzky, D. S. & Redish, A. D. Hippocampal Replay Is Not a Simple Function of Experience. Neuron 65, 695–705 (2010).

44. Dragoi, G. & Tonegawa, S. Preplay of future place cell sequences by hippocampal cellular assemblies. Nature 469, 397–401 (2011).

45. Gupta, A. S., van der Meer, M. A. A., Touretzky, D. S. & Redish, A. D. Hippocampal Replay Is Not a Simple Function of Experience. Neuron 65, 695–705 (2010).

46. Pfeiffer, B. E. & Foster, D. J. Hippocampal place-cell sequences depict future paths to remembered goals. Nature 497, 74–79 (2013).

47. Logothetis, N. K. et al. Hippocampal-cortical interaction during periods of subcortical silence. Nature 491, 547–553 (2012).

48. Nitzan, N., Swanson, R., Schmitz, D. & Buzsáki, G. Brain-wide interactions during hippocampal sharp wave ripples. Proc Natl Acad Sci U S A 119, e2200931119 (2022).

49. Kaplan, R. et al. Hippocampal Sharp-Wave Ripples Influence Selective Activation of the Default Mode Network. Current Biology 26, 686–691 (2016).

50. Silva, D., Feng, T. & Foster, D. J. Trajectory events across hippocampal place cells require previous experience. Nat Neurosci 18, 1772–1779 (2015).

51. Gillespie, A. K. et al. Hippocampal replay reflects specific past experiences rather than a plan for subsequent choice. Neuron 109, 3149–3163.e6 (2021).

52. Findlay, G., Tononi, G. & Cirelli, C. The evolving view of replay and its functions in wake and sleep. SLEEP Advances 1, 1–14 (2020).

53. Norman, Y. et al. Hippocampal sharp-wave ripples linked to visual episodic recollection in humans. Science (1979) 365, (2019).

54. Vaz, A. P., Inati, S. K., Brunel, N. & Zaghloul, K. A. Coupled ripple oscillations between the medial temporal lobe and neocortex retrieve human memory. Science (1979) 363, 975–978 (2019).

55. Axmacher, N., Elger, C. E. & Fell, J. Ripples in the medial temporal lobe are relevant for human memory consolidation. Brain 131, 1806–1817 (2008).

56. Hick, W. E. On the Rate of Gain of Information. Quarterly Journal of Experimental Psychology 4, 11–26 (1952).

57. Fernández-Ruiz, A. et al. Long-duration hippocampal sharp wave ripples improve memory. Science (1979) 364, 1082–1086 (2019).

58. Haarsma, J. et al. Precision weighting of cortical unsigned prediction error signals benefits learning, is mediated by dopamine, and is impaired in psychosis. Mol Psychiatry 26, 5320–5333 (2021).

59. Limanowski, J. Precision control for a flexible body representation. Neurosci Biobehav Rev 134, 104401 (2022).

60. Kok, P., Jehee, J. F. M. & de Lange, F. P. Less Is More: Expectation Sharpens Representations in the Primary Visual Cortex. Neuron 75, 265–270 (2012).

61. Friston, K. Hierarchical Models in the Brain. PLoS Comput Biol 4, e1000211 (2008).

62. Friston, K. J. A theory of cortical responses. Philosophical Transactions of the Royal Society B: Biological Sciences 360, 815–836 (2005).

63. Diba, K. & Buzsáki, G. Forward and reverse hippocampal place-cell sequences during ripples. Nat Neurosci 10, 1241–1242 (2007).

64. Buzsáki, G. Hippocampal sharp wave-ripple: A cognitive biomarker for episodic memory and planning. Hippocampus 25, 1073–1188 (2015).

65. Diba, K. Hippocampal sharp-wave ripples in cognitive map maintenance versus episodic simulation. Neuron 109, 3071–3074 (2021).

66. Silva, D., Feng, T. & Foster, D. J. Trajectory events across hippocampal place cells require previous experience. Nat Neurosci 18, 1772–1779 (2015).

67. Tong, A. P. S., Vaz, A. P., Wittig, J. H., Inati, S. K. & Zaghloul, K. A. Ripples reflect a spectrum of synchronous spiking activity in human anterior temporal lobe. Elife 10, 1–25 (2021).

68. de Lange, F. P., Heilbron, M. & Kok, P. How Do Expectations Shape Perception? Trends Cogn Sci 22, 764–779 (2018).

69. Pezzulo, G., Parr, T. & Friston, K. The evolution of brain architectures for predictive coding and active inference. Philosophical Transactions of the Royal Society B: Biological Sciences 377, (2022).

70. Rao, R. P. N. & Ballard, D. H. Predictive coding in the visual cortex: a functional interpretation of some extra-classical receptive-field effects. Nat Neurosci 2, 79–87 (1999).

71. Ramirez-Villegas, J. F., Logothetis, N. K. & Besserve, M. Diversity of sharp-wave-ripple LFP signatures reveals differentiated brain-wide dynamical events. Proc Natl Acad Sci U S A 112, E6379–E6387 (2015).

72. Méndez-Bértolo, C. et al. A fast pathway for fear in human amygdala. Nat Neurosci 19, 1041–1049 (2016).

73. Avants, B. B., Tustison, N. J. & Johnson, H. Advanced normalization tools (ants). Insight j 2, (2009).

74. Treu, S. et al. Deep brain stimulation: Imaging on a group level. Neuroimage 219, 117018 (2020).

75. Oostenveld, R., Fries, P., Maris, E. & Schoffelen, J.-M. FieldTrip: Open Source Software for Advanced Analysis of MEG, EEG, and Invasive Electrophysiological Data. Comput Intell Neurosci 2011, 1–9 (2011).

76. Hamamé, C. M. et al. Functional selectivity in the human occipitotemporal cortex during natural vision: Evidence from combined intracranial EEG and eye-tracking. Neuroimage 95, 276–286 (2014).

77. Shirhatti, V., Borthakur, A. & Ray, S. Effect of Reference Scheme on Power and Phase of the Local Field Potential. Neural Comput 28, 882–913 (2016).

78. Trongnetrpunya, A. et al. Assessing Granger Causality in Electrophysiological Data: Removing the Adverse Effects of Common Signals via Bipolar Derivations. Front Syst Neurosci 9, (2016).

79. Patel, J., Schomburg, E. W., Berényi, A., Fujisawa, S. & Buzsáki, G. Local generation and propagation of ripples along the septotemporal axis of the hippocampus. Journal of Neuroscience 33, 17029–17041 (2013).

80. Bates, D., Mächler, M., Bolker, B. & Walker, S. Fitting Linear Mixed-Effects Models Using lme4. J Stat Softw 67, (2015).

